# Large-scale Analysis of 2,152 dataset reveals key features of B cell biology and the antibody repertoire

**DOI:** 10.1101/814590

**Authors:** Xiujia Yang, Minhui Wang, Dianchun Shi, Yanfang Zhang, Huikun Zeng, Yan Zhu, Chunhong Lan, Jiaqi Wu, Yang Deng, Shixin Guo, Lijun Xu, Cuiyu Ma, Yanxia Zhang, Rongrong Wu, Jinxia Ou, Chu-jun Liu, Changqing Chang, Wei Yang, Huijie Zhang, Jun Chen, Lijie Qin, Hongwei Zhou, Jin-Xin Bei, Lai Wei, Guangwen Cao, Xueqing Yu, Zhenhai Zhang

**Affiliations:** State Key Laboratory of Organ Failure Research, National Clinical Research Center for Kidney Disease, Division of Nephrology, Nanfang Hospital, Southern Medical University, Guangzhou, 510515, China; Department of Bioinformatics, School of Basic Medical Sciences, Southern Medical University, Guangzhou 510515, China; Department of Geriatrics, Guangzhou First People’s Hospital, School of Medicine, South China University of Technology, Guangzhou 510030, China; Department of Epidemiology, Second Military Medical University, 800 Xiangyin Rd., Shanghai 200433, China; State Key Laboratory of Ophthalmology, Zhongshan Ophthalmic Center, Sun Yat-Sen University, Guangzhou 510060, China; Department of Emergency, Henan Provincial People’s Hospital, Zhengzhou 450003, China; Microbiome Medicine Center, Division of Laboratory Medicine, Zhujiang Hospital, Southern Medical University, Guangzhou 510282, China; Sun Yat-Sen University Cancer Center, State Key Laboratory of Oncology in South China, Collaborative Innovation Center for Cancer Medicine, Sun Yat-Sen University, Guangzhou 510030, China; Integrate Microbiology Research Center, South China Agricultural University, Guangzhou, 510642, China; Department of Pathology, School of Basic Medical Sciences, Southern Medical University, Guangzhou, 510515, China; Department of Endocrinology and Metabolism, Nanfang Hospital, Southern Medical University, Guangzhou 510515, China; MOE Laboratory of Biosystems Homeostasis & Protection and Innovation Center for Cell Signaling Network, College of Life Sciences, Zhejiang University, Hangzhou 310058, China; Center for precision medicine, Guangzhou First People’s Hospital, School of Medicine, South China University of Technology, Guangzhou 510030, China; Key Laboratory of Mental Health of the Ministry of Education, Guangdong-Hong Kong-Macao Greater Bay Area Center for Brain Science and Brain-Inspired Intelligence, Southern Medical University, Guangzhou 510515, China

**Author notes:** These authors contributed equally to this work (Xiujia Yang, Minhui Wang, Dianchun Shi, Yanfang Zhang, Huikun Zeng, Yan Zhu, Chunhong Lan). To whom correspondence should be addressed: Zhenhai Zhang,; Xueqing Yu,; Guangwen Cao.

**Keywords:** B-cell biology, antibody repertoire, large-scale analysis, high-throughput sequencing, Ig-seq

## Abstract

Antibody repertoire sequencing (Ig-seq) has been widely used in studying humoral responses, with promising results. However, the promise of Ig-seq has not yet been fully realized, and key features of the antibody repertoire remain elusive or controversial. To clarify these key features, we analyzed 2,152 high-quality heavy chain antibody repertoires, representing 582 donors and a total of 360 million clones. Our study revealed that individuals exhibit very similar gene usage patterns for germline V, D, and J genes and that 53 core V genes contribute to more than 99% of the heavy chain repertoire. We further found that genetic background is sufficient but not necessary to determine usage of V, D, and J genes. Although gene usage pattern is not affected by age, we observed a significant sex preference for 24 V genes, 9 D genes and 5 J genes, but found no positional bias for V-D and D-J recombination. In addition, we found that the number of observed clones that were shared between any two repertoires followed a linear model and noted that the mutability of hot/cold spots and single nucleotides within antibody genes suggested a strand-specific somatic hypermutation mechanism. This population-level analysis resolves some critical characteristics of the antibody repertoire and thus may serve as a reference for research aiming to unravel B cell-related biology or diseases. The metrics revealed here will be of significant value to the large cadre of scientists who study the antibody repertoire.

## Introduction

The antibody repertoire is defined as the entire collection of B-cell receptors and antibodies that grant protection against a plethora of pathogens. A deeper understanding of the antibody repertoire under normal physiological conditions and in pathogenic conditions may shed light on functional immune responses and reveal the full scope of their protective and pathogenic functions. However, despite this great potential, collecting enough antibody molecules to capture the immense diversity of the antibody repertoire has been a critical challenge.

Using high-throughput sequencing technology, Weinstein et al. developed antibody repertoire sequencing (Ig-seq) (Weinstein et al., 2009), which allows researchers to capture millions or even billions of antibody variable regions within a single experiment. The vast amount of data acquired by Ig-seq enables a deeper and more thorough evaluation of the key features of the antibody repertoire, as well as its constituent antibody molecules, at the single-nucleotide level. In the past decade, Ig-seq has advanced the study of many important sub-fields of B-cell immunology, such as antibody discovery (Reddy et al., 2010; Zhu et al., 2013a), vaccination development (Jackson et al., 2014; Jiang et al., 2013; Joyce et al., 2016; Li et al., 2012), infection (Krebs et al., 2019; Parameswaran et al., 2013; Wu et al., 2015; Wu et al., 2011a), allergy (Hoh et al., 2016; Patil et al., 2015; Wu et al., 2014), autoimmune disease (Stern et al., 2014; Tipton et al., 2015; von Büdingen et al., 2012), and cancer immunology (Faham et al., 2012; Gawad et al., 2012; Kurtz et al., 2015). For example, using Ig-seq coupled with single-cell cloning technology, we and others identified thousands of HIV-1-neutralizing antibodies that bind to different epitopes and delineated their lineage-dependent maturation pathways (Bonsignori et al., 2016; Wu et al., 2015; Wu et al., 2011b; Zhu et al., 2013b). Studies of antibody repertoires after virus infection also led to the discovery of antibody convergence – a mechanism whereby identical or very similar antibody clonotypes are generated in different individuals facing the same selective pressure (Parameswaran et al., 2013). These results suggested that the antibody repertoire could be used to track an individual’s immune history as well as to monitor the immunological memory of a community.

The use of antibody repertoire in autoimmune diseases has provided important insight into both disease mechanisms and fundamental B cell biology. For example, Tipton et al. revealed that systemic lupus erythematosus (SLE) autoreactivity occurred during a polyclonal activation of IGHV4-34-dominant B cell clones via both germinal center-dependent and germinal center-independent mechanisms (Tipton et al., 2015). Büdingen et al. discovered that a pool of clonal related antibodies participates in a robust bidirectional exchange across the blood-brain barrier (von Büdingen et al., 2012). Analyzing the antibody repertoires of patients with the same disease, Stern et al. found that majority of the disease-related autoantibodies matured outside of the central nervous system and trafficked freely across tissue barriers (Stern et al., 2014).

Prior studies using Ig-Seq accumulated a wealth of antibody repertoire data. This under-explored population-level big data could potentially help us resolve the important yet unclear or controversial features of the B cell biology and the antibody repertoire. For instance, what are the germline V, D, and J gene usage patterns and how similar are they between individuals? What are the factors determining these patterns if they do exist? Is there preferential recombination between V-D and D-J genes? What are the rules that govern the somatic hypermutations (SHM)? What are the proportion of public clones between individuals and what functions do these clones exert?

With these unsolved or controversial questions in mind, we collected 2,152 high quality antibody heavy chain repertoires and performed thorough and in-depth analyses. These analyses revealed patterns of B cell biology as well as key features of the antibody repertoire, which will be of significant value to the large cadre of scientists in the field.

## Results

### Overview of datasets used in this study

The immense diversity of B cells is derived from two important biological processes: germline gene segment recombination, which introduces indels in complementarity-determining region 3, and activation-induced cytidine deaminase, which leads to somatic hypermutations in the antibody variable regions during affinity maturation. Thus, any mutation or indel in the variable region may be important for understanding and revealing B cell biology. It is thus essential to understand the sequencing errors that are intrinsic to Ig-seq approaches. For example, 454 sequencing often generates indels in the homopolymer region, and PCR amplification and high-throughput sequencing can also generate base errors and chimeras. These intrinsic errors would be easily mistaken as somatic hypermutation (SHM) generated during affinity maturations. We therefore only included samples that were sequenced by Illumina instruments. We also required the sequencing reads to cover a minimum of 500 bp, the primers to capture the full spectrum of antibodies generated by any V(D)J recombination, and a minimum number of productive reads (Materials and Methods). After filtering on these stringent criteria, we identified a total of 1,857 repertoires from 33 published studies and 295 repertoires from in-house sequencing for further analysis (Figure 1a).

**Figure 1.**
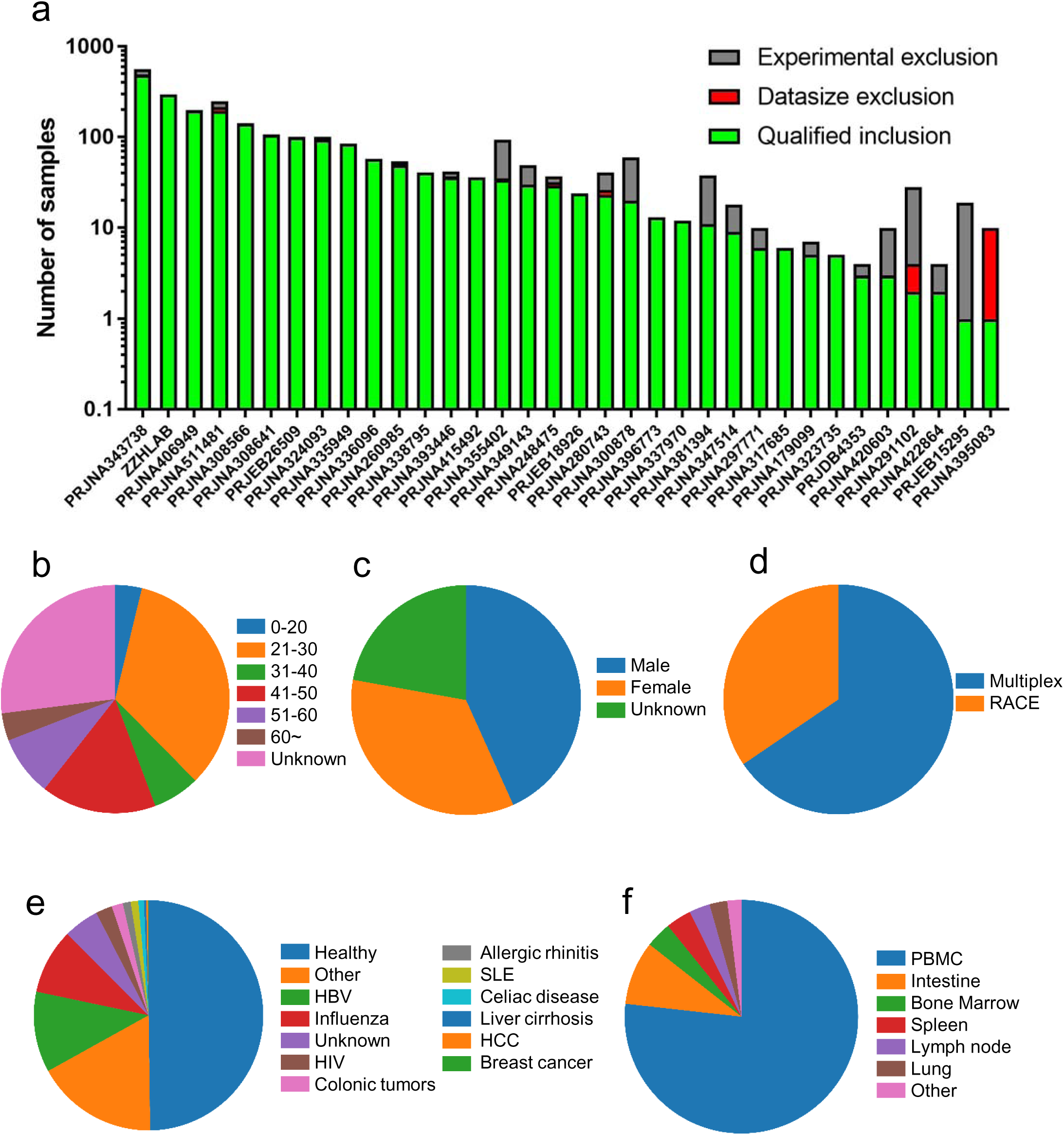
Overview of the enrolled datasets. (a) The number of samples in each enrolled project. The X axis shows NCBI SRA project IDs, and ZZHLAB indicates the antibody repertoires generated in our lab. The Y axis shows the log10 transformed number of samples. The numbers of samples excluded by data size and experimental design are shown in red and grey, respectively. **(b, c, d, e, f)** show sample distribution based on **(b)** age; **(c)** sex; **(d)** PCR amplification strategy; **(e)** classification (healthy or diseased); and **(f)** various tissue or blood.

The sample-associated metadata, including age, sex, physiological condition, tissue origin, and amplification method, are shown in Figures 1b-f. Age composition was more balanced for the sampled individuals than for the samples (Figure 1b and Figure S1), and the number of individuals for each age group is more than 30 (more than 80 samples for each age group). Slightly more than half of the samples were from females (Figure 1c). Sequencing libraries for all recruited samples were mainly amplified using multiplex PCR (Figure 1d). Donor conditions and the sources of the samples were classified into 13 and 6 directories respectively (Figures 1e and 1f). These repertoires covered a broad spectrum of diseases, such as autoimmune disease, cancer, virus infection, and more. The majority (76.8%) of samples were derived from peripheral blood mononuclear cells (PBMCs); there were also samples from bone marrow, intestine, lung, and spleen. Overall, we included a total of 7,378,354,271 raw reads in our analysis.

### The core V gene set determines the clear majority of antibodies

The variable usage of germline genes represents the first level of antibody repertoire diversity and is believed to affect immune function (Glanville et al., 2011). Naïve germline gene usage may be optimized for interactions with common antigens and may serve as a control to detect pathology-driven repertoire variation in the B cell memory compartment (Laserson et al., 2014). For these reasons, the gene usage pattern has been studied at a small scale and under different experimental settings using two different quantification methods: gene usage and gene expression. Gene usage quantifies genes at the level of individual clones, whereas gene expression quantifies the occurrence of genes with each read. Gene expression is sensitive to cell type composition, such as clonal expansion in response to an adaptive immune stimulus, and thus is less optimal for comparisons between samples with differences in source tissues, immune status, and donor health.

Library preparation technique also affects the quantification of genes. Two amplification strategies were used in the high-quality Ig-seq datasets: multiplex polymerase chain reaction (MPCR) and rapid amplification of cDNA ends (RACE). Previous studies showed that MPCR can introduce bias in library sequencing, even with an optimized primer set, while RACE introduces less bias (He et al., 2015; Liu et al., 2016; Robins, 2013). We compared quantitative metrics for 1,409 and 743 samples amplified by MPCR and RACE, respectively. D and J gene usage were less influenced by either RACE or MPCR. However, V gene usage was more consistent between RACE and MPCR (Figure S2a-f and Materials and Methods) due to the various primers on the 5’ ends. We therefore selected gene usage for the following analyses unless otherwise specified.

It has been long known that V(D)J gene segments are preferentially used or expressed, and the idea of a core gene set has been proposed (Boyd et al., 2010). However, which genes are in the core set and the extent to which they contribute to the antibody repertoire remains unclear. Taking advantage of the large data set used in this study, we plotted the gene usage of V, D, and J genes. As shown in Figure 2, although the number of V genes present in each sample varies from 15 (SRR3620039) to 99 (SRR8365259 and SRR4417619) along with the sequencing depth, we observed preferential usage of V genes. For example, IGHV3-30 and IGHV3-23 were present in all samples, while IGHV3-30-52 only appeared in one sample (SRR4417619). Accounting for both the prevalence of specific genes and their contribution to the antibody repertoire, we identified a core set of 53 V genes (Figures 2b and 2c, Materials and Methods). To our surprise, there are 3 pseudogenes, IGHV3-11 (2,147 samples, 581 donors), IGHV3-69-1 (2,128 samples, 579 donors), and IGHV3-71 (1,691 samples, 464 donors), in the core gene set. All core V genes contribute to a median of 99.33% of clones (Figure S3). The remaining V genes thus either contribute little to the repertoire or are not present. IGHJ3, IGHJ4, and IGHJ5 are present in all samples, while IGHJ1, IGHJ2, and IGHJ6 occur in 2,149, 2,150, and 2,151 samples, respectively. Of these, IGHJ4 and IGHJ6 are found in a median of 50.56% and 18.37% clones. IGHJ1 is the least used gene, contributing to a median of 2.15% clones. Three D genes, IGHD3-10, IGHD3-22, and IGHD6-19, are more prevalent. Statistics for V, D, and J gene segments are shown in Table S1, sorted by their occurrence in 582 individuals.

**Figure 2.**
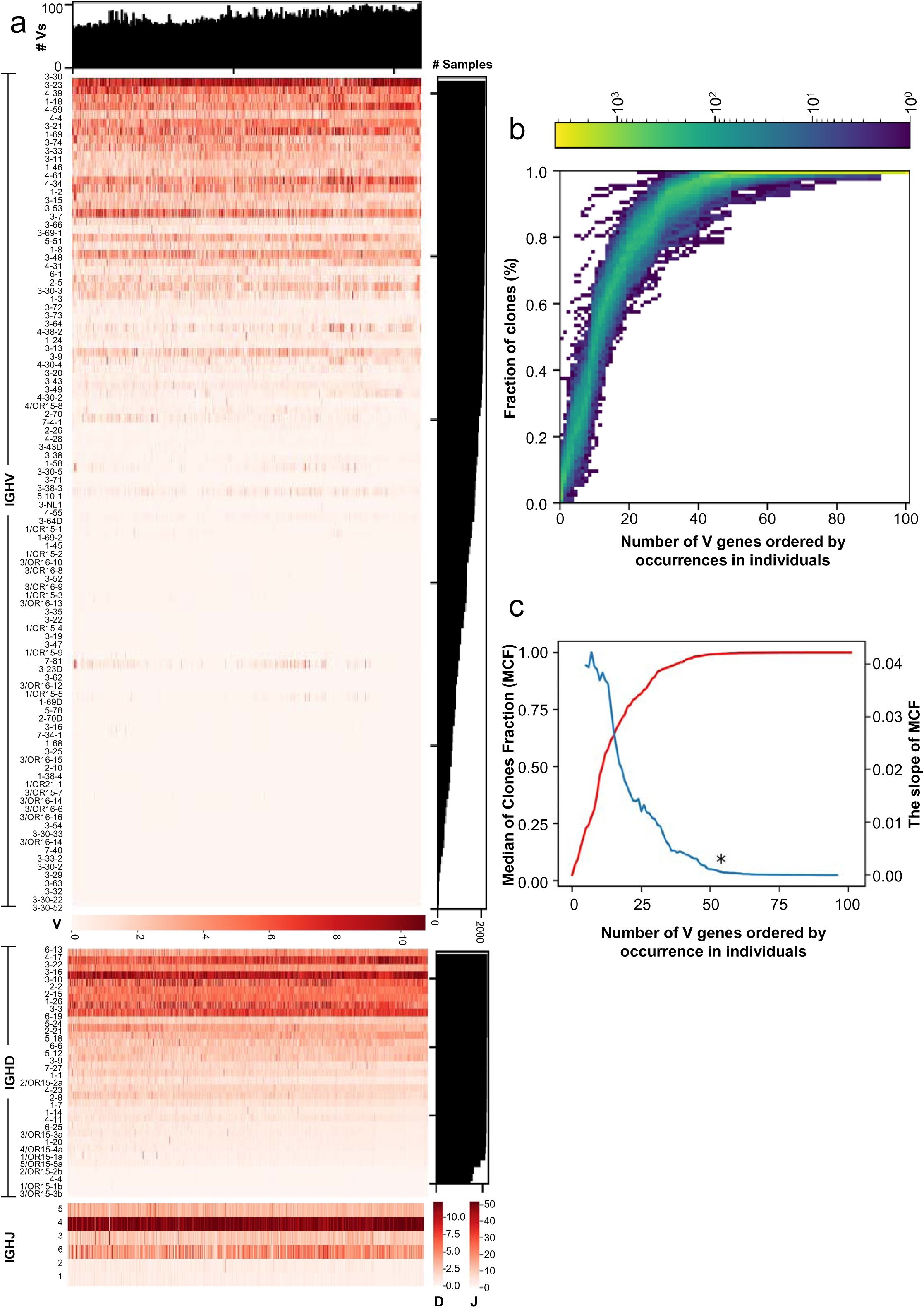
Germline gene usage and core V genes. (a) The heatmaps show the normalized usage of V (top panel), D (middle panel), and J (bottom panel) genes. Each column shows color-coded gene usage for a dataset. Each row shows usage pattern of a particular gene (IDs labeled on the left side) in different datasets. The bar graphs to the right of the heatmaps show the number of samples in which each gene was present. J genes were present in almost every sample. The bar graph on top of the V gene usage heatmap shows the number of V genes present in each sample. **(b)** and **(c)** The V genes were ordered based on their occurrences in 582 individuals from high (left) to low (right). The X-axis shows the number of high frequency V genes included. **(b)** The Y-axis shows the percent of total clones that were represents by the most frequent V genes shown on X-axis. The color shows the log 10 transformed number of samples each pixel represents. **(c)** The red line indicates the median fractions of total clones that were represents (left Y-axis) by the inclusion of top number of clones shown on X-axis. The blue line represents the slope of median clone fraction variation (on the red line) based on the adjacent 10 data points, 5 on the left and 5 on the right.

### Genetic background is sufficient but not necessary for achieving consistent germline gene usage patterns

The factors that determine V gene usage patterns have been of great interest in the field, with different studies yielding different results. By comparing the repertoires of monozygotic twins and unrelated individuals, Glanville et al. concluded that gene usage patterns are heritable, whereas Arnaout et al., Briney et al., and Laserson et al. reported that an individual’s gene usage pattern is almost identical or remarkably consistent among individuals (Briney et al., 2012; Glanville et al., 2011; Laserson et al., 2014). Thus, the effect of genetic background on V gene usage pattern is still unclear.

We therefore selected 109 repertoires, all amplified using 5’RACE, from 23 unrelated males and 3 pairs of monozygotic twins. From these repertoires, we calculated Pearson’s correlation coefficients (Pearson’s r values) for pairwise V gene usage (sample pairs from the same donor were excluded). As shown in Figure 3a, the overall coefficients of all 104 genes are higher than 53 core genes. Further scrutinizing the data revealed that most of the non-core genes had values of 0 (Figure S4a and S4b). For the male-derived samples, the minimum and maximum number of uncaptured core genes are 0 and 13, respectively, with a median of 1 and a mean of 1.53. For the non-core gene set, the minimum number is 5, with a maximum of 47, a median of 21, and a mean of 23.12. For the female-derived samples, the minimum and maximum number of uncaptured core genes were 0 and 12, respectively, with a median of 1 and a mean of 1.10. For the non-core gene set, the minimum number is 5, with a maximum of 43, a median of 21, and a mean of 22.10. These values elevated the pairwise coefficients. Therefore, we decided to use the 53 core V genes identified earlier (Figures 2b and 2c) for further analyses. The Pearson’s r values of unrelated donors ranged from 0.3681 to 0.9517, while those of monozygotic twins of the same cell type ranged from 0.9130 to 0.9952. The higher coefficient observed in monozygotic twins indicates that genetic background is sufficient to account for consistent V gene usage. However, we also observed 16 unrelated sample pairs that showed a coefficient higher than 0.9130, the minimum coefficient observed between monozygotic twins with the same cell type. Thus, a shared genetic background is not necessary for generating repertoires with very similar V gene usage. The usage patterns of D and J genes also showed the same phenomena (Figure S5a and S5b), and results were similar in the female-derived samples (Figure S5c-e). We therefore conclude that genetic background plays a critical role in defining antibody repertoire by influencing germline gene usage. However, individuals with different genetic backgrounds can also achieve a remarkably similar repertoire.

**Figure 3.**
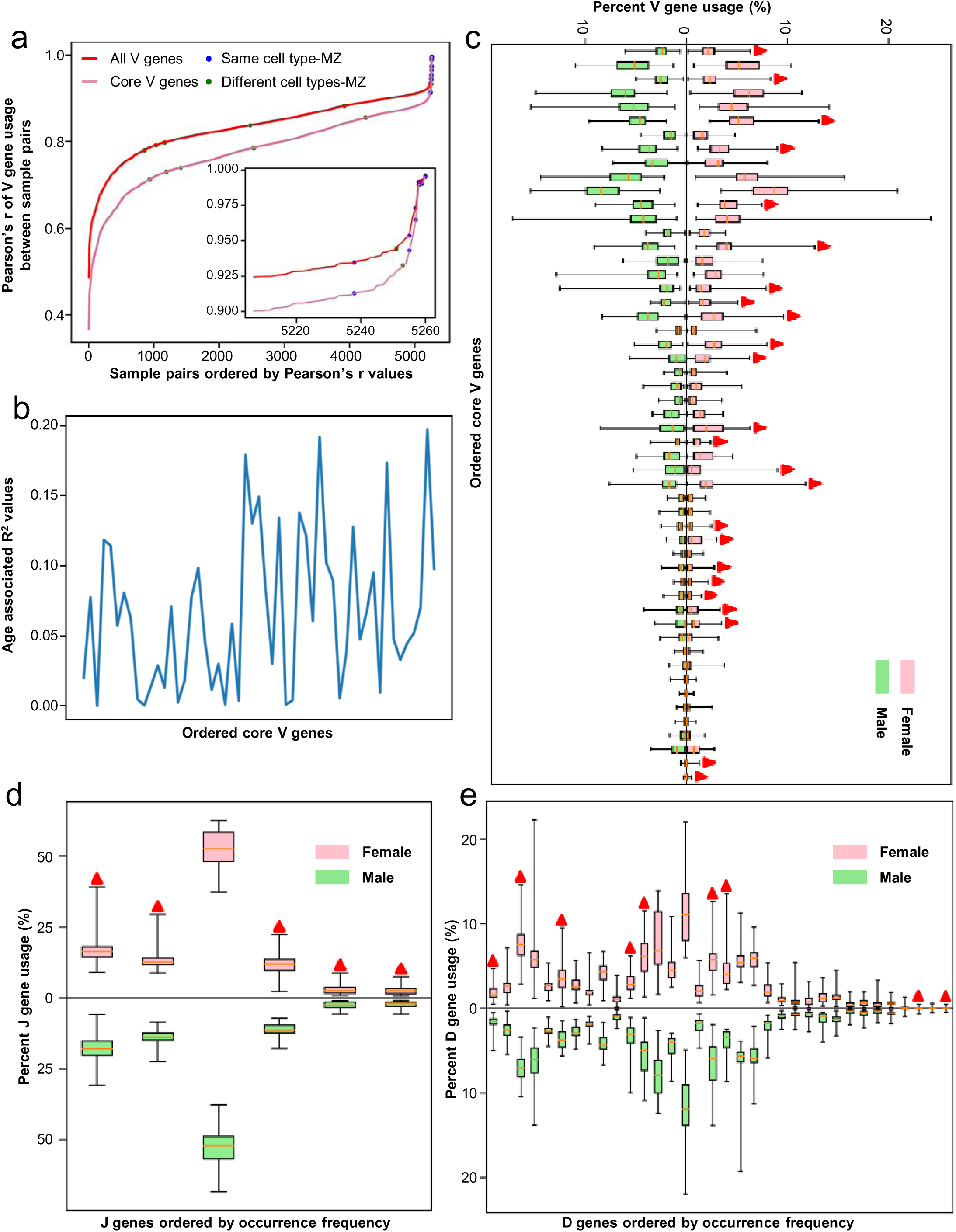
Gene usage patterns with regard to genetic background, age, and gender. (a) The Pearson’s correlation (Pearson’s r) distribution of the gene usage between 5,261 paired samples. The Pearson’s r values were ordered from low to high. The red and light pink lines represent Pearson’s r values calculated using all V genes and 53 core genes, respectively. The blue and green dots indicate the Pearson’s r values between same and different cell types for monozygotic twins, respectively. **(b)** The relationship between core V genes and ages. The X-axis shows V gene ordered by frequency (Table S1). The Y-axis indicates the R2 values calculated for a particular V gene at different ages (Supplementary. Fig. 6 and Materials and methods). **(c)**, **(d)** and **(e)** show comparisons of core V **(c)**, D **(d)**, and J **(e)** genes between male and female. The red triangles indicate genes whose usage was significantly different between sexes.

### V, D, and J gene usage shows sex and isotype preferences

After defining the relationship between gene usage and genetic background, we went on to analyze two other major factors: age and sex. Consistent with a previous study (Wang et al., 2014), our results showed that there is no linear relationship between gene usage and age, regardless of sex (Figures 3b and Figure S6a-f). To rigorously examine the impact of sex differences on antibody repertoire gene usage, we calculated pairwise germline gene usage patterns for 499 healthy PBMC samples amplified with RACE from 94 male and 164 female individuals. To our surprise, we observed that 24 core V genes (Figure 3c), 5 J genes (Figure 3d), and 9 D genes (Figure 3e) showed significant differences between male- and female-derived samples (p < 0.05, P values are listed in Table S2).

Previous studies reported that patients with influenza and SLE had characteristic changes in antibody gene usage (Pugh-Bernard et al., 2001; Sui et al., 2009). We therefore used healthy donors as background and investigated gene usage in individuals with different diseases (Figure S7a-f and Table S3). For the female-derived samples, 12 V and 8 D genes were out of the range defined by 283 healthy samples. We found that IGHV4-38-2 (SRR4026039 and SRR4026040) and IGHV3-23D (SRR4026032, SRR4026025, and SRR4026031) had increased usage in 1 and 2 out of 6 female Myasthenia Gravis patients. IGHD4-17 (SRR4026038 and SRR7230358), and IGHD3-3 (SRR4026022 and SRR4026031) were upregulated in 2 out of 6 female Myasthenia Gravis patients. For the male-derived samples, 19 V, 7 D, and 3 J genes had either higher or lower usage compared to 216 healthy male samples. For example, IGHV1-18 (H7N9_00004 and H7N9_00009) and IGHV3-73 (H7N9_00011 and H7N9_00005) showed higher usage in 2 out of 4 H7N9-infected samples. Thus, a statistical analysis of large data sets may be a powerful tool in studying the antibody repertoires of unhealthy individuals.

We also examined gene usage in different antibody isotypes, namely IgA, IgD, IgG, and IgM. There were total 51 repertoires from 5 females (14 samples) and 12 males (37 samples) available for this analysis (Figure S8). IgA and IgG were clustered together, while IgD and IgM gathered in the same subtree within the same donor. This is true for male (Figure S9a-c) and female (Figure S10a-c) samples and is consistent with a previous study (Laserson et al., 2014).

### DJ recombination shows no positional bias

During the recombination process, exonuclease trimming and the random addition of nucleotides between VD and DJ segments create diverse junctions to account for a substantial amount of antigens that may be encountered (Early et al., 1980; Tonegawa, 1983). These junctions, together with the D genes, are known as complementarity-determining region 3 (CDR3), which largely determines the binding specificity of an antibody (Chothia et al., 1989). Due to the functional importance of CDR3, there have been extensive studies looking at VDJ recombination preferences and indels in the junctions (Hansen et al., 2015; Hong et al., 2018; Saada et al., 2007; Souto-Carneiro et al., 2005; Truck et al., 2015).

For the recombination bias studies, D and J gene segments were first classified as 5D, 3D, 5J, or 3J based on their position on the chromosome. The 5D and 5J categories include the D and J segments located in the upstream region of their respective cluster. The 3D and 3J categories include the downstream D and J gene segments. Thus, 3D and 5J segments are proximal, and 5D and 3J segments are distal (Hong et al., 2018; Saada et al., 2007; Souto-Carneiro et al., 2005; Truck et al., 2015). Comparing DJ recombination between neonates and adults, Souto et al. found that 3D segments preferentially coupled to 5J segments (a proximal bias) throughout development, while 5D segments showed biased recombination to 3J segments (a distal bias) in full-term neonates rearrangements (Souto-Carneiro et al., 2005). Kidd et al. also observed a clear recombinational preference of 5D to 3J and 3D to 5J segments (Kidd et al., 2016). We thus plotted VD and DJ recombination (Figure 4a) using a total of 352 million productive clones that have D genes assigned (Figure 4b). Surprisingly, apart from the preferential usage of core V genes, IGHJ4, IGHJ6, and a few IGHD genes, we did not observe either proximal DJ or distal DJ recombination biases in our data. However, the datasets from previous neonate donors did not meet our inclusion criteria, so we cannot evaluate the positional bias of DJ recombination during neonatal development.

**Figure 4.**
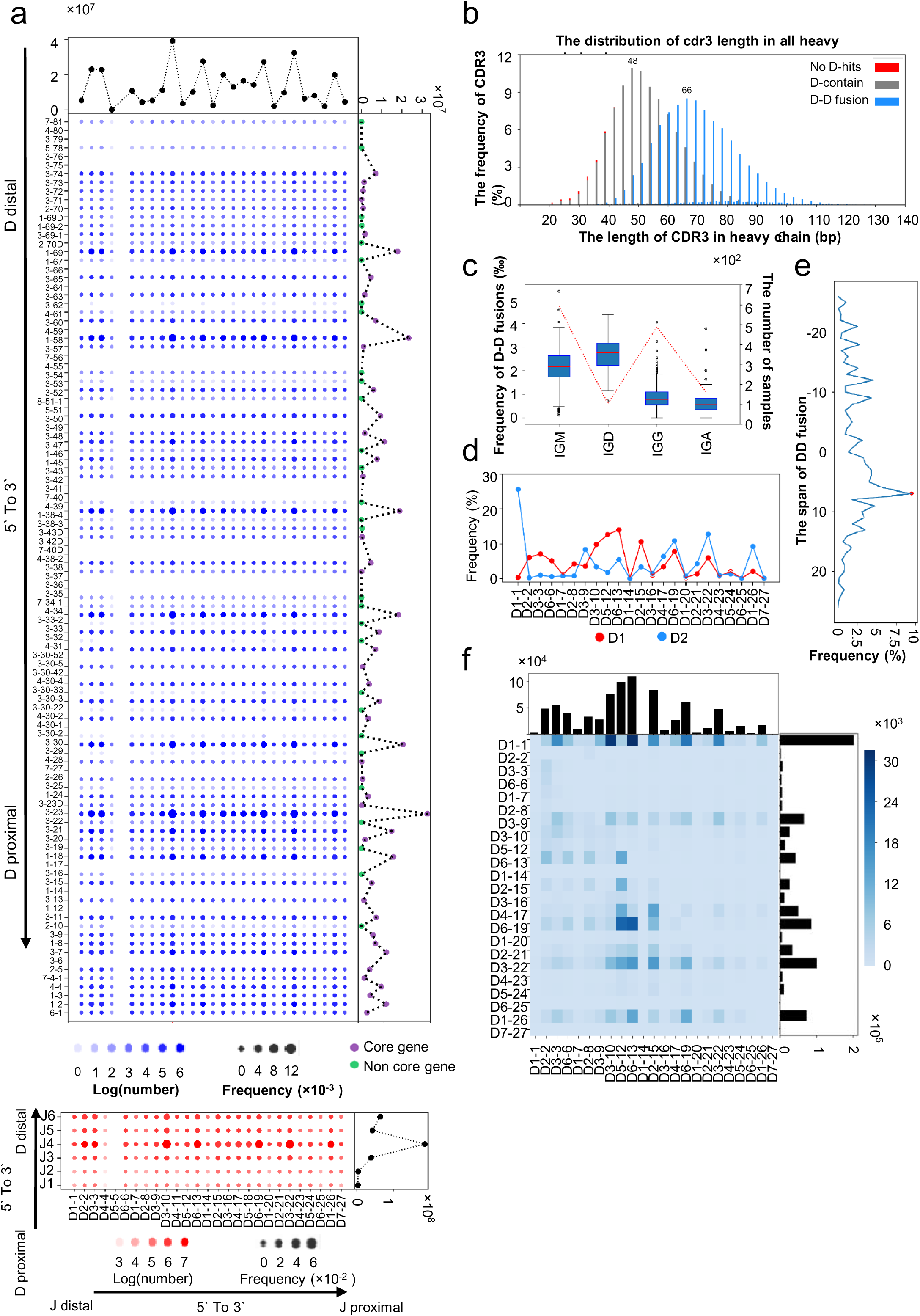
Recombination and modification between V(D)J recombination. (a) Recombination count and frequency of different VD/DJ segments. The logarithm of the count is shown by the color of the points, and the frequency of recombination is shown by the size of the points. The line at the margin shows the number of each gene segment. V genes, core genes and non-core genes are marked. The arrow shows the direction in IGH locus. **(b)** Distribution of CDR3 length in all sample, clones with whole V, D, J assignment, and DD fusion. **(c)** Frequency of DD fusion in each isotype. The line plot shows the number of samples with at least 5,000 clones in each isotype. **(d)** D gene usage in DD fusion. **(e)** Frequency of DD fusion with different span; the span of adjacent D gene is 1. **(f)** The number of DD fusions in all clones. The x axis represents the D gene at the 5’ end, and the y axis represents the D gene at the 3’ end. The bar plot at the margin shows the number of each row or column.

### D-D fusion exhibits isotype and distance preferences

D-D fusion, the incorporation of multiple diversity (D) genes during heavy chain recombination, contributes markedly to antibody repertoire diversity and has been thought to generate long CDR3 loops that frequently associate in self-reactive and polyreactive antibodies (Briney et al., 2012; Larimore et al., 2012). Briney et al. reported the first quantification of V(DD)J recombinants in naïve, memory IgM and IgG B cells from peripheral blood using Roche 454 sequencing of 4 healthy donors (Briney et al., 2012). Using stringent criteria, they found no antibodies with D-D fusion in the memory IgG population. They also reported that D gene order in cases of D-D fusion matches the order of their loci in the genome.

In bulky antibody repertoire sequencing, it is common practice to use different 3’ primers targeting different isotypes. We therefore went on to explore D-D fusion in different repertoires as well as different isotypes using IgScout (Safonova and Pevzner, 2019). We first examined CDR3 length. Total CDR3s displayed a normal distribution with a peak length of 48 nucleotides. However, the lengths of CDR3s with D-D fusions were much longer, with a peak length of 66 nucleotides (Figure 4b). Hence, D-D fusion does result in longer CDR3s.

To explore how often D-D fusion recombinants present in different antibody isotypes, we chose repertoires with at least 5,000 C gene assigned clones for corresponding isotype and calculated the frequency of D-D fusions. IgD had the highest D-D fusion frequency of 0.260% (median, n=104), followed by 0.216% of IgM (median, n=594). IgG (median, n=489) and IgA (n=163) exhibited much lower D-D fusion frequencies of 0.089% and 0.060% (median value), respectively (Figures 4c and Table S4). We did not calculate the D-D fusion frequencies for IgE because too few repertoires were available. These results are consistent with previous findings that D-D fusion recombinants may be negatively selected during isotype switching (Souto-Carneiro et al., 2005).

In contrast with previous findings, however, gene order in D-D fusion did not match the order of the corresponding loci in the genome (Figure 4d). The upstream D gene is defined as the “first” D gene (D1) in the fused recombinants and the downstream D genes could be the “second” D gene (D2). In other word, the first D gene (D1) located more 5’ in the genome prefer to be the second D gene (D2) in a D-D fusion event. However, D1 gene seem to prefer to fuse with downstream D genes with a span of 7 (The span of adjacent D gene is 1) (Figure 4e). Surprisingly, we did not observe a positive correlation between D-D fusion and D gene usage (Briney et al., 2012). The most abundant pairs were D3-10-D1-1 and D6-1-D1-1. D6-19 often served as D2, and D5-12 or D6-13 as D1. These findings may shed light on the recombination mechanistic studies.

### Stochastic recombination contributes to the public clone

Public clones are defined as to clonotypes shared by multiple individuals (Greiff et al., 2015; Jackson et al., 2013; Miho et al., 2019). It has been suggested that public clones are valuable for designing vaccines, monitoring the immune response to infection or vaccination, developing biomarker patterns of disease states, and mediating the undesirable immune responses associated with autoimmune diseases (Briney et al., 2019; Bürckert et al., 2017; Greiff et al., 2017; Maecker et al., 2012). Recent studies reported that individuals exposed to the same antigen, such as HIV, influenza, or dengue, may develop identical or similar Ig sequences – a phenomenon called antibody convergence (Jackson et al., 2014; Parameswaran et al., 2013; Setliff et al., 2018; Truck et al., 2015). Thus, a comprehensive atlas of public clones may help reconstruct the immunological history of an individual and may enable immunotherapeutic targeting within a population with a specific disease.

Ig-seq has enabled public clone studies via multiple means. Greiff et al. developed an approach that learned the high-dimensional immunogenomic features from the repertoire and enabled the prediction of public and private clones (Greiff et al., 2017). By comparing multiple donors’ ultra-deep repertoire sequencing data, Burton et al. and Soto et al. estimated the fraction of public clones in an individual to be approximately 1% and 1% to 6%, respectively (Briney et al., 2019; Soto et al., 2019). Taking advantage of the unprecedented amount of data collected for the present study, we investigated the prevalence of public clones in 2,152 samples. As shown in Figure 5a, we found that the abundance of public clones in a sample decreased when the total number of clones decreased. This result suggests that methodological undersampling may compromise the detection of public clones (Greiff et al., 2015). Furthermore, we also found that the number of public clones in two samples correlates linearly with the product of their respective clone numbers (Figure 5b) and that this correlation improves when the clone numbers for both samples increase (Figure S11a-g). The total number of clones in a given volume of blood varies with an upper boundary. Thus, getting more clones requires more blood samples. Based on different methods, the total clones in an individual’s circulating blood has been estimated to be between twenty-five million and one billion (Briney et al., 2019; Soto et al., 2019). Using our linear models with a minimum number of 5 million clones, an individual may possess between 4.2 × 10^4^ and 6.87 × 10^7^ public clones in his or her circulating blood.

**Figure 5.**
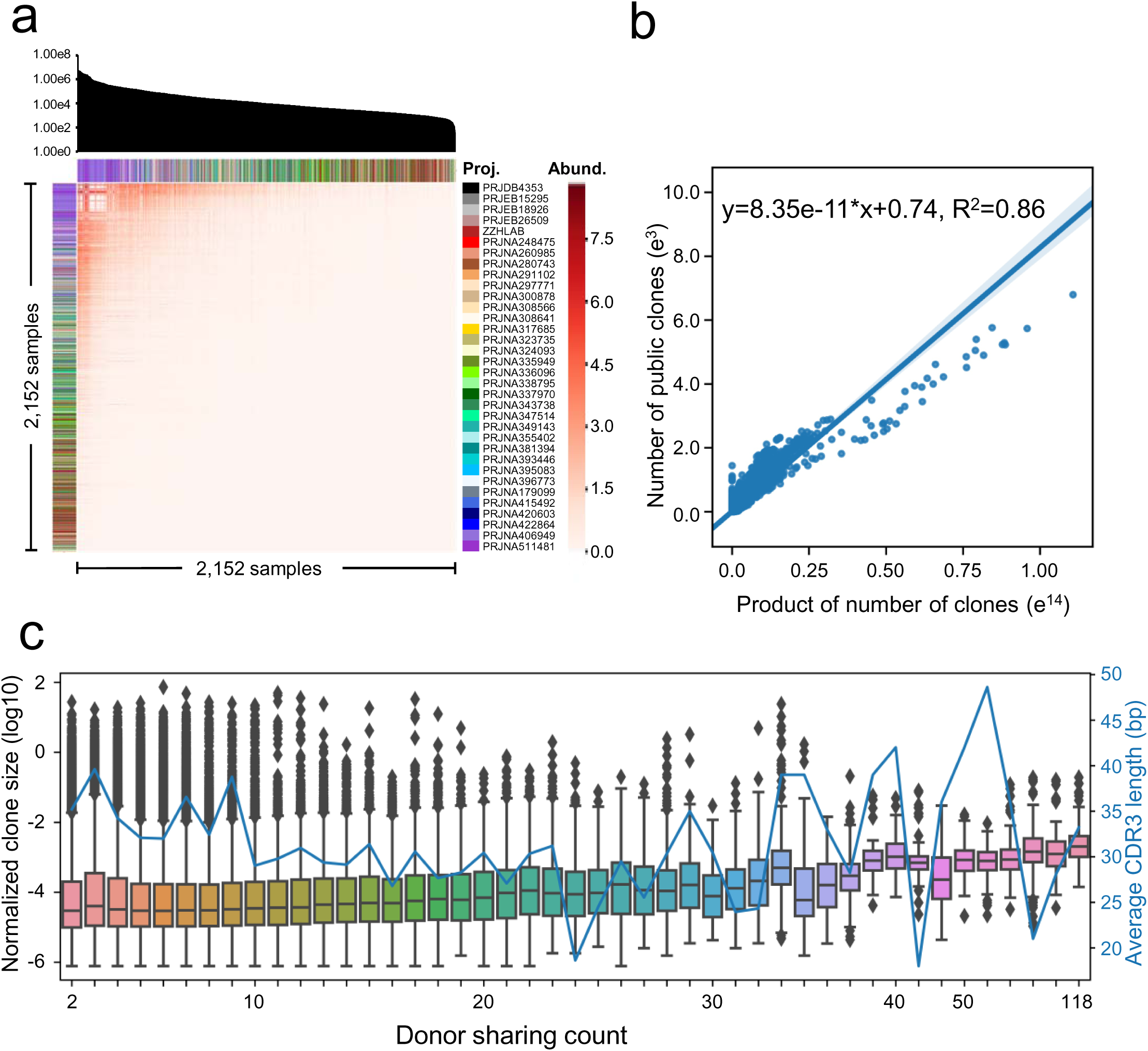
Inter-sample abundance and gene usage of public clones. (a) The heatmap in the center indicates the abundance of public clones between samples. The top bar chart indicates the number of recovered total clones for each sample. The number of public clones between each pair of sample has been subjected to logarithmic transformation (T=log(1+Pab)). The number of public clones between samples within the same project has been set to 0 to remove chimera-related effects. Note that some samples from PRJNA260985 and PRJNA280743, were predicted to come from the same donors and the observed public clones between these samples was set to 0. **(b)** Linear model delineating the correlation between inter-sample public clone abundance and the product of their clone abundance. **(c)** Public clone size percentage as a function of donor sharing count.

More in-depth analyses revealed that V and J gene usage is almost identical to the gene usage for all clones (Figure S12). On the other hand, public clones possess significantly shorter CDR3s (Figure S13). Statistical analyses of the deletions, non-template additions, and P additions showed significant differences in most elements between public clones and private clones (Figure S14). In particular, the N1 and N2 additions in public clones between the VD and DJ junctions were shorter than those of private clones. This may explain the short CDR3s in public clones and why D genes could not be assigned in many public clones (Figure S15).

Of the 162,975 (transformed to the number of unique CDR3 amino acid sequences) public clones identified in this study, 1,059 CDR3s were identical to published antigen-specific or disease-associated antibodies (Figure S16a). Further analyses showed that these CDR3s are enriched for the HIV, influenza, hematological malignancies, EBV, tetanus, and rheumatic categories (Table S5). This enrichment confirmed that antibody convergence was a source of public clones. In addition to the CDR3 enrichment in the antibodies with rheumatic autoimmune disease, we also found a CDR3 corresponding to SLE-specific antibodies in one of the healthy donors in our data. In addition, the clonotypes shared by more donors were more abundant (Figure 5c), and this change in abundance was not related to CDR3 length. Previous studies in T cell receptors (TCRs) found that shared TCRs are more likely to be autoreactive (Madi et al., 2014) and that these autoreactive TCRs are important for maintaining an individual’s health. It is possible that public clones in an antibody repertoire serve the same function. Surprisingly, we also found 31,226 (66.2%) IgM and 7,699 (70.7%) IgD clones with identical sequences compared to their respective germline V and J genes (see Materials and Methods). These clones are generated solely by VDJ recombination but have no somatic hypermutation, regardless whether the individuals have been exposed to antigen or not. This result suggests that in addition to antibody convergence, the stochastic nature of somatic recombination alone could be a key mechanism of generating public clones. We believe this collection of public clones will be helpful for studies relating to vaccine and therapeutic design targeting shared antibodies.

### Strand specificity features somatic hypermutation

Somatic hypermutation (SHM) takes place in the germinal centers of peripheral lymphoid tissues and increases the number of realizable antibodies by several orders of magnitude in addition to combinatorial diversity. The preferences and patterns of SHM allow us to trace the clonal evolution of antibodies under the selective pressure of particular antigen and to facilitate vaccine design (Schramm and Douek, 2018). The nucleotides and amino acid sites that are preferred or disfavored in SHM have been investigated using limited data and in model systems (Schramm and Douek, 2018). SHM in the antibody repertoire results from two types of sequential events. First, activation-induced cytidine deaminase and other molecular components of the SHM machinery introduce mutations to the antibody variable regions. The selective pressure of a particular antigen then acts on these mutations and preserves the favored ones. Thus, the majority of SHM studies worked from unproductive reads to emphasize the effect of mutations rather than the effect of antigen selection. This is particularly beneficial for mechanistic research on SHM because it simplifies the model. However, antigens only place selective pressure on antibodies containing mutations and do not introduce additional mutations. Despite some antigens that may preferably retain rare mutations, the clear majority of mutations in functional antibodies would also reflect the selective flavors of the SHM machinery. Moreover, the selective bias only acts on antigen-specific clones. Thus, this bias would be minimized if the effects of clonal expansion are compensated or removed during computational analyses.

With this in mind, we used consensus sequences and a position weight matrix (PWM) to represent a clone and probed mutations in the V genes at different levels (Materials and methods). We first depicted the mutational propensities at the single nucleotide level (Figures S17a and S17b). Three types of transitions (*A* to *G*, *G* to *A*, and *C* to *T*) occurred with high frequencies, and *T* to *C* mutations occurred at a significantly lower frequency. Transversions between purines and pyrimidines were less frequent, with the exception of *G* to *C* mutations. While *C* and *G* showed comparable mutability, *A* exhibited significantly higher mutability than T. Because *A*: *T* and *G*: *C* present as pairs on the chromosome and SHM occurs at the DNA level, it is interesting to observe that the mutational tendencies are not reciprocal.

Having observed disproportional mutation tendencies in nucleotide pairs, we investigated the mutability of reported motifs in a strand-specific manner (Material and methods). It is worth noting that all nucleotides and motifs were extracted from the forward V gene sequences, and the reverse sequences were discarded. Every nucleotide was classified exclusively in a single motif. Therefore, the bases between categories have no overlap. We confirmed that SY<u>C</u> (where S is C or G; and Y is C or T) and <u>G</u>RS (where R is A or G) are *bona fide* coldspots that showed the lowest frequency of mutations. The motifs WR<u>C</u>Y, R<u>G</u>YW, W<u>A</u>, and <u>T</u>W also showed much higher mutations than did coldspots as reported by others (Liu and Schatz, 2009; Pham et al., 2003). However, significantly different mutabilities were observed again between reciprocal motifs (WR<u>C</u>Y and R<u>G</u>YW, W<u>A</u> and <u>T</u>W) (Figure 6a, Figure S17c and Table S6). This result suggested strongly that SHM is introduced in a strand-specific manner.

**Figure 6.**
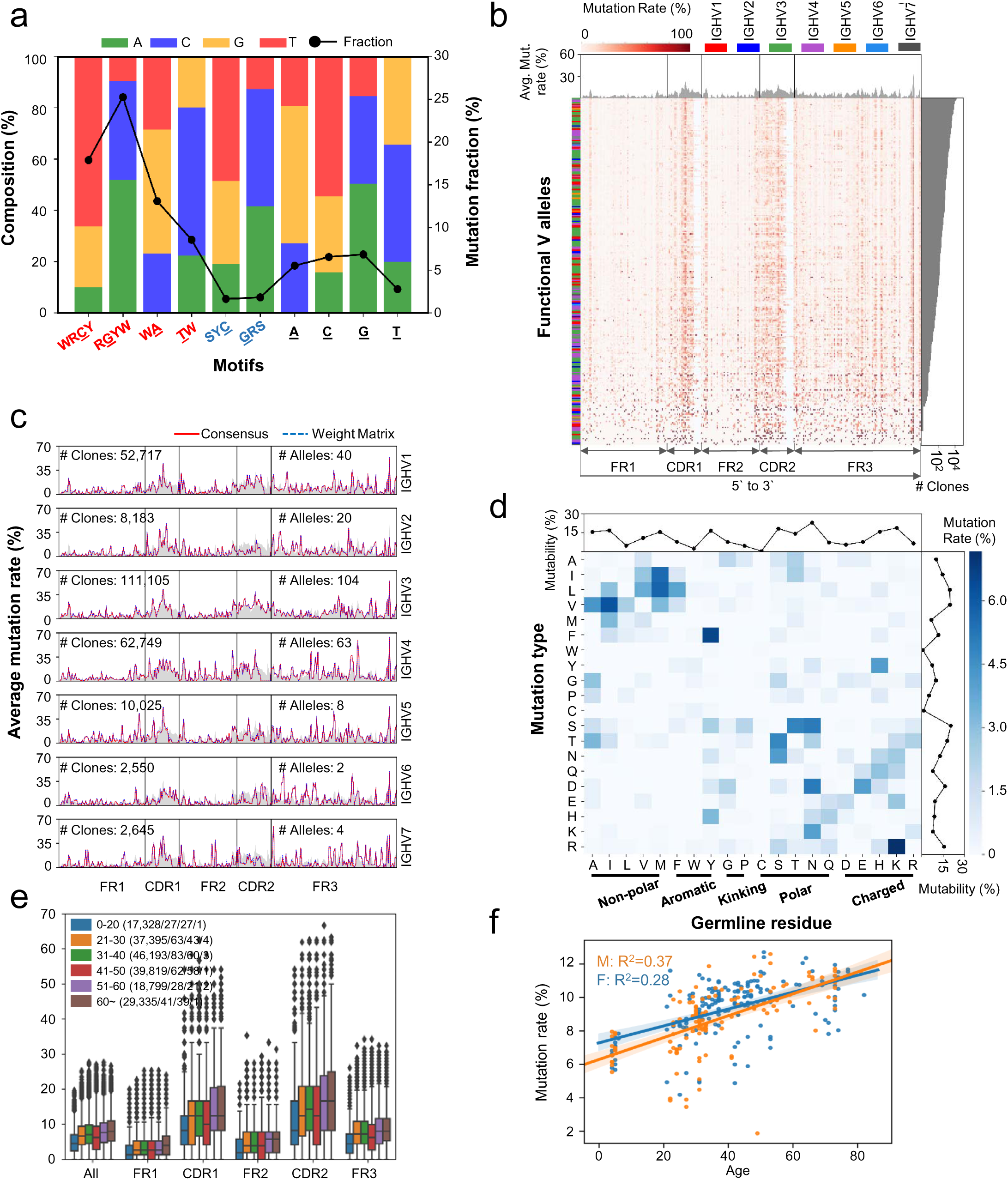
Somatic hypermutation patterns and influence factors. (a) The stacked column diagram shows the mutation percentage of motifs and composition of mutation targets. The X axis shows the different motifs in germline sequences. The Y axis shows the composition of the mutated nucleotide of this motif. The line chart shows the mutation fraction of every motif. The red-colored label represents hot-spot, the blue colored label represents cold-spot. The underlined letter represents the mutation site. **(b)** and **(c)** show the mutation rate among different functional alleles and families. **(b)** The combined heatmap shows the mutation rate among used functional alleles in selected IgG samples. Each column shows the position of completion of the V segment from FR1 to FR3. Each row shows the functional alleles occurred in datasets. The area chart represents the average mutation rate in every position. The bar graph left to the heatmap shows the family of occurred alleles which ordered by the number of clones who were shows in the right bar graph. The color of the heat map represents the mutation rate of every position from used functional alleles. **(c)** The X axis shows the position of the V segment from FR1 to FR3. The Y axis shows the average mutation rate from different families. The area chart shows the overall average mutation rate about used functional alleles. The red lines and blue dotted lines show the result of the mutation rate of every family based on consensus and weight matrix methods. **(d)** The combined heatmap shows the substitution among amino acid. Each column and each row represents an amino acid. The germline residue is located on the x axis, and the mutated amino acid is located on the Y axis. The line graph represents the ability of each amino acid to be mutated and mutated. **(e)** The boxplot shows the mutation rate for different age groups across multiple functional region and whole region. The points on top of each boxplot indicates the outliers. **(f)** The scatter plot (orange for male and blue for female) shows the correlation between mutation rate and age. Two lines in the figure are the predicted linear regression model for male and female. R-squared value were marked on the top left in this figure.

Complementarity-determining region (CDR) 1 and CDR2 exhibited higher mutations as expected (Figure 6b and Figure S17d). While framework region (FR) 1 and FR2 displayed lower mutation rate in contrast with CDRs, the region immediately adjacent to the CDR regions was also subjected to a high frequency of mutations. These results support the idea that FRs provide the backbone of the antibody, while CDRs accumulate mutations to achieve high affinity binding to a target antigen. Consistent with previous observations (Shapiro et al., 1999), there was a considerable amount of SHM in the FR3 region. Interestingly, the base with highest mutation rate was found near the end of the FR3. A closer look at the germline sequence revealed that this nucleotide represents the third position within a codon. The space for nucleotides, associated codons and amino acids was dominated by *G*, *GTG*, and *V (valine)*, respectively (Figures S18a-c). Interestingly, however, in most cases, this site does not occupy any previously identified canonical hotspot (Figure S18d). Nucleotide substitution analysis at this locus showed no preference and has no impact on the encoded amino acid except for eight alleles in the IGHV5 gene family (Figures S18e-g and S18h). In addition, the mutation spectrum profiles varied in different IGHV families (Figure 6c).

Although the conservative substitution (transition within the same amino acid group) dominated amino acid substitution profile, we observed relatively-high frequencies for non-conservative substitutions, such as *H* to *Y* and *N* to *D*, that were not identified before using limited number of datasets. In addition, we found *W* and *C* were least mutated (3.95% and 1.63%) and mutated to (1.00% and 2.17%) (Figure 6c and Figure S17e).

The level of SHM as a function of confounding factors, such as age, sex, and isotypes, has also been explored to some extent (Jiang et al., 2013; Kitaura et al., 2017; Wang et al., 2014). Nonetheless, there have been no studies to date profiling SHM from a large data set. Using 363 samples from 290 donors, we reviewed the role of age, sex and isotypes on the frequency of SHM. We found negligible differences between males and females, except for those between 41-50 years of age which probably result from uncommon sampling bias. (Figure S19). We therefore combined data from male and female donors to investigate the effects of age and isotype. Switched isotypes, namely IgG, IgA, and IgE, had comparable level of SHM (7 - 8%), while IgM and IgD had much lower level of SHM (1 - 2%) (Figure S20), consistent with a previous report (Kitaura et al., 2017). We also observed a positive correlation between the level of IgG SHM and age, except for individuals in the 41-50 age range (Figure 6e). When we measure the contribution of age to SHM levels using a linear model, we obtained r-square values of 0.37 and 0.28 for male and female, respectively. Despite the compromised goodness of the model, we confirmed this correlation at the population level and estimated that the SHM increases by approximately 0.05% each year. This increase would mean that in general, a parent bears 1% more SHM than their children (Figure 6f).

## Discussion

Extremely large data sets have proven to be powerful for computational analysis to reveal patterns, trends, and associations. The advent and application of high-throughput sequencing technology advanced the study of complex biological systems, launching projects such as the 1000 Genomes Project, The Cancer Genome Atlas, the Encyclopedia of DNA Elements, and the NIH Human Microbiome Project. Inspired by the success of these projects, we systematically analyzed the largest antibody repertoire dataset to date and scrutinized, for the first time at this scale, the key features of the antibody repertoire.

In addition to the uneven usage of germline genes, we identified a set of core V genes that contribute to the clear majority of the repertoire. Although the other V genes are less frequently observed in the current datasets, we believe their absence is the result of shallow sequencing depth compared to the complexity of antibody repertoire. Nonetheless, these core and “rare” V gene sets may serve as a reference for discovering gene usage fluctuations that are associated with or specific to particular diseases. We found that the number of public clones between two repertoires also relied on the sequencing depth. Moreover, a fraction of public clones identified in repertoire comparisons were reported to be disease-associated or antigen-specific. This result supports the notion of antibody convergence and also suggests that the antibody repertoire may help us trace an individual’s immune history and may therefore be useful in selecting vaccines and immunotherapy for certain diseases.

The fundamental B cell biology that underlies the specific patterns of germline usage, D-D fusion, and SHMs revealed in our analyses remains controversial. Follow-up experiments may reveal the mechanisms behind these phenomena and thus advance our understanding of B cell development as well as its response to immune perturbations.

Due to the intrinsic amplification bias caused by different amplification strategies and various primer sets, we did not perform analyses of clonal expansion, diversity, and evenness. A common standard for both experimental design and bioinformatics analysis will be critical for future studies.

The human antibody repertoire possesses extreme diversity. Compared to the aforementioned prior studies, the number of samples analyzed here is far from sufficient to capture all this diversity. Moreover, antibody repertoires from a broad spectrum of diseases as well as different isotypes from various tissue types are still needed for a better understanding of humoral immunity. Due to the limited source of human samples, it is likely that further studies with model systems such as mouse, rat, and macaque will bring us more insights.

## Acknowledgements

This study was supported by the National Natural Science Foundation of China (NSFC) (31771479) (Z. Z.), NSFC Projects of International Cooperation and Exchanges of NSFC (61661146004), the Local Innovative and Research Teams Project of Guangdong Pearl River Talents Program (2017BT01S131), the Thousand Talent Plan of China, and the Guangdong Natural Science Funds for Distinguished Young Scholar (2017A030306030), the Guangdong Innovative and Entrepreneurial Research Team Program (2016ZT06S638), the National High Technology Research and Development Program of China (2012AA02A206), National Natural Science Foundation of China (NSFC) (81822036 and 31770931) (W. Y.), the National Science Foundation for Excellent Young Scholars (81222035), the National Program for Support of Top-Notch Young Professionals (X. B.), the Chang Jiang Scholars Program (X. B.), the Special Support Program of Guangdong (X. B.).

## Author Contributions

X. Y., Y. Z., H. Z., Y. Z., C. L., J. W., C. M., and Y. Z. performed the bioinformatics analyses on the data. M. W., D. S., C. L., Y. D., S. G., L. X., R. W., and J. O. collected blood samples and conducted the biological experiments. M. W., S. G., R. W., and C.J. L. prepared the libraries and performed Illumina sequencing. C. C., W. Y., H. Z., J. C., L. Q., H. Z., J.X. B., L. W., G. C., X. Y. and Z. Z. designed the project, biological experiments as well as bioinformatics analyses. X. Y., M. W., D. S., Y. Z., H. Z., Y. Z., C. L., G. C., X. Y. and Z. Z. co-wrote the manuscripts.

## Materials and Methods

### Dataset enrollment criteria

We searched for bioprojects that were related to the antibody repertoire on the Sequence Read Archive (SRA: https://www.ncbi.nlm.nih.gov/sra) from the National Center for Biotechnology Information (NCBI: https://www.ncbi.nlm.nih.gov/). We identified thirty-eight projects before Feb 28, 2019. The datasets from the included projects were subjected to two consecutive filter processes. The first filter procedure was based entirely on sample metadata provided by SRA and the corresponding papers. The criteria include:

- Homo sapiens
- Illumina platform
- Pair-end Library Layout
- Sequencing length >= 250
- Natural sample directly extracted from human tissues (excludes those samples derived from cell lines)
- No specific amplification
- Library source is either GENOMIC or TRANSCRIPTOME
- No spike-in sequences

The second filter procedure was based on the results when preprocessing finished, criteria here consists of,

- Number of productive reads for heavy chain > =10,000
- Fraction of heavy chain >= 20%

### In-house dataset

#### Subjects

A total of 295 peripheral blood mononuclear cells (PBMCs) samples were collected. Of these, 254 were derived from healthy individuals (without recent infection events), 18 were from HBV patients, 16 were from H7N9 patients, 6 were from individuals involved in traffic accidents, and 1 was from a patient with meningitis. Peripheral blood samples (1 ml) obtained from each volunteer were collected in an EDTA-containing sterile tube and stored at room temperature for no more than 6 hours. PBMCs were isolated by Ficoll-Paque density centrifugation using Lymphoprep™ (Axis-Shield, 1114547), and the isolated cells were lysed in RLT buffer (Qiagen) supplemented with 1% β-mercaptoethanol (Sigma) before being stored in −80_LJ_ for short-term storage. This protocol was approved by the Ethics Committee at Southern Medical University. Informed consent was obtained from all participants.

#### RNA extraction, reverse transcription, 5’RACE amplification, and next-generation sequencing procedures

RNA purification was carried out using the RNeasy Mini Kit (Qiagen, 74106) according to the manufacturer’s instructions. The concentration of the RNA was determined using a NanoDrop 2000c Spectrophotometer (ThermoFisher Scientific). Five hundred nanograms of RNA purified from each sample was used for cDNA synthesis with a total volume of 20 µl. cDNA was prepared using a SMARTer RACE cDNA Amplification Kit (Clontech, 634928) according to the manufacturer’s instructions. Forward primers were synthesized according to SMARTer RACE protocol. The first 50 bp of the first constant domain (CH1) of heavy chain (IgG) were used to design the reverse primers. We also designed 8-11bp barcode at the upstream of these primers to distinguish samples. One microliter of the reverse transcription mixture was used as a template in a 20 µl PCR reaction. Primers were used at a final concentration of 100 nM. The thermal cycling conditions were programmed as follows: denaturation at 95°C for 3min, 30 cycles of denaturation at 98°C for 20s, annealing of primer to DNA at 60°C for 15s, and extension by Kapa HiFi HotStart Ready Mix (KAPA Biosystems, kk2602) at 72°C for 15s, followed by a final extension for 5 min at 72_LJ_. PCR products were analyzed on a 1.5% agarose gel, and the appropriate bands (∼600 bp) were purified using the Nucleospin Gel & PCR Clean-up kit (Macherey-Nagel, 704609.25). DNA Concentration was measured using the NanoDrop 2000c Spectrophotometer (Thermo Fisher Scientific), and 400 ng of DNA was used to prepare libraries using a Universal DNA Library Prep Kit for Illumina V3 (Vazyme, ND607-01), strictly following the manufacturer’s instructions. Libraries were quantified using the Qubit 4.0 fluorometer (ThermoFisher Scientific) and re-quantified using the KAPA qPCR kit (KAPA Biosystems, 4824). The size of adapter-ligated DNA fragments (approximately 800 bp) was determined using a Bioanalyzer 2100 system (Agilent). Each library was subjected to 2 × 300 bp paired-end sequencing using MiSeq Reagent V3 kits (Illumina, MS-102-3003).

#### Germline gene assignment and clonotype assemble

Paired-end FASTQ files downloaded from SRA and generated by our laboratory were inputted into *MiXCR* (version 3.0.7) and run with the following parameters:

Align: *mixcr align --library my_library -t 8 -r align_log.txt R1 R2 alignments.vdjca -s hs* Assemble: *mixcr assemble -r assemble_log.txt -OseparateByV=true -OseparateByJ=true -OseparateByC=true alignments.vdjca clones.clns*

Export clones: *mixcr exportClones clones.clns clones.txt*

Export Alignments: *mixcr exportAlignments -f -readIds -vHit -dHit -jHit -cHit -vGene -dGene -jGene -nFeature CDR3 -aaFeature CDR3 -defaultAnchorPoints alignments.vdjca alignments.txt* We built germline references for V, D, J, and C gene segments locally, and the germline references for V, D and J gene used in this study were customized using *repseqio* (v1.2.12, https://github.com/repseqio/repseqio). Reference sequences were obtained from IMGT/GENE DB (http://www.imgt.org/genedb/) and are provided in Table S7. The formatted information for the reference constant region sequences was directly extracted from the *MiXCR* built-in reference (v 1.5) and then appended to the formatted customized reference for V, D and J genes. *MiXCR* clustered sequences with the same V, J, and C allele assignment and CDR3 nucleotide sequence into a clone with the parameters above. An in-house *Python* script was used to merge clo nes with the same V and J gene and CDR3 nucleotide sequence into a clone. If we investiga ted isotypes effect on some indices such gene usage, the C gene was also taken consideration.

### Comparison of gene usage and expression between Multiplex and RACE

Gene usage was defined as the number of clones with a given gene segment divided by the total number of clones. Gene expression was defined as the number of reads with a gene divided by the number of productive reads. For each gene segment, the median of usage and expression for either Multiplex or RACE was used for linear regression. Usage or expression from Multiplex was defined as independent variable while that from RACE was considered as dependent variable. The *regplot, r2_score*, and *pearsonr* functions in *seaborn* (version 0.9.1) and the *sklearn* (version 0.20.2) and *scipy* (version 1.2.1) *Python* modules were used to visualize the linear regression and to calculate R squared values and a Pearson Correlation Coefficient.

### Overview and core gene set selection of gene usage

To show gene usage for all 2,152 samples clearly, we set thresholds for V, D, and J genes. If the usage was greater than the threshold, we used the threshold value instead of the original value. The average of the maximum of each sample for V, D, and J gene were calculated as thresholds. For core V gene set selection, we first sorted genes according to their occurrence in 582 donors. We then enrolled 102 V genes one by one and computed the accumulated clone fraction with specific V genes for 2,152 samples. The median of clone fraction for all samples was selected and the slope of them was computed. The slope of *x_i_* was equal to the distance of clone fraction at *x_i-5_* and *x_i+5_* divided by 11. Finally, if the slope was less than 0.001, we determined the clone fraction arrived the plateau and chose this gene set as the core gene.

### Features’ effect on gene usage

#### Genetic background

Two hundred and twenty-two healthy peripheral blood mononuclear cell samples, obtained from 29 male and 22 female individuals from 21 to 30 years of age, were amplified by RACE and used to explore how genetic background affects gene usage. We examined two gene sets containing 53 core V gene and 102 V genes. The Pearson Correlation Coefficient for every sample pair was calculated using *pearsonr* from a Python module named *scipy* (version 1.2.1). Sample pairs from the same donor were excluded. We performed statistics for male and female samples separately.

#### Age

We chose 499 healthy PBMC samples drawn from 94 male donors and 164 female donors by RACE. Linear regression was conducted using *LinearRegression* and *r2_socre* from *sklearn* (version 0.20.2) module. The independent and dependent variables were age and gene usage (of 53 V genes, 34 D genes, and 6 J genes), respectively. Samples derived from males and females were analyzed separately.

#### Sex

We performed two independent sample t-tests for 53 V genes, 34 D genes, and 6 J genes on the 499 samples selected above using *ttest_ind* from *scipy* (version 1.2.1). Genes whose P values were less than 0.05 were defined as differentially used in the male samples and the female samples.

#### Isotype

We first obtained isotypes composition including IgA, IgD, IgE, IgM, and IgG for the 499 samples above. Because the fraction of IgE was too low to compare, we discarded this isotype. Based on the isotype fraction, there were 14 female and 37 male samples that could be used for this analysis. We then merged clones for each isotype from different samples derived from the same donor and recalculated gene usage for them. Gene usage was regarded as a vector, and *Euclidean distance* was calculated using *DistanceMatrix* from *scikit-bio* (version 0.5.5) to measure the similarity of gene usage between different isotypes. The *nj* function from *scikit-bio* was used to build a neighbor joining tree. Finally, we used *Dendroscope* (version 3.6.3) to generate the trees. Samples analyzed for gene usage related to genetic background, age, and so on, are shown in Table S8.

#### VD/DJ recombination

To compare the recombination bias in all clones, we only analyzed clones with a D gene assignment. Clones with full V, D, J gene assignments were pooled together. Clones without stop codons or out-of–frame mutations in the CDR3 region were considered to be productive clones. If multiple assignments occurred, the gene with the highest score was used for the analysis. The clones were separated into subgroups according to VD/DJ recombination, and the frequency of each group was calculated. The number of each gene was calculated according to about 352 million productive clones.

#### D-D fusion detection

*IgScout* was used to detect D-D fusion. Input files were extracted from the *MiXCR* results using a custom generated script, which used default parameters and the same reference as *MiXCR*. No other filter was used to detect D-D fusion. The identity of D-D fusion and alignment length were calculated by a custom script written in python. Levenshtein distance was used to quantify the difference between the reference and the aligned sequences. The length of the aligned sequence was calculated directly from the result of *IgScout*.

#### Position bias in D-D fusion

Tandem CDR3s from all samples were pooled together to calculate gene usage in all D-D fusions. We defined D1 as the D gene at the 5’ end and D2 as the D gene at the 3’ end. The span of two D genes was defined as the genes position number between 5’ D and 3’D on the corresponding chromosome. The span of two adjacent D genes was 1. A negative value indicated that the D1 gene was located at the 5’ end of D2 on the chromosome, and a positive value represented a D2 gene located at the 3’ end of D1 on the chromosome.

#### Comparison of D-D frequency between isotypes

To compare the D-D frequency between isotypes, we also included C gene annotations. Due to the low frequency of D-D fusion, we only included samples that contained at least 5,000 clones for corresponding isotypes. Our analysis included 104, 594, 489, 163 samples for IgD, IgM, IgG, and IgA, respectively. IgE was not included in the analysis because none of the samples met our criteria of at least 5000 annotated clones. The frequency of D-D fusion in each sample was calculated as the number of D-D fusions in the corresponding isotype divided by the total number of corresponding isotypes.

#### CDR3 length distribution

The number of nucleotides in each clone was defined as the CDR3 length of the clone. The distribution of all clones was showed calculated from total clones. CDR3 clones with a D gene assignment were calculated as D-containing CDR3. The length of Tandem CDR3s was calculated from an output file named tandem_cdr3s.txt generated by *IgScout*. To make the distribution comparable, the frequency of CDR3s in each length group was used.

#### Public clone abundance profile

Public clones were those clonotypes (defined before) shared by at least two donors from two or more projects. Therefore, number of public clones between two samples from the same project or the same donor was set to zero. This strict criterion for clonotypes was applied to remove ‘public’ clones resulting from chimeric artifacts.

#### Linear model to delineate the stochastic nature of gene recombination

Linear models were constructed using only valid sample pairs that derived from different donors in different projects. Associated coefficients for regression equation and R squared were estimated using function *linear_model.LinearRegression* within the Python package *sklearn* (version 0.20.2).

#### Profile of gene usage, CDR3 length, and Junctional Modification

Non-redundant public clones were used to profile gene usage and CDR3 length distribution, while redundant public clones were used for junctional modification analysis. The junctional modification calculation method is the same as above. Statistical analysis was carried out using a two-tailed unpaired Student’s t-test.

#### Antigen- or disease-related antibody database overlapping

We generated custom antigen- and disease-related antibody databases (unpublished results). All curated antibody sequences (heavy chain) were collected from following databases: i) IMGT/LIGM_DB (http://www.imgt.org/ligmdb/); ii) abYsis (http://www.bioinf.org.uk/abysis3.1/index.html); iii) EMBLIG (http://acrmwww.biochem.ucl.ac.uk/abs/abybank/emblig/); iv) bNAber (http://bnaber.org/), v) HIV_DB (http://www.hiv.lanl.gov/); vi) NCBI Nucleotide database (https://www.ncbi.nlm.nih.gov/nuccore); and vii) EBI ENA (https://www.ebi.ac.uk/ena). For now, it is comprised of 65,088 non-redundant antibody heavy chain sequences, corresponding to 53,579 unique CDR3 amino acid sequences and 163 types of antigen or disease, including HIV, hematological malignancies, preterm birth, influenza and etc. Antigen- or disease-related enrichment analysis of overlapping antibodies was performed using a hypergeometric model, implemented with the *stats.hypergeom.cdf* function within the Python package *scipy* (version 1.2.1). The false discovery rate was calculated using the *Benjamini-Hochberg* method implemented with an in-house script.

### Somatic hypermutation

#### Sample selection

Only samples which were from healthy donors’ PBMC and were amplified using RACE protocol were included in the somatic hypermutation analysis. Since some experimentally qualified datasets with which the alignment information failed to be exported, 363 samples were included in the end (Table S8).

#### Export alignment

Alignment information used to measure somatic hypermutation was exported using *MiXCR* with the following parameters:

Assemble: *mixcr assemble -r assemble_log.txt -OseparateByV=true -OseparateByJ=true -OseparateByC=true -a alignments.vdjca clones.clna*

Export Alignments: *mixcr exportAlignments -f -readIds -cloneId -vHit -vAlignment -jHit -jAlignment -cHit -cAlignment -nFeature FR1 -nFeature CDR1 -nFeature FR2 -nFeature CDR2 -nFeature FR3 -nFeature CDR3 -nFeature FR4 -aaFeature FR1 -aaFeature CDR1 -aaFeature FR2 -aaFeature CDR2 -aaFeature FR3 -aaFeature CDR3 -aaFeature FR4 -defaultAnchorPoints clones.clna alignments.txt*

#### Quality filtering and data preprocessing

1. Read QC. Removal of reads that could not be merged by *MiXCR*, those without complete variable regions (VR), those having been assigned with a pseudogene or a different V assignment compared with their corresponding clones’, those containing insertions or deletions and those with stop codons or frameshifts in the variable region.
2. Clone QC. Removal of clones with only single qualified reads following read QC procedure above.
3. VR deduplication. Deduplicating VR to obtain non-redundant sequence set
4. VR grouping. Grouping VR according to the isotypes reported in the clone files.

#### Implementation of consensus and position-weighted matrix approaches

The position-weighted matrix approach considers all qualified non-redundant reads within each clone. Because each clone was a basic unit in the somatic hypermutation analysis, the mutation rate for a certain position was calculated as the sum of mutation rate for all mutation events observed within reads supporting this clone. The number of substitution types for a nucleotide (nt) or an amino acid (aa) for a certain position was defined as 1 if a nucleotide or amino acid in a given position underwent the same substitution event for all reads within a clone (with the same target nt or aa), otherwise it would have a value less than 1.

For every clone, a theoretical consensus sequence was calculated based on the *motifs* module in *Biopython* (version 1.73). The Hamming distance was used to calculate the distance between the theoretical sequence and each true sequence, where the true sequence closest to the theoretical sequence was taken as the representative sequence of the clone.

### Software

In-house scripts were written in Python (version 3.7.4) based on the numpy (version 1.16.4), Biopython (version 1.73), Levenshtein (version 0.12.0) and pandas (version 0.24.2) modules. To visualize these results, we used the Python modules seaborn (version 0.9.1) and matplotlib (version 3.0.2) as well as GraphPad Prism (version 7.04).

## Supplemental Information titles and legends

**Figure S1. Age (a) and sex (b) composition of enrolled donors.** Total number of enrolled donors is 582.

**Figure S2. The Pearson correlation coefficients of gene expression and usage between Multiplex and RACE.** Left: the distribution of Pearson correlation coefficients of V (a), D (b), and J (c) gene expression between Multiplex and RACE. Right: Pearson correlation coefficients of V (d), D (e), and J (f) gene usage between these two groups. Note: There are 1,409 datasets amplified by Multiplex and 743 datasets amplified by RACE.

**Figure S3. Fraction of repertoire containing the 53 core V genes.**

**Figure S4. Number of uncaptured V genes for 109 male (a) and 113 female (b) samples.**

**Figure S5. The Pearson correlation coefficients of D (a), J (b) for male and V (c), D (d), and J (e) for female.** Lines in red show all genes, and the line in light purple shows the core V genes. The dots indicate monozygotic twins from PRJNA300878. The blue dots indicate the same cell type of monozygotic twins, and the green dots indicate different cell types (naïve and memory) for them.

**Figure S6. Effect of Age on gene usage for females and males.** The R square of linear regression between D (a), and J (b) gene usage and age in female. (c) The scatter plot for 53 V, 34 D, and 6 J genes usage and age. The X-axis means the age and the Y-axis stands for the usage. R squared for V (d), D (e), and J (f) usage and age in the male.

**Figure S7. Gene usage in the infected and uninfected samples of the female (a, b, and c) and male (d, e, and f).** Boxplots show usage from uninfected samples, while the red dots represent usage from infected ones.

**Figure S8. The isotype composition of clone from the male (a) and the female (b).** Each column shows one isotype including IgA, IgD, IgE, IgG, IgM, and None, and each row represents a sample. For the row side color at the left of heatmap, the leftmost one was used to mark an individual while the right one was used to distinguish a project. Note: None means those clones cannot be aligned to a C gene.

**Figure S9. Clustering of V (a), D (b), and J (c) gene usage with different isotype in the male.** IgA was labeled in light blue, IgG was filled in blue, IgD was colored in light green, and IgM was labeled in green.

**Figure S10. Clustering of V (a), D (b), and J (c) gene usage with different isotype in female.** IgA was labeled in light blue, IgG was filled in blue, IgD was colored in light green, and IgM was labeled in green.

**Figure S11. Linear model for describing the correlation between number of public clones and product of numbers of clones with each sample pair.** Only sample pairs with both clone number being greater than (a) 10,000, (b) 100,000, (c)1,000,000, (d) 2,000,000, (e) 3,000,000, (f) 4,000,000 and (g) 5,000,000 were selected to demonstrate the correlation. The regression functions are at the top of figures. Selected sample pair that with more clones show more fitness of the linear model.

**Figure S12. Public clone gene segment usage.** (a, b) The two barplots show V and J gene usage frequency between public and private clones. Gene segments have been sorted by overall frequency and for v gene only those comprised more than 1% of the total repertoire were listed here. (c, d) The scatter plot demonstrated gene usage frequency correlation between public and private clones. The top left value (ρ) indicates Pearson’s correlation coefficient.

**Figure S13. CDR3 nucleotide length distribution comparison between total productive clones (n=267,761,654) and unique public clones (n=429,157).** Note that the upper limit for length is determined according to a threshold of 1%. Two-sample Kolmogorov-Smirnov tests were performed to investigate length distribution difference (P-value <1.149e-13).

**Figure S14. Junctional modification comparison between total clones from 2,165 samples and public clones.** The boxplot in each subfigure demonstrates the distribution of each kind of junction modification length, as indicated by the schematic in bottom right (a). (b) Non-templated insertion length. (c) Palindromic insertion length distributions. (d) Deletion length distributions.

**Figure S15. Percent of clones with D hit(s) for 2,165 samples and all public clones.** The boxplot on the top demonstrates the percent distribution for 2,165 samples (with a median of 97.9%), and the red point at the bottom indicates the percent for public clones (67.6%).

**Figure S16. Antigen-specific or disease-associated annotation of public clones.** (a) Overlapping of unique clonotypes between public clones and antibody sequences curated with related antigen or disease information. A clonotype here was defined as a unique CDR3 amino acid sequence with deprecated conserved residuals at both ends. (b) Disease or antigen percentage of annotated public clones. Terms in the same line in the legend are indicated by same color. Terms in legend match the pie chart from top to bottom and from left to right.

**Figure S17. Somatic hypermutation patterns and influence factors.** (a) and (b) represent the transform among nucleotides based on the algorithm of consensus and position weight matrix (PWM). Each column and each row represent a nucleotide. The germline nucleotide is located on the x axis and the mutated nucleotide is located on the Y axis. The total mutation rate and target preference of every nucleotide are marked in the figure. (c), (d) and (e) used position weight matrix (PWM) to describe the patterns of somatic hypermutation.

**Figure S18. Description of the 290th position, which has the highest mutation rate.** (a), (b), (c), and (d) show the composition of nucleotide, amino acid, codon, and motif in the germline sequence sorted by ratio, respectively. (e), (f) and (g) showed the fraction and composition of mutation from different families. (h) The boxplot shows the comparison of mutation rate between synonymous mutations and nonsynonymous mutations from different families.

**Figure S19. Mutation rate comparison between male and female based on IgG clones.** (a-e) were based on consensus approach and (f-k) were based on PWM approach. Comparison were performed independently in different age groups to remove age-related effect (marked on the top right of each subfigure). and only those age groups with at least 10 donors for both genders were presented here. The four numbers following each isotype in figure legends represent the number of clones, samples, donors and projects, respectively.

**Figure S20. Mutation rate comparison between different isotypes.** (a-c) were based on consensus approach and (d-f) were based on PWM approach. Comparison were performed independently in different age groups to remove age-related effect (marked on the top right of each subfigure). Only 3 age groups have two or more kinds of clones. The four numbers following each isotype show the number of clones, samples, donors and projects, respectively.

## DATA AVAILABILITY

In-house generated datasets are available at the NCBI Sequencing Read Archive (www.ncbi.nlm.nih.gov/sra) under BioProject number PRJNA564936. A table linking dataset accessions to their corresponding sample ids was provided in Table S9.

## References

Bonsignori, M., Zhou, T., Sheng, Z., Chen, L., Gao, F., Joyce, M.G., Ozorowski, G., Chuang, G.Y., Schramm, C.A., and Wiehe, K., et al. (2016). Maturation Pathway from Germline to Broad HIV-1 Neutralizer of a CD4-Mimic Antibody. CELL 165, 449–463.

Boyd, S.D., Gaeta, B.A., Jackson, K.J., Fire, A.Z., Marshall, E.L., Merker, J.D., Maniar, J.M., Zhang, L.N., Sahaf, B., and Jones, C.D., et al. (2010). Individual variation in the germline Ig gene repertoire inferred from variable region gene rearrangements. J IMMUNOL 184, 6986–6992.

Briney, B., Inderbitzin, A., Joyce, C., and Burton, D.R. (2019). Commonality despite exceptional diversity in the baseline human antibody repertoire. NATURE 566, 393–397.

Briney, B.S., Willis, J.R., Hicar, M.D., Thomas, J.W., and Crowe, J.E. (2012). Frequency and genetic characterization of V(DD)J recombinants in the human peripheral blood antibody repertoire. IMMUNOLOGY 137, 56–64.

Briney, B.S., Willis, J.R., McKinney, B.A., and Crowe, J.J. (2012). High-throughput antibody sequencing reveals genetic evidence of global regulation of the naive and memory repertoires that extends across individuals. GENES IMMUN 13, 469–473.

Bürckert, J., Dubois, A.R.S.X., Faison, W.J., Farinelle, S., Charpentier, E., Sinner, R., Wienecke-Baldacchino, A., and Muller, C.P. (2017). Functionally Convergent B Cell Receptor Sequences in Transgenic Rats Expressing a Human B Cell Repertoire in Response to Tetanus Toxoid and Measles Antigens. FRONT IMMUNOL 8.

Chothia, C., Lesk, A.M., Tramontano, A., Levitt, M., Smith-Gill, S.J., Air, G., Sheriff, S., Padlan, E.A., Davies, D., and Tulip, W.R., et al. (1989). Conformations of immunoglobulin hypervariable regions. NATURE 342, 877–883.

Early, P., Huang, H., Davis, M., Calame, K., and Hood, L. (1980). An immunoglobulin heavy chain variable region gene is generated from three segments of DNA: VH, D and JH. CELL 19, 981–992.

Faham, M., Zheng, J., Moorhead, M., Carlton, V.E., Stow, P., Coustan-Smith, E., Pui, C.H., and Campana, D. (2012). Deep-sequencing approach for minimal residual disease detection in acute lymphoblastic leukemia. BLOOD 120, 5173–5180.

Gawad, C., Pepin, F., Carlton, V.E.H., Klinger, M., Logan, A.C., Miklos, D.B., Faham, M., Dahl, G., and Lacayo, N. (2012). Massive evolution of the immunoglobulin heavy chain locus in children with B precursor acute lymphoblastic leukemia. BLOOD 120, 4407–4417.

Glanville, J., Kuo, T.C., von Budingen, H.C., Guey, L., Berka, J., Sundar, P.D., Huerta, G., Mehta, G.R., Oksenberg, J.R., and Hauser, S.L., et al. (2011). Naive antibody gene-segment frequencies are heritable and unaltered by chronic lymphocyte ablation. Proceedings of the National Academy of Sciences 108, 20066–20071.

Greiff, V., Miho, E., Menzel, U., and Reddy, S.T. (2015). Bioinformatic and Statistical Analysis of Adaptive Immune Repertoires. TRENDS IMMUNOL 36, 738–749.

Greiff, V., Weber, C.R., Palme, J., Bodenhofer, U., Miho, E., Menzel, U., and Reddy, S.T. (2017). Learning the High-Dimensional Immunogenomic Features That Predict Public and Private Antibody Repertoires. J IMMUNOL 199, 2985–2997.

Hansen, T.O., Lange, A.B., and Barington, T. (2015). Sterile DJH rearrangements reveal that distance between gene segments on the human Ig H chain locus influences their ability to rearrange. J IMMUNOL 194, 973–982.

He, L., Sok, D., Azadnia, P., Hsueh, J., Landais, E., Simek, M., Koff, W.C., Poignard, P., Burton, D.R., and Zhu, J. (2015). Toward a more accurate view of human B-cell repertoire by next-generation sequencing, unbiased repertoire capture and single-molecule barcoding. SCI REP-UK 4.

Hoh, R.A., Joshi, S.A., Liu, Y., Wang, C., Roskin, K.M., Lee, J., Pham, T., Looney, T.J., Jackson, K.J.L., and Dixit, V.P., et al. (2016). Single B-cell deconvolution of peanut-specific antibody responses in allergic patients. J ALLERGY CLIN IMMUN 137, 157–167.

Hong, B., Wu, Y., Li, W., Wang, X., Wen, Y., Jiang, S., Dimitrov, D.S., and Ying, T. (2018). In-Depth Analysis of Human Neonatal and Adult IgM Antibody Repertoires. FRONT IMMUNOL 9, 128.

Jackson, K.J., Kidd, M.J., Wang, Y., and Collins, A.M. (2013). The shape of the lymphocyte receptor repertoire: lessons from the B cell receptor. FRONT IMMUNOL 4, 263.

Jackson, K.J., Liu, Y., Roskin, K.M., Glanville, J., Hoh, R.A., Seo, K., Marshall, E.L., Gurley, T.C., Moody, M.A., and Haynes, B.F., et al. (2014). Human responses to influenza vaccination show seroconversion signatures and convergent antibody rearrangements. CELL HOST MICROBE 16, 105–114.

Jiang, N., He, J., Weinstein, J.A., Penland, L., Sasaki, S., He, X.S., Dekker, C.L., Zheng, N.Y., Huang, M., and Sullivan, M., *et* al. (2013). Lineage structure of the human antibody repertoire in response to influenza vaccination. SCI TRANSL MED 5, 119r–171r.

Joyce, M.G., Wheatley, A.K., Thomas, P.V., Chuang, G., Soto, C., Bailer, R.T., Druz, A., Georgiev, I.S., Gillespie, R.A., and Kanekiyo, M., et al. (2016). Vaccine-Induced Antibodies that Neutralize Group 1 and Group 2 Influenza A Viruses. CELL 166, 609–623.

Kidd, M.J., Jackson, K.J., Boyd, S.D., and Collins, A.M. (2016). DJ Pairing during VDJ Recombination Shows Positional Biases That Vary among Individuals with Differing IGHD Locus Immunogenotypes. J IMMUNOL 196, 1158–1164.

Kitaura, K., Yamashita, H., Ayabe, H., Shini, T., Matsutani, T., and Suzuki, R. (2017). Different Somatic Hypermutation Levels among Antibody Subclasses Disclosed by a New Next-Generation Sequencing-Based Antibody Repertoire Analysis. FRONT IMMUNOL 8.

Krebs, S.J., Kwon, Y.D., Schramm, C.A., Law, W.H., Donofrio, G., Zhou, K.H., Gift, S., Dussupt, V., Georgiev, I.S., and Schätzle, S., et al. (2019). Longitudinal Analysis Reveals Early Development of Three MPER-Directed Neutralizing Antibody Lineages from an HIV-1-Infected Individual. IMMUNITY 50, 677–691.

Kurtz, D.M., Green, M.R., Bratman, S.V., Scherer, F., Liu, C.L., Kunder, C.A., Takahashi, K., Glover, C., Keane, C., and Kihira, S., et al. (2015). Noninvasive monitoring of diffuse large B-cell lymphoma by immunoglobulin high-throughput sequencing. BLOOD 125, 3679–3687.

Larimore, K., McCormick, M.W., Robins, H.S., and Greenberg, P.D. (2012). Shaping of human germline IgH repertoires revealed by deep sequencing. J IMMUNOL 189, 3221–3230.

Laserson, U., Vigneault, F., Gadala-Maria, D., Yaari, G., Uduman, M., Vander, H.J., Kelton, W., Taek, J.S., Liu, Y., and Laserson, J., et al. (2014). High-resolution antibody dynamics of vaccine-induced immune responses. Proc Natl Acad Sci U S A 111, 4928–4933.

Li, G.M., Chiu, C., Wrammert, J., McCausland, M., Andrews, S.F., Zheng, N.Y., Lee, J.H., Huang, M., Qu, X., and Edupuganti, S., et al. (2012). Pandemic H1N1 influenza vaccine induces a recall response in humans that favors broadly cross-reactive memory B cells. Proceedings of the National Academy of Sciences 109, 9047–9052.

Liu, M., and Schatz, D.G. (2009). Balancing AID and DNA repair during somatic hypermutation. TRENDS IMMUNOL 30, 173–181.

Liu, X., Zhang, W., Zeng, X., Zhang, R., Du, Y., Hong, X., Cao, H., Su, Z., Wang, C., and Wu, J., et al. (2016). Systematic Comparative Evaluation of Methods for Investigating the TCRβ Repertoire. PLOS ONE 11, e152464.

Madi, A., Shifrut, E., Reich-Zeliger, S., Gal, H., Best, K., Ndifon, W., Chain, B., Cohen, I.R., and Friedman, N. (2014). T-cell receptor repertoires share a restricted set of public and abundant CDR3 sequences that are associated with self-related immunity. GENOME RES 24, 1603–1612.

Maecker, H.T., Lindstrom, T.M., Robinson, W.H., Utz, P.J., Hale, M., Boyd, S.D., Shen-Orr, S.S., and Fathman, C.G. (2012). New tools for classification and monitoring of autoimmune diseases. NAT REV RHEUMATOL 8, 317–328.

Miho, E., Roskar, R., Greiff, V., and Reddy, S.T. (2019). Large-scale network analysis reveals the sequence space architecture of antibody repertoires. NAT COMMUN 10, 1321.

Parameswaran, P., Liu, Y., Roskin, K.M., Jackson, K.K.L., Dixit, V.P., Lee, J., Artiles, K.L., Zompi, S., Vargas, M.J., and Simen, B.B., et al. (2013). Convergent Antibody Signatures in Human Dengue. CELL HOST MICROBE 13, 691–700.

Patil, S.U., Ogunniyi, A.O., Calatroni, A., Tadigotla, V.R., Ruiter, B., Ma, A., Moon, J., Love, J.C., and Shreffler, W.G. (2015). Peanut oral immunotherapy transiently expands circulating Ara h 2–specific B cells with a homologous repertoire in unrelated subjects. J ALLERGY CLIN IMMUN 136, 125–134.

Pham, P., Bransteitter, R., Petruska, J., and Goodman, M.F. (2003). Processive AID-catalysed cytosine deamination on single-stranded DNA simulates somatic hypermutation. NATURE 424, 103–107.

Pugh-Bernard, A.E., Silverman, G.J., Cappione, A.J., Villano, M.E., Ryan, D.H., Insel, R.A., and Sanz, I. (2001). Regulation of inherently autoreactive VH4-34 B cells in the maintenance of human B cell tolerance. J CLIN INVEST 108, 1061–1070.

Reddy, S.T., Ge, X., Miklos, A.E., Hughes, R.A., Kang, S.H., Hoi, K.H., Chrysostomou, C., Hunicke-Smith, S.P., Iverson, B.L., and Tucker, P.W., et al. (2010). Monoclonal antibodies isolated without screening by analyzing the variable-gene repertoire of plasma cells. NAT BIOTECHNOL 28, 965–969.

Robins, H. (2013). Immunosequencing: applications of immune repertoire deep sequencing. CURR OPIN IMMUNOL 25, 646–652.

Saada, R., Weinberger, M., Shahaf, G., and Mehr, R. (2007). Models for antigen receptor gene rearrangement: CDR3 length. IMMUNOL CELL BIOL 85, 323–332.

Safonova, Y., and Pevzner, P.A. (2019). De novo Inference of Diversity Genes and Analysis of Non-canonical V(DD)J Recombination in Immunoglobulins. FRONT IMMUNOL 10, 987.

Schramm, C.A., and Douek, D.C. (2018). Beyond Hot Spots: Biases in Antibody Somatic Hypermutation and Implications for Vaccine Design. FRONT IMMUNOL 9, 1876.

Setliff, I., McDonnell, W.J., Raju, N., Bombardi, R.G., Murji, A.A., Scheepers, C., Ziki, R., Mynhardt, C., Shepherd, B.E., and Mamchak, A.A., et al. (2018). Multi-Donor Longitudinal Antibody Repertoire Sequencing Reveals the Existence of Public Antibody Clonotypes in HIV-1 Infection. CELL HOST MICROBE 23, 845–854.

Shapiro, G.S., Aviszus, K., Ikle, D., and Wysocki, L.J. (1999). Predicting regional mutability in antibody V genes based solely on di- and trinucleotide sequence composition. J IMMUNOL 163, 259–268.

Soto, C., Bombardi, R.G., Branchizio, A., Kose, N., Matta, P., Sevy, A.M., Sinkovits, R.S., Gilchuk, P., Finn, J.A., and Crowe, J.E. (2019). High frequency of shared clonotypes in human B cell receptor repertoires. NATURE.

Souto-Carneiro, M.M., Sims, G.P., Girschik, H., Lee, J., and Lipsky, P.E. (2005). Developmental Changes in the Human Heavy Chain CDR3. The Journal of Immunology 175, 7425–7436.

Stern, J.N.H., Yaari, G., Vander Heiden, J., Church, G., Donahue, W.F., Hintzen, R., Huttner, A.J., Laman, J., Nagra, R.M., and Nylander, A., et al. (2014). B cells populating the multiple sclerosis brain mature in the draining cervical lymph nodes. SCI TRANSL MED 6, 107r–248r.

Sui, J., Hwang, W.C., Perez, S., Wei, G., Aird, D., Chen, L., Santelli, E., Stec, B., Cadwell, G., and Ali, M., et al. (2009). Structural and functional bases for broad-spectrum neutralization of avian and human influenza A viruses. NAT STRUCT MOL BIOL 16, 265–273.

Tipton, C.M., Fucile, C.F., Darce, J., Chida, A., Ichikawa, T., Gregoretti, I., Schieferl, S., Hom, J., Jenks, S., and Feldman, R.J., et al. (2015). Diversity, cellular origin and autoreactivity of antibody-secreting cell population expansions in acute systemic lupus erythematosus. NAT IMMUNOL 16, 755–765.

Tonegawa, S. (1983). Somatic generation of antibody diversity. NATURE 302, 575–581.

Truck, J., Ramasamy, M.N., Galson, J.D., Rance, R., Parkhill, J., Lunter, G., Pollard, A.J., and Kelly, D.F. (2015). Identification of antigen-specific B cell receptor sequences using public repertoire analysis. J IMMUNOL 194, 252–261.

von Büdingen, H., Kuo, T.C., Sirota, M., van Belle, C.J., Apeltsin, L., Glanville, J., Cree, B.A., Gourraud, P., Schwartzburg, A., and Huerta, G., et al. (2012). B cell exchange across the blood-brain barrier in multiple sclerosis. The Journal of clinical investigation 122, 4533–4543.

Wang, C., Liu, Y., Xu, L.T., Jackson, K.J., Roskin, K.M., Pham, T.D., Laserson, J., Marshall, E.L., Seo, K., and Lee, J.Y., et al. (2014). Effects of aging, cytomegalovirus infection, and EBV infection on human B cell repertoires. J IMMUNOL 192, 603–611.

Weinstein, J.A., Jiang, N., White, R.A., Fisher, D.S., and Quake, S.R. (2009). High-Throughput Sequencing of the Zebrafish Antibody Repertoire. SCIENCE 324, 807–810.

Wu, X., Zhang, Z., Schramm, C.A., Joyce, M.G., Do Kwon, Y., Zhou, T., Sheng, Z., Zhang, B., O Dell, S., and McKee, K., et al. (2015). Maturation and Diversity of the VRC01-Antibody Lineage over 15 Years of Chronic HIV-1 Infection. CELL 161, 470–485.

Wu, X., Zhou, T., Zhu, J., Zhang, B., Georgiev, I., Wang, C., Chen, X., Longo, N.S., Louder, M., and McKee, K., et al. (2011a). Focused Evolution of HIV-1 Neutralizing Antibodies Revealed by Structures and Deep Sequencing. SCIENCE 333, 1593–1602.

Wu, X., Zhou, T., Zhu, J., Zhang, B., Georgiev, I., Wang, C., Chen, X., Longo, N.S., Louder, M., and McKee, K., et al. (2011b). Focused Evolution of HIV-1 Neutralizing Antibodies Revealed by Structures and Deep Sequencing. SCIENCE 333, 1593–1602.

Wu, Y.B., James, L.K., Vander Heiden, J.A., Uduman, M., Durham, S.R., Kleinstein, S.H., Kipling, D., and Gould, H.J. (2014). Influence of seasonal exposure to grass pollen on local and peripheral blood IgE repertoires in patients with allergic rhinitis. J ALLERGY CLIN IMMUN 134, 604–612.

Zhu, J., Wu, X., Zhang, B., McKee, K., O’Dell, S., Soto, C., Zhou, T., Casazza, J.P., Mullikin, J.C., and Kwong, P.D., et al. (2013a). De novo identification of VRC01 class HIV-1-neutralizing antibodies by next-generation sequencing of B-cell transcripts. Proceedings of the National Academy of Sciences 110, E4088–E4097.

Zhu, J., Wu, X., Zhang, B., McKee, K., O’Dell, S., Soto, C., Zhou, T., Casazza, J.P., Mullikin, J.C., and Kwong, P.D., et al. (2013b). De novo identification of VRC01 class HIV-1-neutralizing antibodies by next-generation sequencing of B-cell transcripts. Proceedings of the National Academy of Sciences 110, E4088–E4097.

