## Supplementary figures for "Large-scale Analysis of 2,152 dataset reveals key features of B cell biology and the antibody repertoire"

### Slide 1
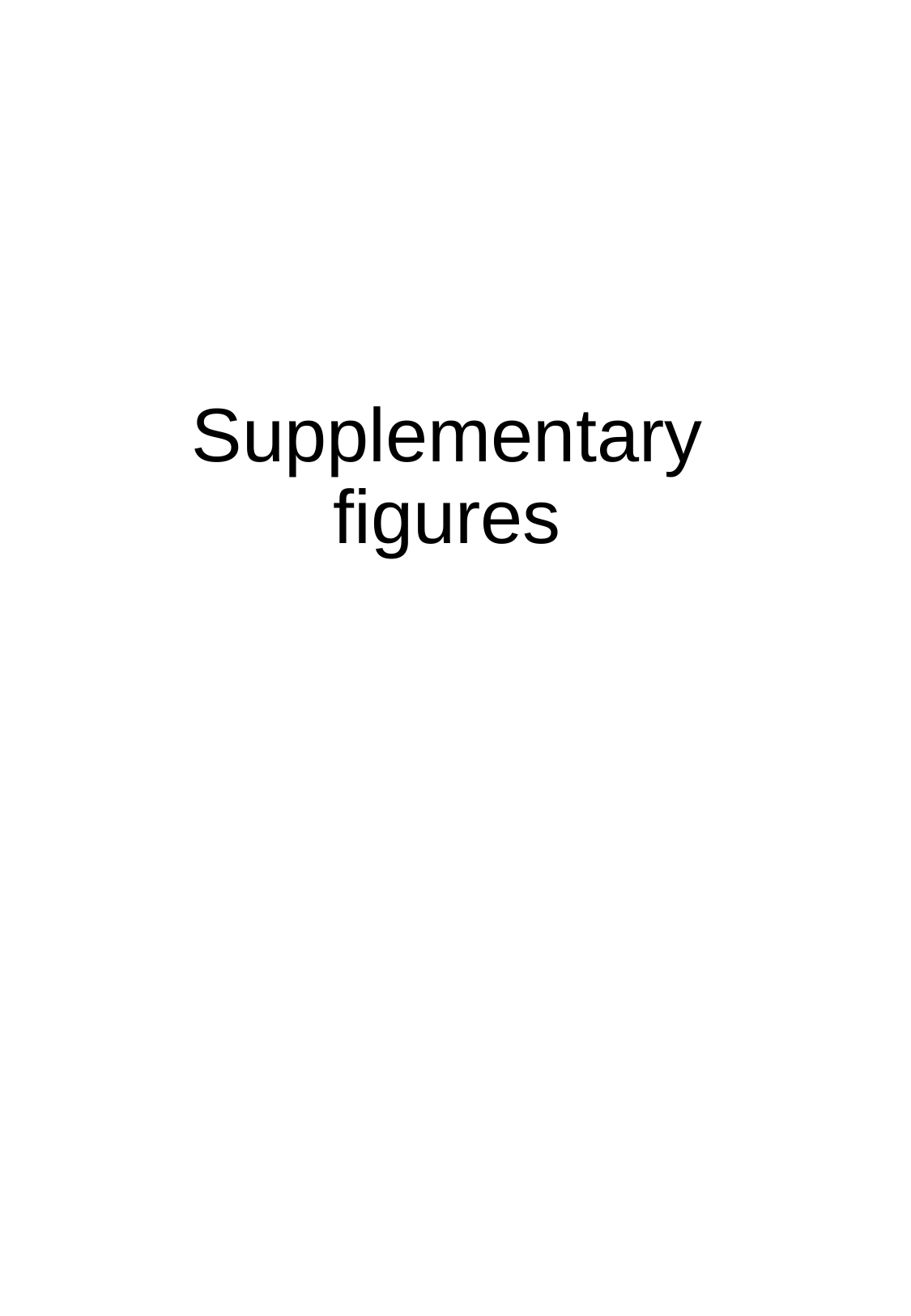

Supplementary figures

### Slide 2
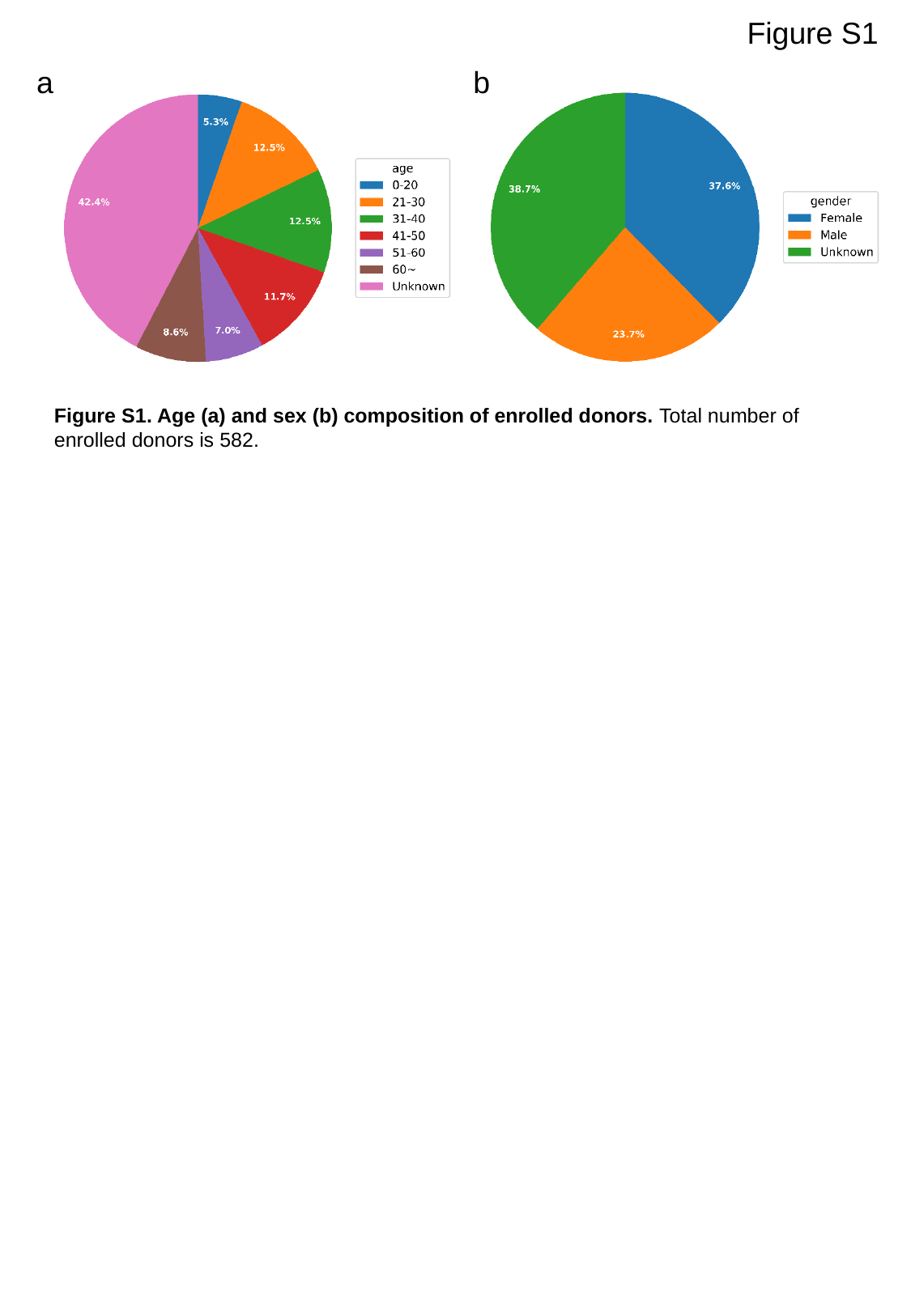

Figure S1
a
b
Figure S1. Age (a) and sex (b) composition of enrolled donors. Total number of enrolled donors is 582.

### Slide 3
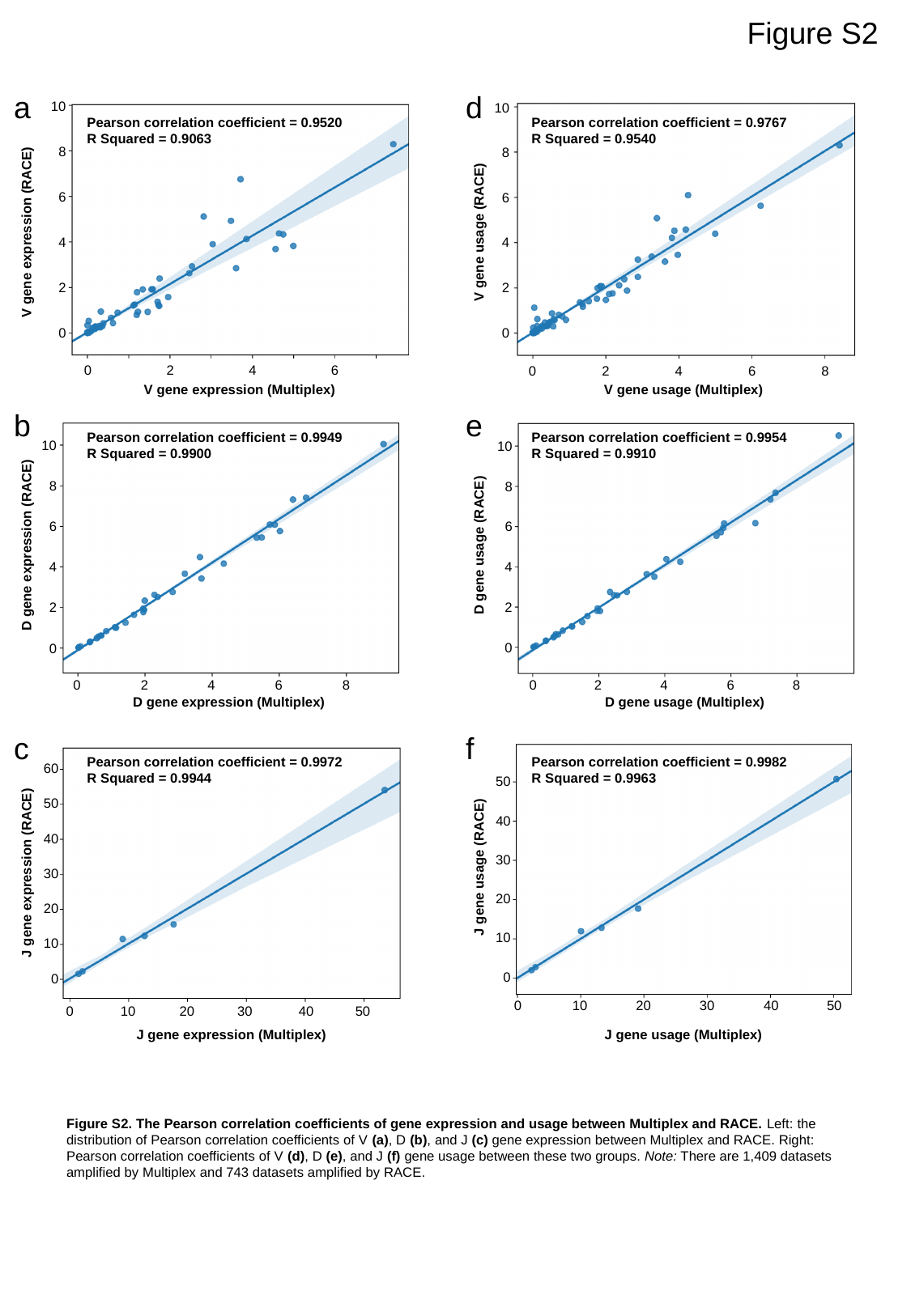

Figure S2
a
d
10
8
6
4
2
0
10
8
6
4
2
0
Pearson correlation coefficient = 0.9520
R Squared = 0.9063
Pearson correlation coefficient = 0.9767
R Squared = 0.9540
V gene usage (RACE)
V gene expression (RACE)
0
2
4
6
0
2
4
6
8
V gene expression (Multiplex)
V gene usage (Multiplex)
b
e
Pearson correlation coefficient = 0.9949
R Squared = 0.9900
Pearson correlation coefficient = 0.9954
R Squared = 0.9910
10
8
6
4
2
0
10
8
6
D gene expression (RACE)
D gene usage (RACE)
4
2
0
0
2
4
6
8
0
2
4
6
8
D gene expression (Multiplex)
D gene usage (Multiplex)
c
f
Pearson correlation coefficient = 0.9972
R Squared = 0.9944
Pearson correlation coefficient = 0.9982
R Squared = 0.9963
60
50
40
30
20
10
0
50
40
J gene usage (RACE)
J gene expression (RACE)
30
20
10
0
0
10
20
30
40
50
0
10
20
30
40
50
J gene expression (Multiplex)
J gene usage (Multiplex)
Figure S2. The Pearson correlation coefficients of gene expression and usage between Multiplex and RACE. Left: the distribution of Pearson correlation coefficients of V (a), D (b), and J (c) gene expression between Multiplex and RACE. Right: Pearson correlation coefficients of V (d), D (e), and J (f) gene usage between these two groups. Note: There are 1,409 datasets amplified by Multiplex and 743 datasets amplified by RACE.

### Slide 4
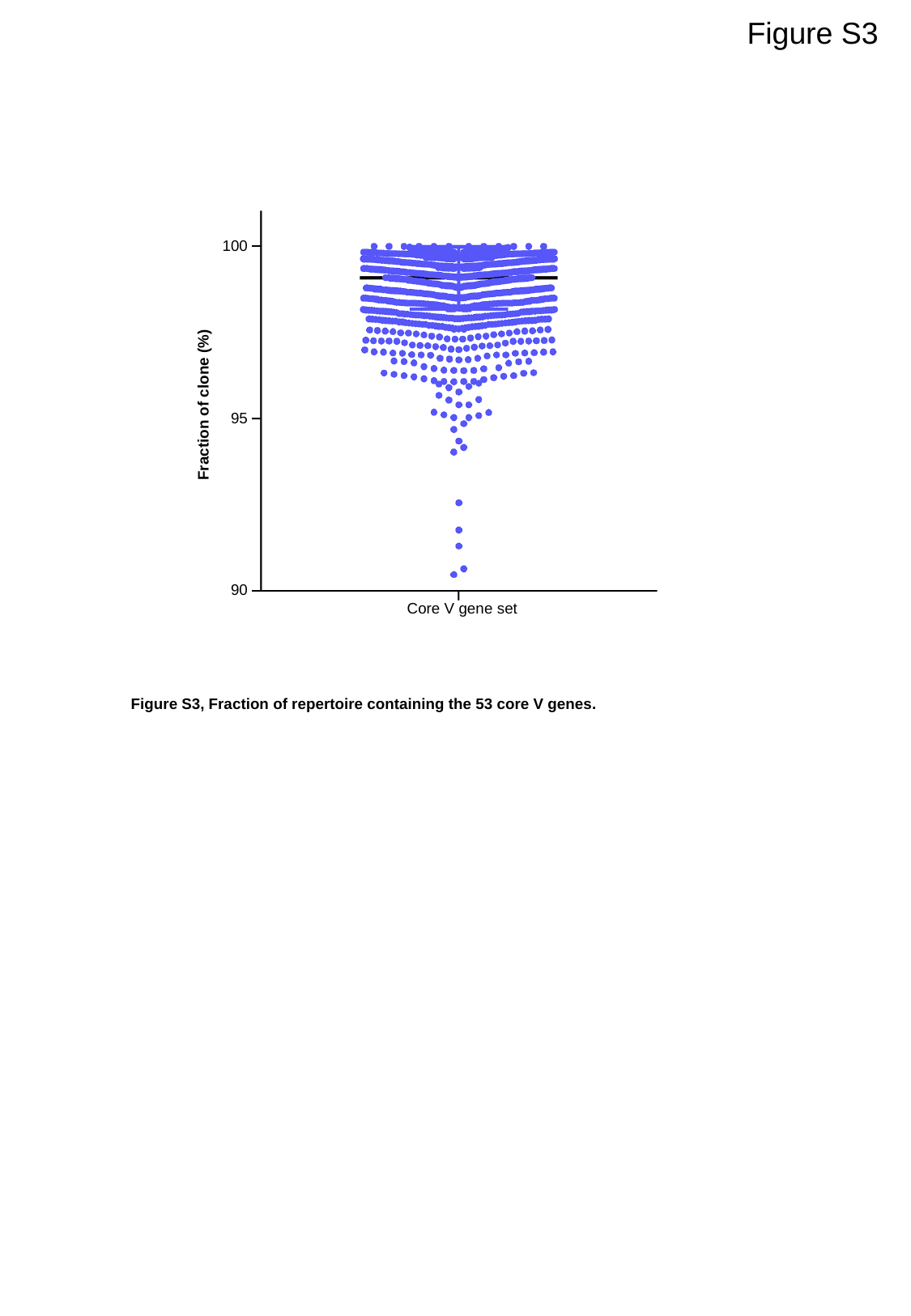

Figure S3
100
95
90
Fraction of clone (%)
Core V gene set
Figure S3, Fraction of repertoire containing the 53 core V genes.

### Slide 5
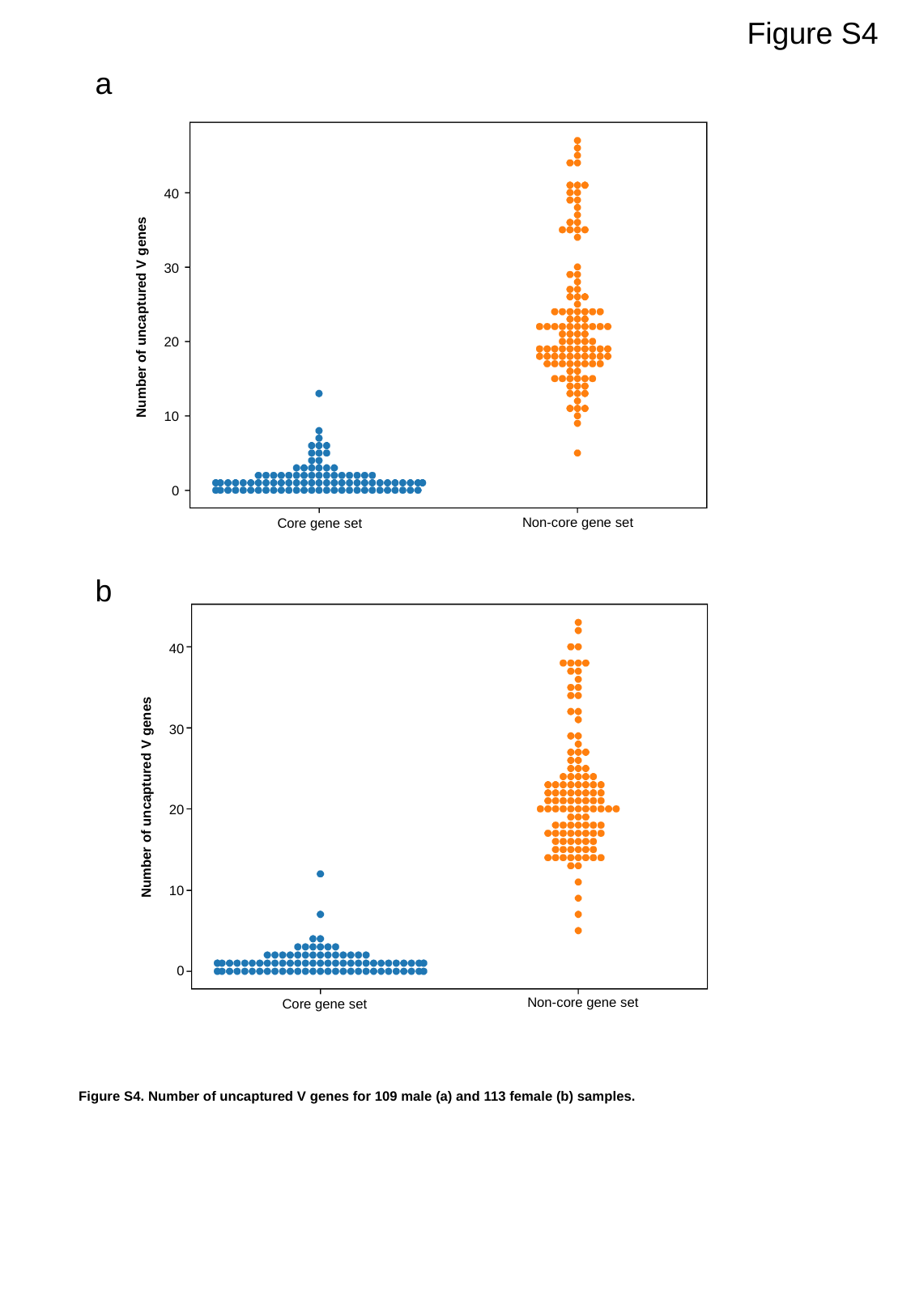

Figure S4
a
40
30
20
10
0
Number of uncaptured V genes
Non-core gene set
Core gene set
b
40
30
20
10
0
Number of uncaptured V genes
Non-core gene set
Core gene set
Figure S4. Number of uncaptured V genes for 109 male (a) and 113 female (b) samples.

### Slide 6
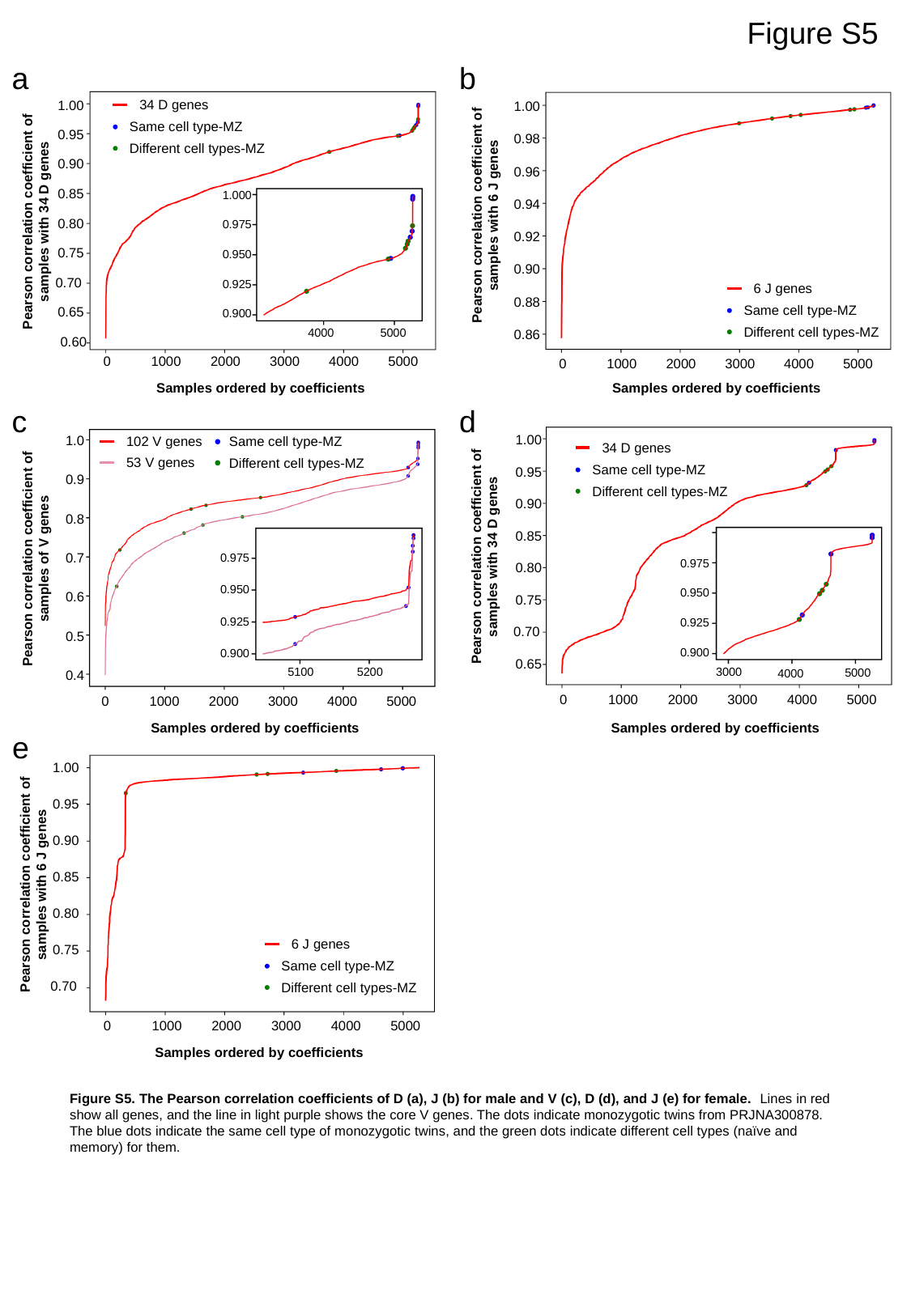

Figure S5
a
b
34 D genes
Same cell type-MZ
Different cell types-MZ
1.00
0.95
0.90
0.85
0.80
0.75
0.70
0.65
0.60
1.00
0.98
0.96
0.94
0.92
0.90
0.88
0.86
1.000
Pearson correlation coefficient of samples with 6 J genes
Pearson correlation coefficient of samples with 34 D genes
0.975
0.950
0.925
6 J genes
Same cell type-MZ
Different cell types-MZ
0.900
5000
4000
0
1000
2000
3000
4000
5000
0
1000
2000
3000
4000
5000
Samples ordered by coefficients
Samples ordered by coefficients
c
d
1.00
0.95
0.90
0.85
0.80
0.75
0.70
0.65
1.0
0.9
0.8
0.7
0.6
0.5
0.4
Same cell type-MZ
102 V genes
53 V genes
Different cell types-MZ
34 D genes
Same cell type-MZ
Different cell types-MZ
Pearson correlation coefficient of samples with 34 D genes
Pearson correlation coefficient of samples of V genes
0.975
0.975
0.950
0.950
0.925
0.925
0.900
0.900
3000
5100
5200
5000
4000
0
1000
2000
3000
4000
5000
0
1000
2000
3000
4000
5000
Samples ordered by coefficients
Samples ordered by coefficients
e
1.00
0.95
0.90
0.85
0.80
0.75
0.70
Pearson correlation coefficient of samples with 6 J genes
6 J genes
Same cell type-MZ
Different cell types-MZ
0
1000
2000
3000
4000
5000
Samples ordered by coefficients
Figure S5. The Pearson correlation coefficients of D (a), J (b) for male and V (c), D (d), and J (e) for female. Lines in red show all genes, and the line in light purple shows the core V genes. The dots indicate monozygotic twins from PRJNA300878. The blue dots indicate the same cell type of monozygotic twins, and the green dots indicate different cell types (naïve and memory) for them.

### Slide 7
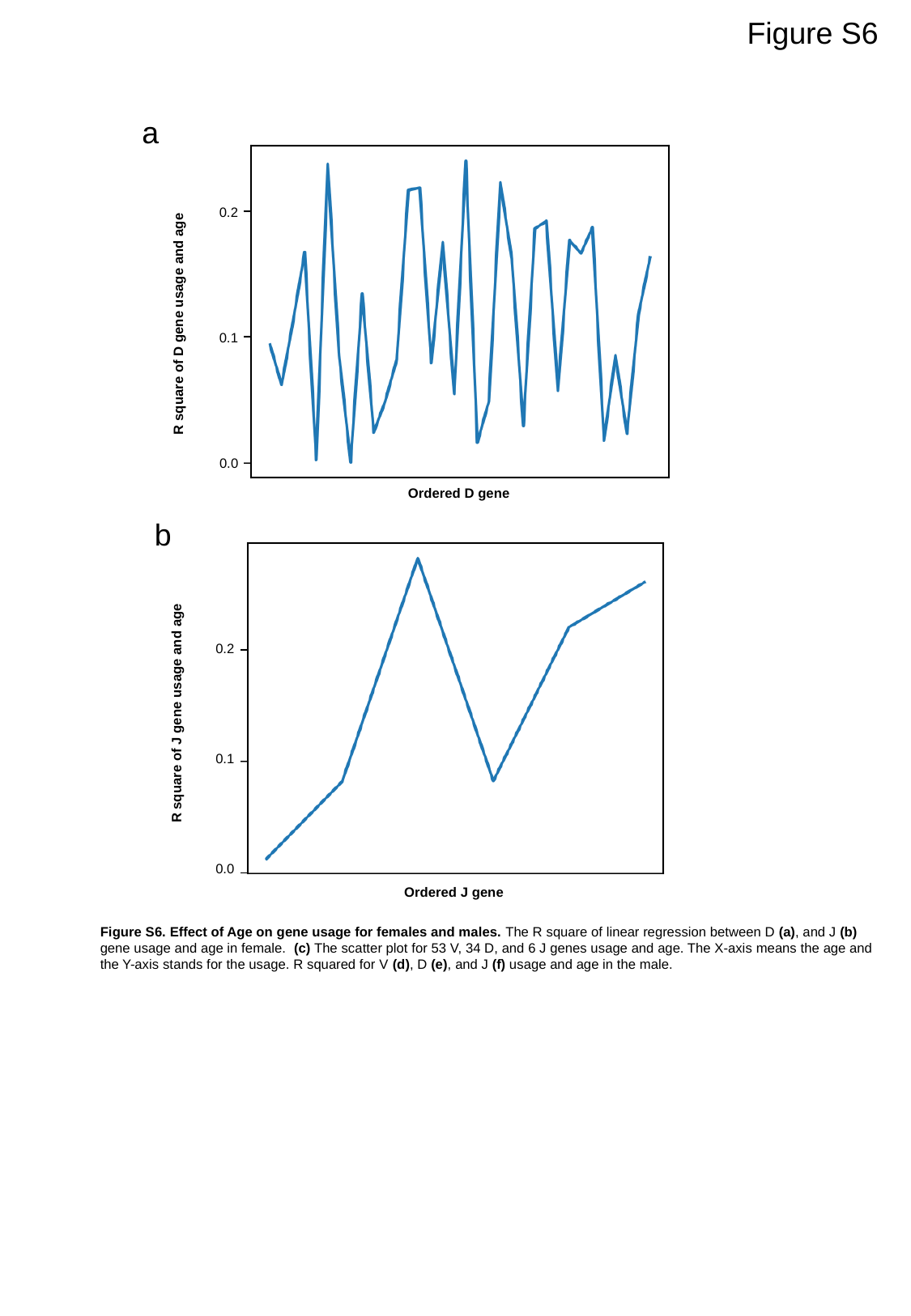

Figure S6
a
0.2
R square of D gene usage and age
0.1
0.0
Ordered D gene
b
0.2
R square of J gene usage and age
0.1
0.0
Ordered J gene
Figure S6. Effect of Age on gene usage for females and males. The R square of linear regression between D (a), and J (b) gene usage and age in female. (c) The scatter plot for 53 V, 34 D, and 6 J genes usage and age. The X-axis means the age and the Y-axis stands for the usage. R squared for V (d), D (e), and J (f) usage and age in the male.

### Slide 8
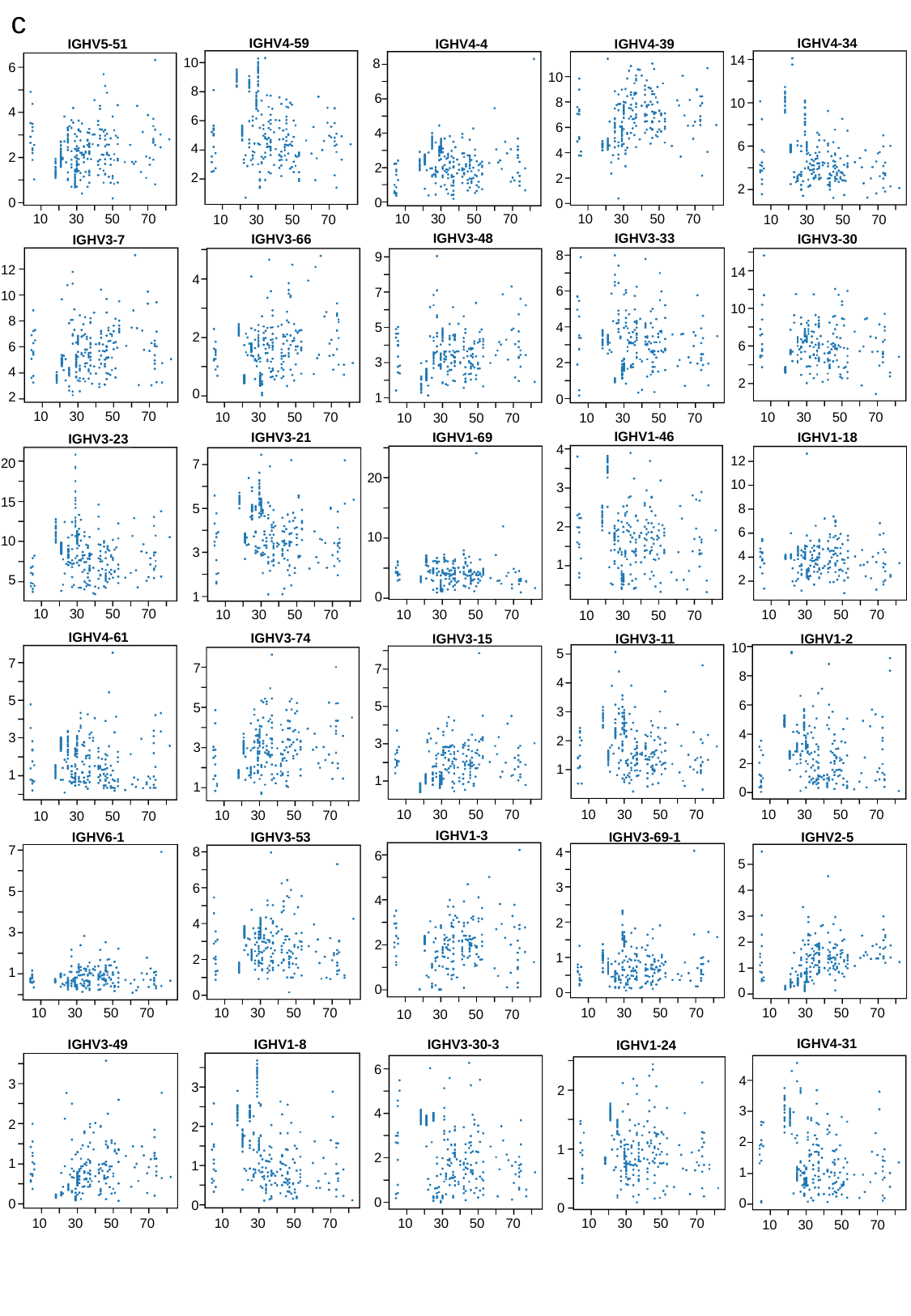

c
IGHV4-59
10
8
6
4
2
10
30
50
70
IGHV4-34
14
10
6
2
10
30
50
70
IGHV4-4
8
6
4
2
0
10
30
50
70
IGHV4-39
10
8
6
4
2
0
10
30
50
70
IGHV5-51
6
4
2
0
10
30
50
70
IGHV3-33
8
6
4
2
0
10
30
50
70
IGHV3-48
9
7
5
3
1
10
30
50
70
IGHV3-30
14
10
6
2
10
30
50
70
IGHV3-66
4
2
0
10
30
50
70
IGHV3-7
12
10
8
6
4
2
10
30
50
70
IGHV1-46
4
3
2
1
10
30
50
70
IGHV1-18
12
10
8
6
4
2
10
30
50
70
IGHV1-69
20
10
0
10
30
50
70
IGHV3-21
7
5
3
1
10
30
50
70
IGHV3-23
20
15
10
5
10
30
50
70
IGHV4-61
7
5
3
1
10
30
50
70
IGHV3-74
7
5
3
1
10
30
50
70
IGHV3-11
5
4
3
2
1
10
30
50
70
IGHV3-15
7
5
3
1
10
30
50
70
IGHV1-2
10
8
6
4
2
0
10
30
50
70
IGHV1-3
6
4
2
0
10
30
50
70
IGHV6-1
7
5
3
1
10
30
50
70
IGHV3-69-1
4
3
2
1
0
10
30
50
70
IGHV3-53
8
6
4
2
0
10
30
50
70
IGHV2-5
5
4
3
2
1
0
10
30
50
70
IGHV4-31
4
3
2
1
0
10
30
50
70
IGHV3-49
3
2
1
0
10
30
50
70
IGHV1-8
3
2
1
0
10
30
50
70
IGHV3-30-3
6
4
2
0
10
30
50
70
IGHV1-24
2
1
0
10
30
50
70

### Slide 9
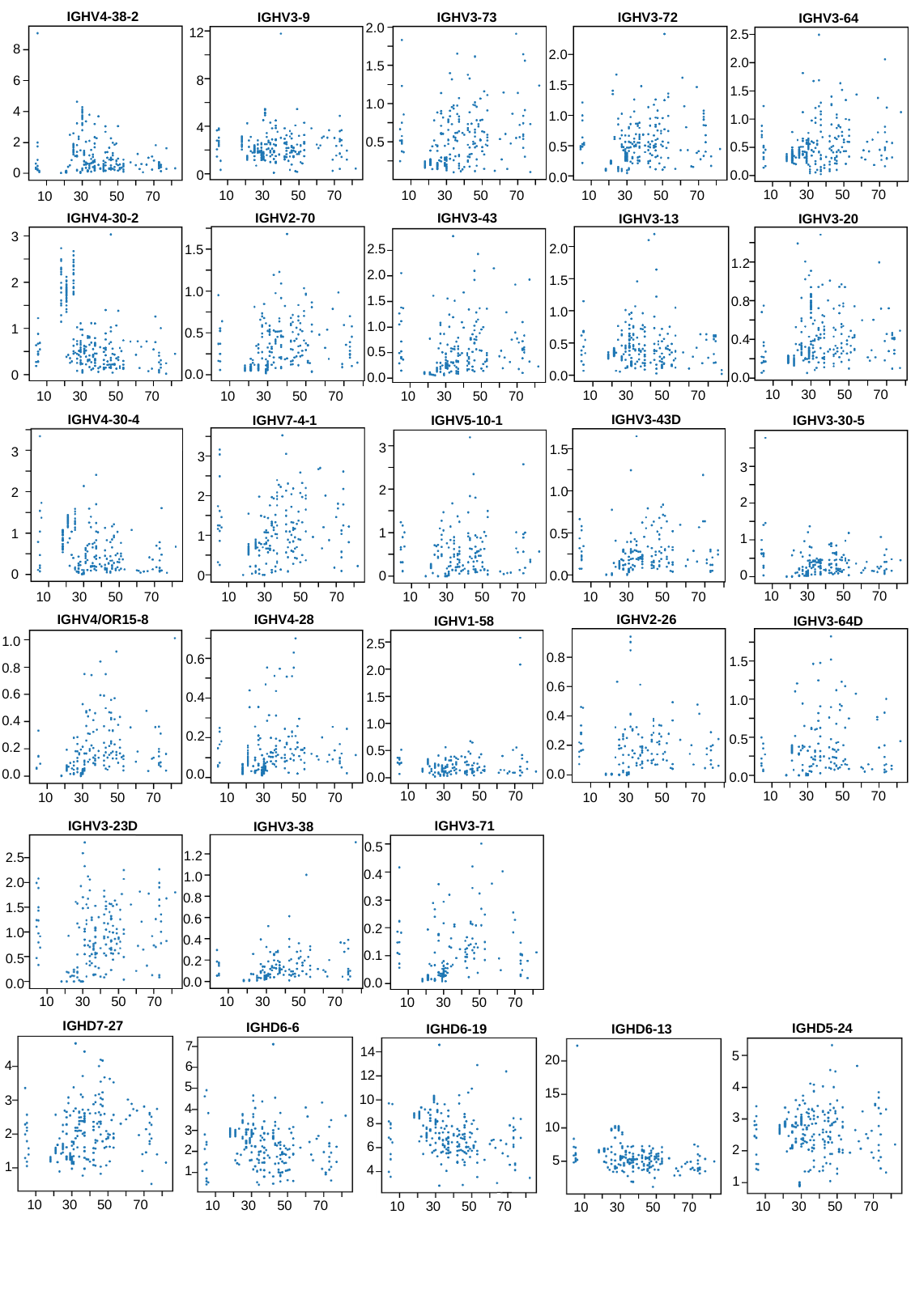

IGHV4-38-2
8
6
4
2
0
10
30
50
70
IGHV3-73
2.0
1.5
1.0
0.5
10
30
50
70
IGHV3-9
12
8
4
0
10
30
50
70
IGHV3-72
2.0
1.5
1.0
0.5
0.0
10
30
50
70
IGHV3-64
2.5
2.0
1.5
1.0
0.5
0.0
10
30
50
70
IGHV3-43
2.5
2.0
1.5
1.0
0.5
0.0
10
30
50
70
IGHV4-30-2
3
2
1
0
10
30
50
70
IGHV2-70
1.5
1.0
0.5
0.0
10
30
50
70
IGHV3-13
2.0
1.5
1.0
0.5
0.0
10
30
50
70
IGHV3-20
1.2
0.8
0.4
0.0
10
30
50
70
IGHV4-30-4
3
2
1
0
10
30
50
70
IGHV3-43D
1.5
1.0
0.5
0.0
10
30
50
70
IGHV7-4-1
3
2
1
0
10
30
50
70
IGHV5-10-1
3
2
1
0
10
30
50
70
IGHV3-30-5
3
2
1
0
10
30
50
70
IGHV4/OR15-8
1.0
0.8
0.6
0.4
0.2
0.0
10
30
50
70
IGHV4-28
0.6
0.4
0.2
0.0
10
30
50
70
IGHV2-26
0.8
0.6
0.4
0.2
0.0
10
30
50
70
IGHV1-58
2.5
2.0
1.5
1.0
0.5
0.0
10
30
50
70
IGHV3-64D
1.5
1.0
0.5
0.0
10
30
50
70
IGHV3-71
0.5
0.4
0.3
0.2
0.1
0.0
10
30
50
70
IGHV3-23D
2.5
2.0
1.5
1.0
0.5
0.0
10
30
50
70
IGHV3-38
1.2
1.0
0.8
0.6
0.4
0.2
0.0
10
30
50
70
IGHD7-27
4
3
2
1
10
30
50
70
IGHD6-6
7
6
5
4
3
2
1
10
30
50
70
IGHD5-24
5
4
3
1
2
10
30
50
70
IGHD6-19
14
12
10
8
6
4
10
30
50
70
IGHD6-13
20
15
10
5
10
30
50
70

### Slide 10
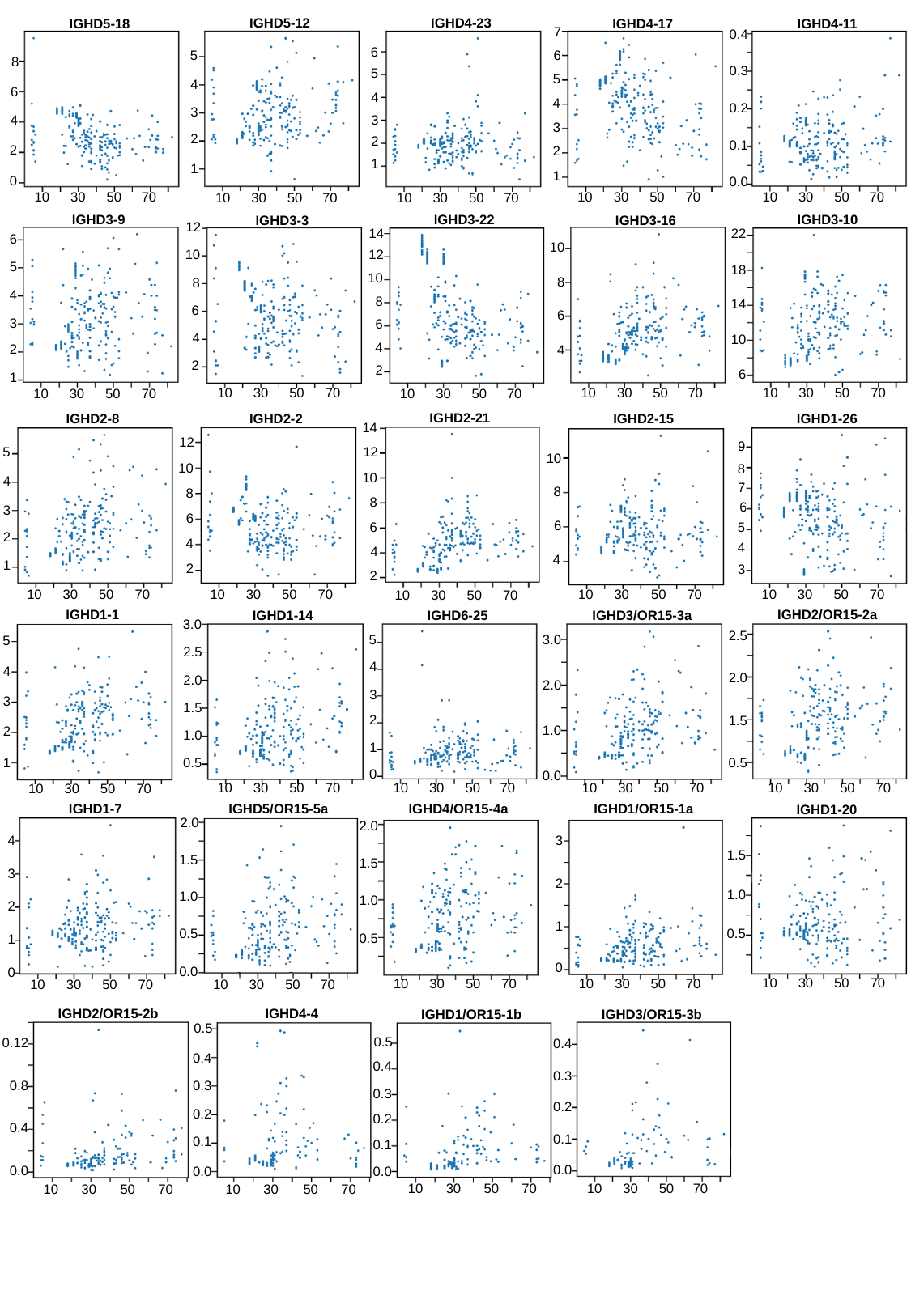

IGHD4-23
6
5
4
3
1
2
10
30
50
70
IGHD5-12
5
4
3
1
2
10
30
50
70
IGHD4-11
0.4
0.3
0.2
0.1
0.0
10
30
50
70
IGHD4-17
7
6
5
4
3
2
1
10
30
50
70
IGHD5-18
8
6
4
2
0
10
30
50
70
IGHD3-9
6
5
4
3
2
1
10
30
50
70
IGHD3-22
14
12
10
8
6
4
2
10
30
50
70
IGHD3-10
22
18
14
10
6
10
30
50
70
IGHD3-3
12
10
8
6
4
2
10
30
50
70
IGHD3-16
10
8
6
4
10
30
50
70
IGHD2-21
14
12
10
8
6
4
2
10
30
50
70
IGHD2-8
5
4
3
1
2
10
30
50
70
IGHD2-15
10
8
6
4
10
30
50
70
IGHD2-2
12
10
8
6
4
2
10
30
50
70
IGHD1-26
9
8
7
5
4
3
6
10
30
50
70
IGHD1-1
5
4
3
1
2
10
30
50
70
IGHD2/OR15-2a
2.5
2.0
1.5
0.5
10
30
50
70
IGHD1-14
3.0
2.5
2.0
1.5
1.0
0.5
10
30
50
70
IGHD6-25
5
4
3
1
0
2
10
30
50
70
IGHD3/OR15-3a
3.0
2.0
1.0
0.0
10
30
50
70
IGHD1-7
4
3
1
0
2
10
30
50
70
IGHD5/OR15-5a
2.0
1.5
0.5
0.0
1.0
10
30
50
70
IGHD4/OR15-4a
2.0
1.5
1.0
0.5
10
30
50
70
IGHD1/OR15-1a
3
2
1
0
10
30
50
70
IGHD1-20
1.5
1.0
0.5
10
30
50
70
IGHD2/OR15-2b
0.12
0.8
0.4
0.0
10
30
50
70
IGHD4-4
0.5
0.4
0.3
0.2
0.1
0.0
10
30
50
70
IGHD3/OR15-3b
0.4
0.3
0.2
0.1
0.0
10
30
50
70
IGHD1/OR15-1b
0.5
0.4
0.3
0.2
0.1
0.0
10
30
50
70

### Slide 11
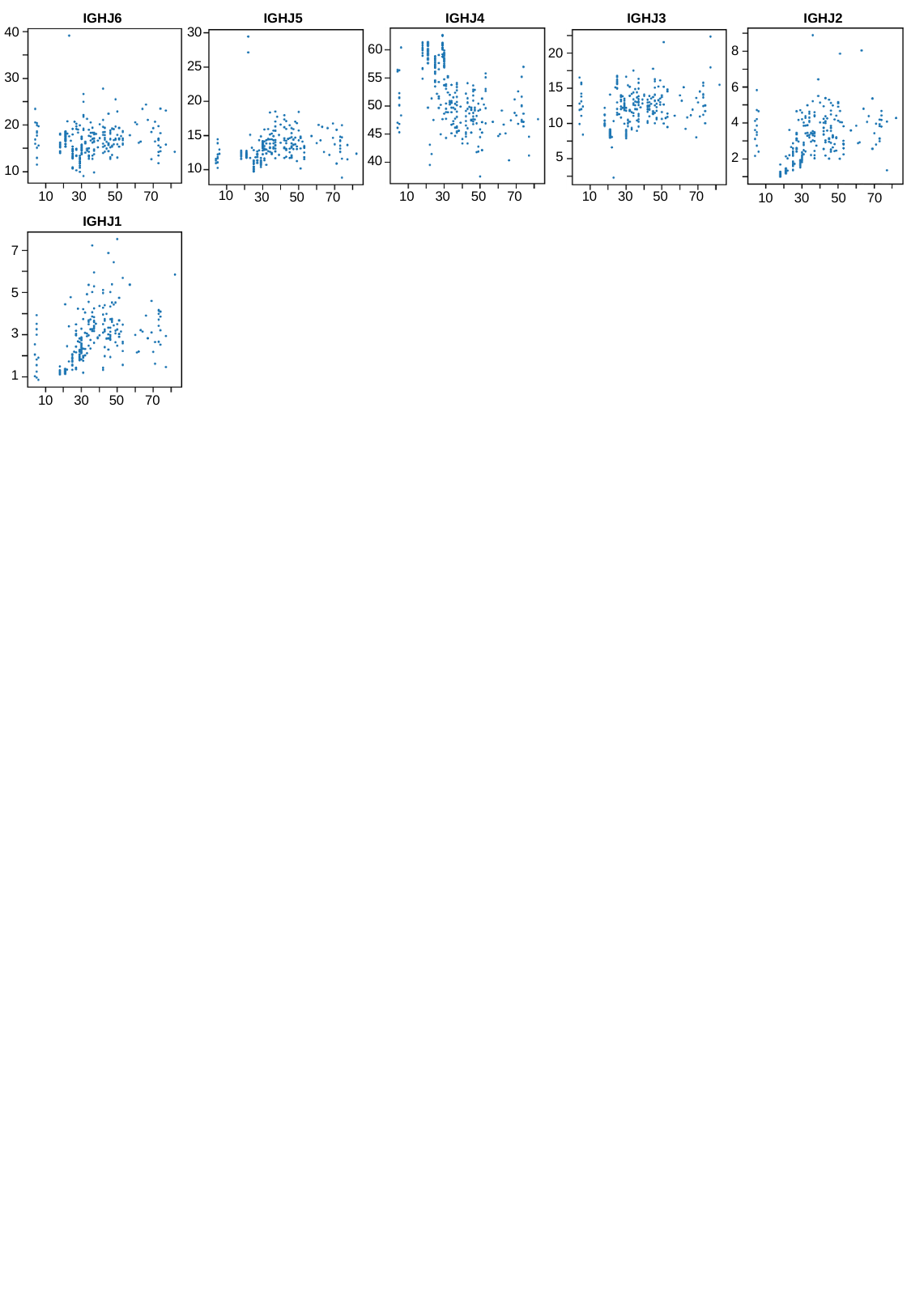

IGHJ6
40
30
20
10
10
30
50
70
IGHJ5
30
25
20
15
10
10
30
50
70
IGHJ4
60
55
50
45
40
10
30
50
70
IGHJ3
20
15
10
5
10
30
50
70
IGHJ2
8
6
4
2
10
30
50
70
IGHJ1
7
5
3
1
10
30
50
70

### Slide 12
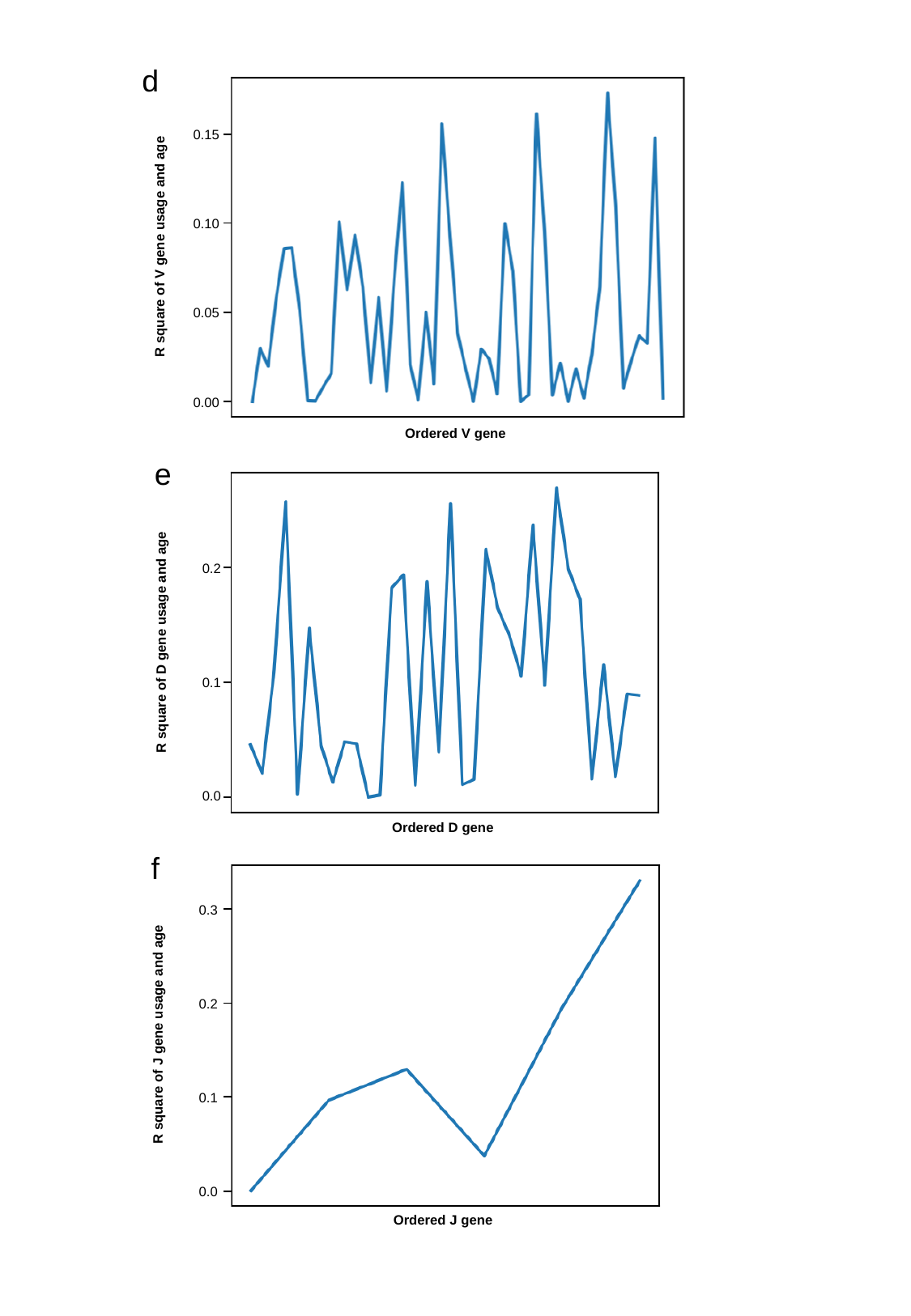

d
0.15
0.10
0.05
0.00
R square of V gene usage and age
Ordered V gene
e
0.2
0.1
0.0
R square of D gene usage and age
Ordered D gene
f
0.3
0.2
0.1
0.0
R square of J gene usage and age
Ordered J gene

### Slide 13
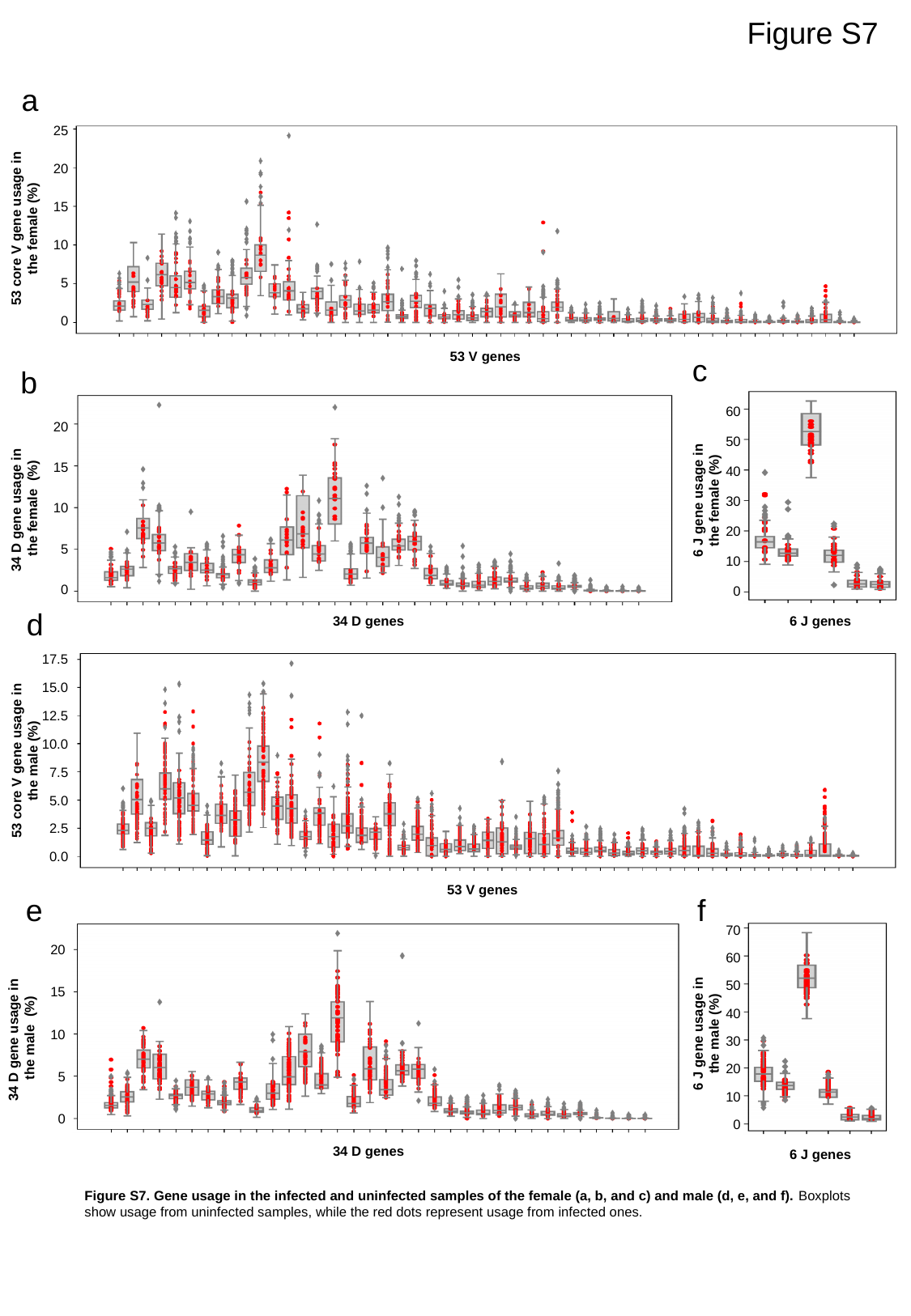

Figure S7
a
25
20
15
53 core V gene usage in the female (%)
10
5
0
53 V genes
c
b
60
20
50
15
40
6 J gene usage in the female (%)
34 D gene usage in the female (%)
30
10
20
5
10
0
0
d
34 D genes
6 J genes
17.5
15.0
12.5
10.0
53 core V gene usage in the male (%)
7.5
5.0
2.5
0.0
53 V genes
e
f
70
20
60
50
15
40
6 J gene usage in the male (%)
34 D gene usage in the male (%)
10
30
20
5
10
0
0
34 D genes
6 J genes
Figure S7. Gene usage in the infected and uninfected samples of the female (a, b, and c) and male (d, e, and f). Boxplots show usage from uninfected samples, while the red dots represent usage from infected ones.

### Slide 14
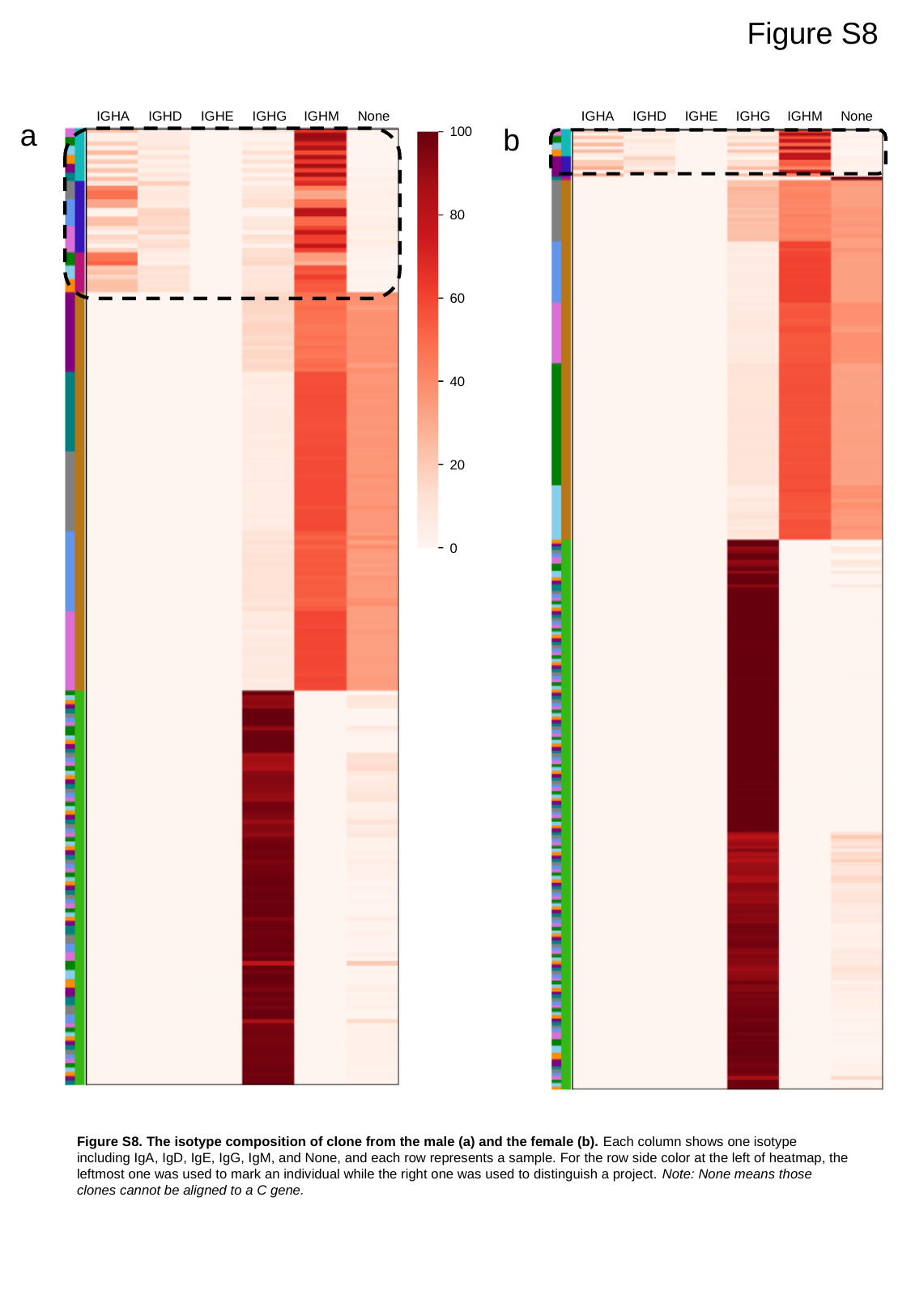

Figure S8
IGHA
IGHD
IGHE
IGHG
IGHM
None
IGHA
IGHD
IGHE
IGHG
IGHM
None
a
b
100
80
60
40
20
0
Figure S8. The isotype composition of clone from the male (a) and the female (b). Each column shows one isotype including IgA, IgD, IgE, IgG, IgM, and None, and each row represents a sample. For the row side color at the left of heatmap, the leftmost one was used to mark an individual while the right one was used to distinguish a project. Note: None means those clones cannot be aligned to a C gene.

### Slide 15
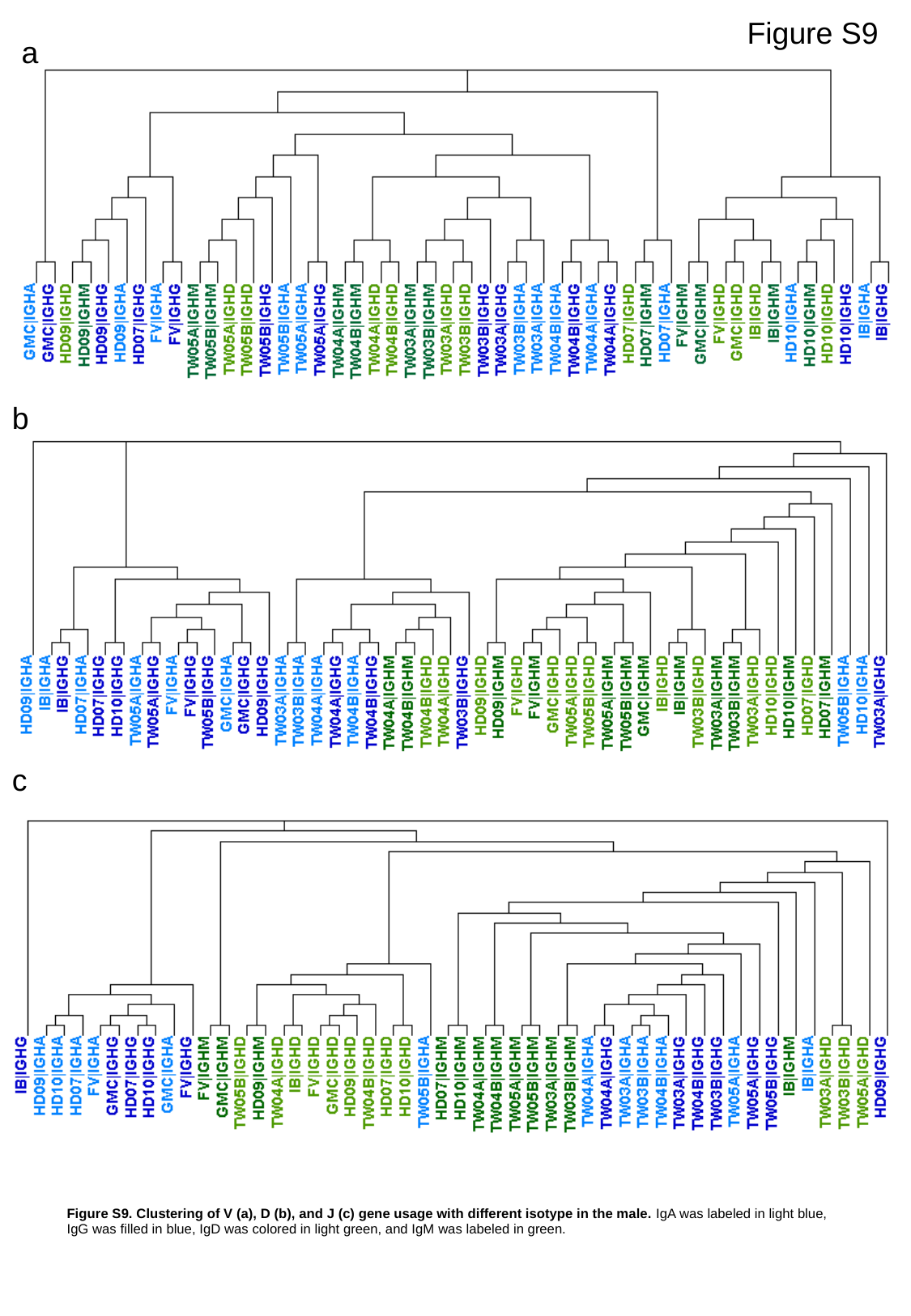

Figure S9
a
b
c
Figure S9. Clustering of V (a), D (b), and J (c) gene usage with different isotype in the male. IgA was labeled in light blue, IgG was filled in blue, IgD was colored in light green, and IgM was labeled in green.

### Slide 16
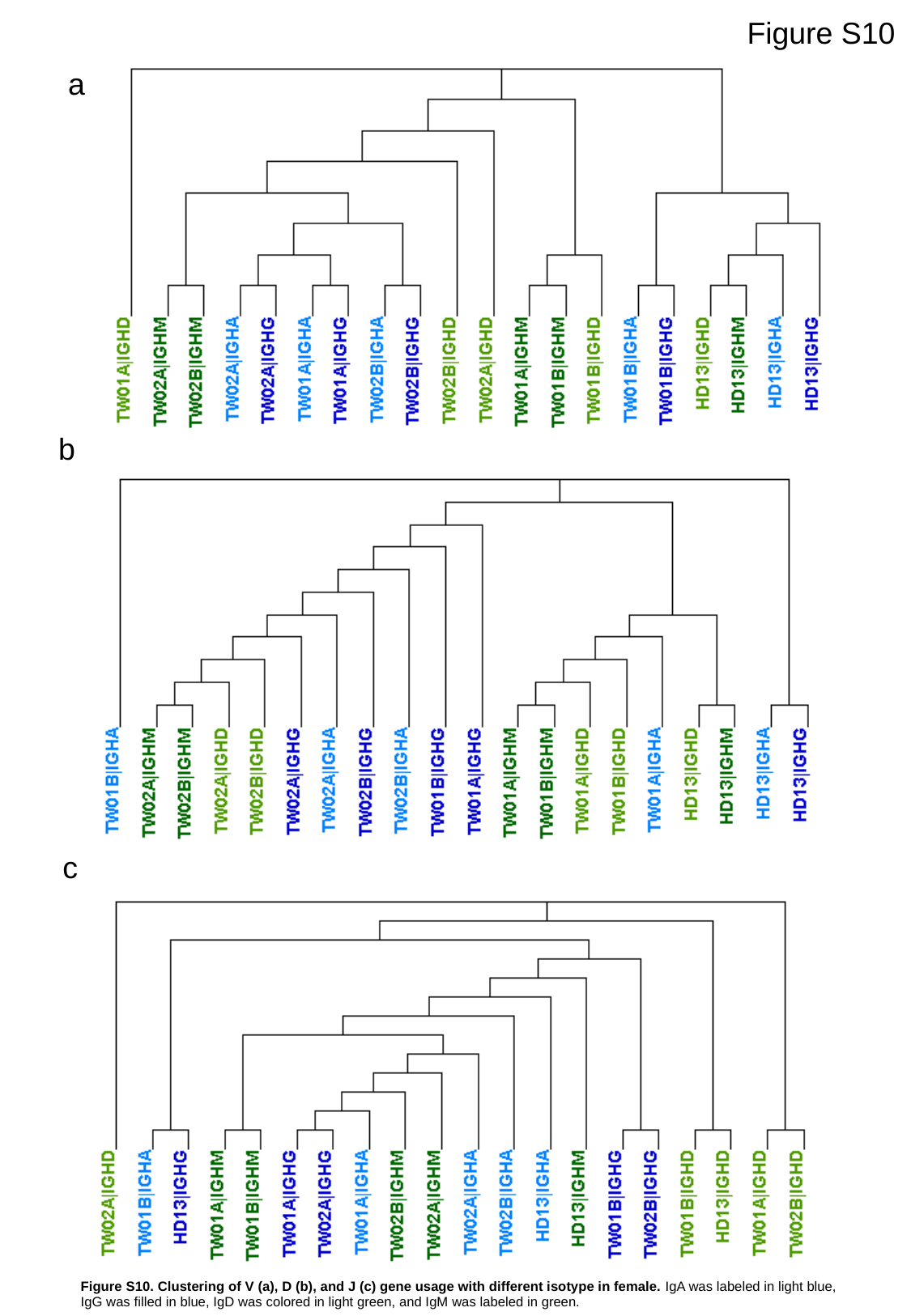

Figure S10
a
b
c
Figure S10. Clustering of V (a), D (b), and J (c) gene usage with different isotype in female. IgA was labeled in light blue, IgG was filled in blue, IgD was colored in light green, and IgM was labeled in green.

### Slide 17
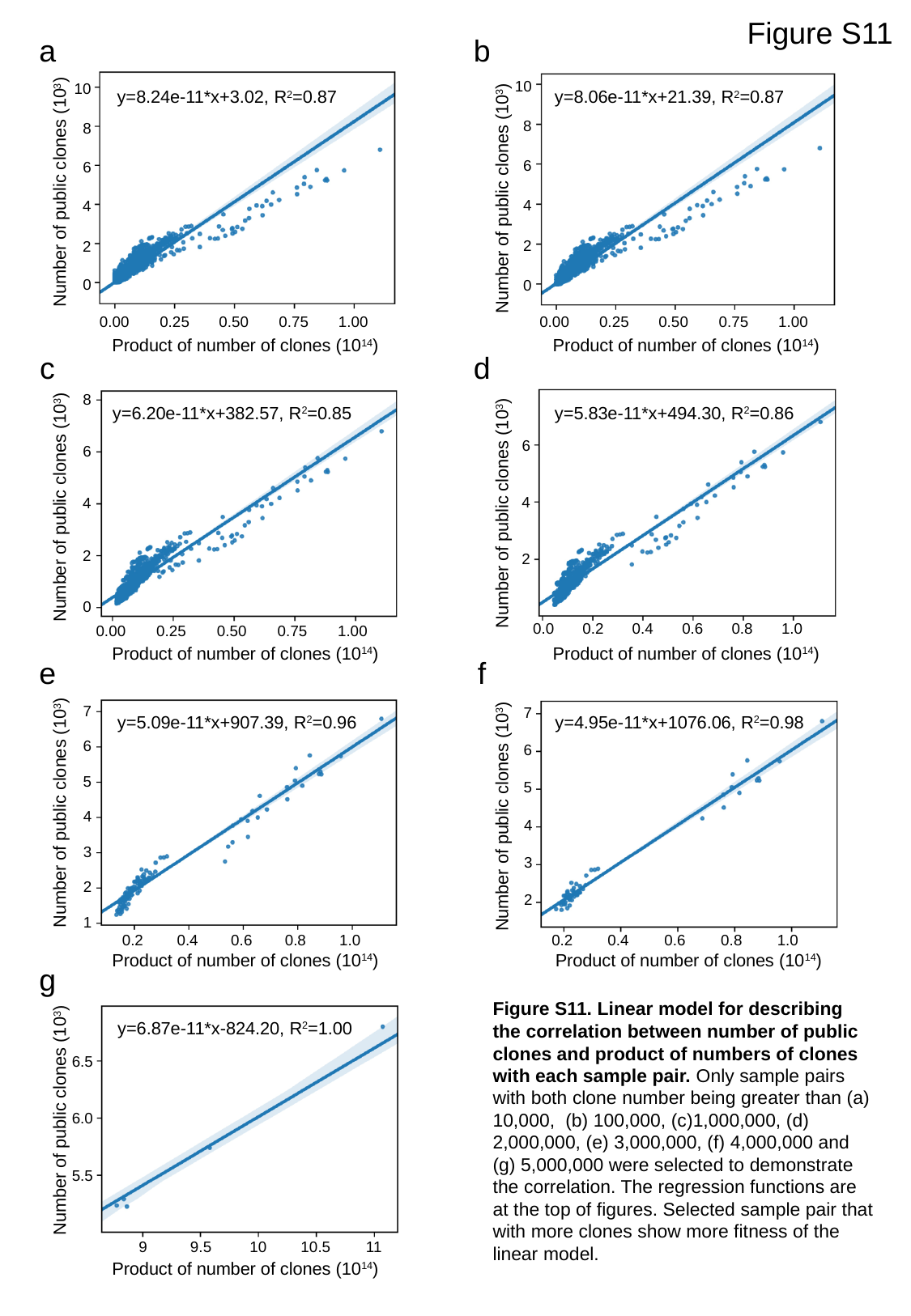

Figure S11
a
b
10
8
6
4
2
0
10
8
6
4
2
0
y=8.24e-11*x+3.02, R^2^=0.87
y=8.06e-11*x+21.39, R^2^=0.87
Number of public clones (10^3^)
Number of public clones (10^3^)
0.00
0.25
0.50
0.75
1.00
0.00
0.25
0.50
0.75
1.00
Product of number of clones (10^14^)
Product of number of clones (10^14^)
c
d
8
6
4
2
0
y=6.20e-11*x+382.57, R^2^=0.85
y=5.83e-11*x+494.30, R^2^=0.86
6
4
2
Number of public clones (10^3^)
Number of public clones (10^3^)
0.2
0.0
0.4
0.6
0.8
1.0
0.00
0.25
0.50
0.75
1.00
Product of number of clones (10^14^)
Product of number of clones (10^14^)
e
f
7
6
5
4
3
2
1
7
6
5
4
3
2
y=5.09e-11*x+907.39, R^2^=0.96
y=4.95e-11*x+1076.06, R^2^=0.98
Number of public clones (10^3^)
Number of public clones (10^3^)
0.2
0.4
0.6
0.8
1.0
0.2
0.4
0.6
0.8
1.0
Product of number of clones (10^14^)
Product of number of clones (10^14^)
g
Figure S11. Linear model for describing the correlation between number of public clones and product of numbers of clones with each sample pair. Only sample pairs with both clone number being greater than (a) 10,000, (b) 100,000, (c)1,000,000, (d) 2,000,000, (e) 3,000,000, (f) 4,000,000 and (g) 5,000,000 were selected to demonstrate the correlation. The regression functions are at the top of figures. Selected sample pair that with more clones show more fitness of the linear model.
y=6.87e-11*x-824.20, R^2^=1.00
6.5
6.0
5.5
Number of public clones (10^3^)
9
9.5
10
10.5
11
Product of number of clones (10^14^)

### Slide 18
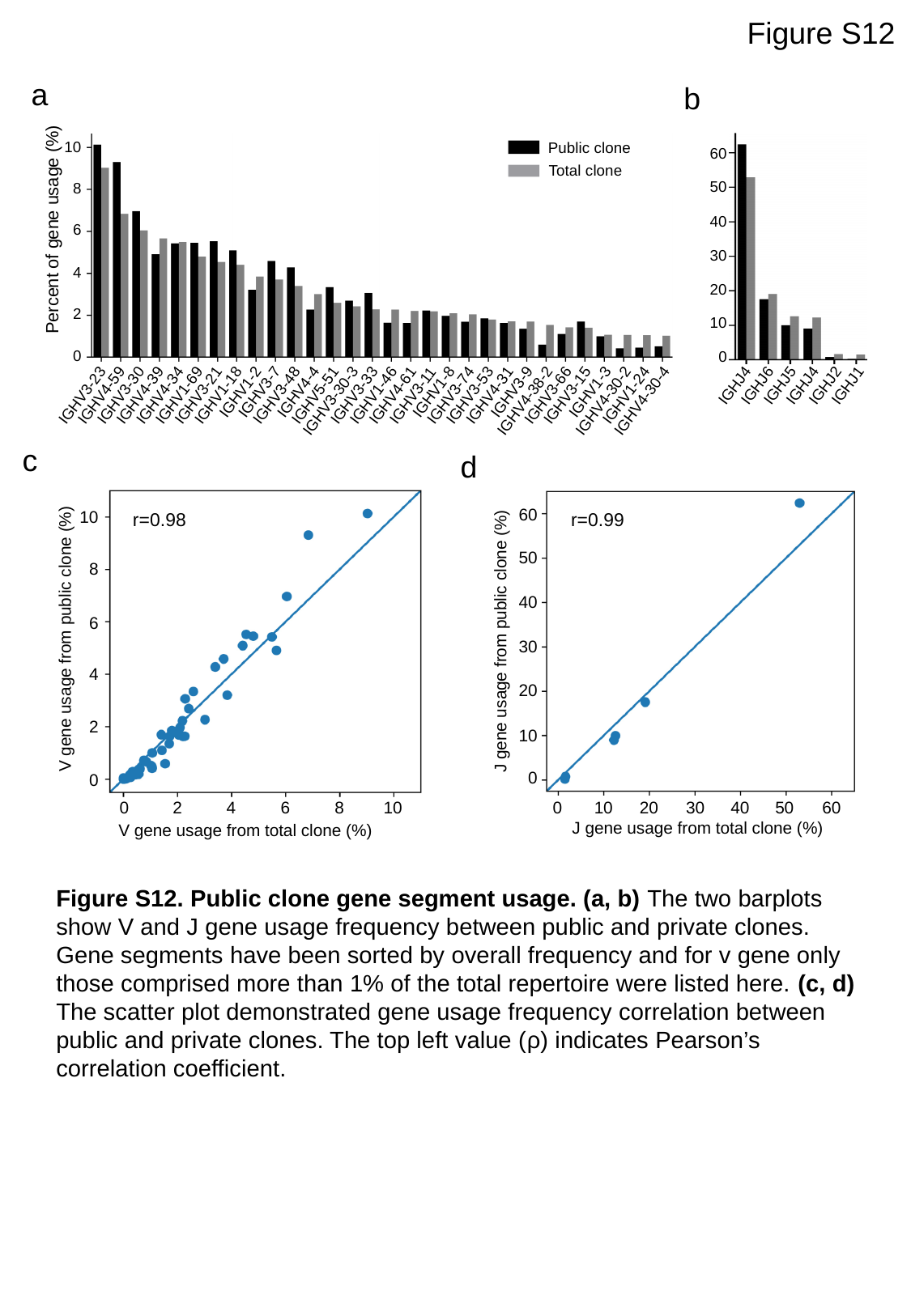

Figure S12
a
b
10
8
6
4
2
0
Public clone
60
50
40
30
20
10
0
Total clone
Percent of gene usage (%)
IGHV1-2
IGHV3-7
IGHV4-4
IGHV1-8
IGHV3-9
IGHV1-3
IGHV3-23
IGHV4-59
IGHV3-30
IGHV4-39
IGHV4-34
IGHV1-69
IGHV3-21
IGHV1-18
IGHV3-48
IGHV5-51
IGHV3-33
IGHV1-46
IGHV4-61
IGHV3-11
IGHV3-74
IGHV3-53
IGHV4-31
IGHV3-66
IGHV3-15
IGHV1-24
IGHV3-30-3
IGHV4-38-2
IGHV4-30-2
IGHV4-30-4
IGHJ4
IGHJ6
IGHJ5
IGHJ4
IGHJ2
IGHJ1
c
d
60
50
40
30
20
10
0
10
8
6
4
2
0
r=0.99
r=0.98
 V gene usage from public clone (%)
J gene usage from public clone (%)
0
2
4
6
8
10
0
10
20
30
40
50
60
J gene usage from total clone (%)
V gene usage from total clone (%)
Figure S12. Public clone gene segment usage. (a, b) The two barplots show V and J gene usage frequency between public and private clones. Gene segments have been sorted by overall frequency and for v gene only those comprised more than 1% of the total repertoire were listed here. (c, d) The scatter plot demonstrated gene usage frequency correlation between public and private clones. The top left value (ρ) indicates Pearson’s correlation coefficient.

### Slide 19
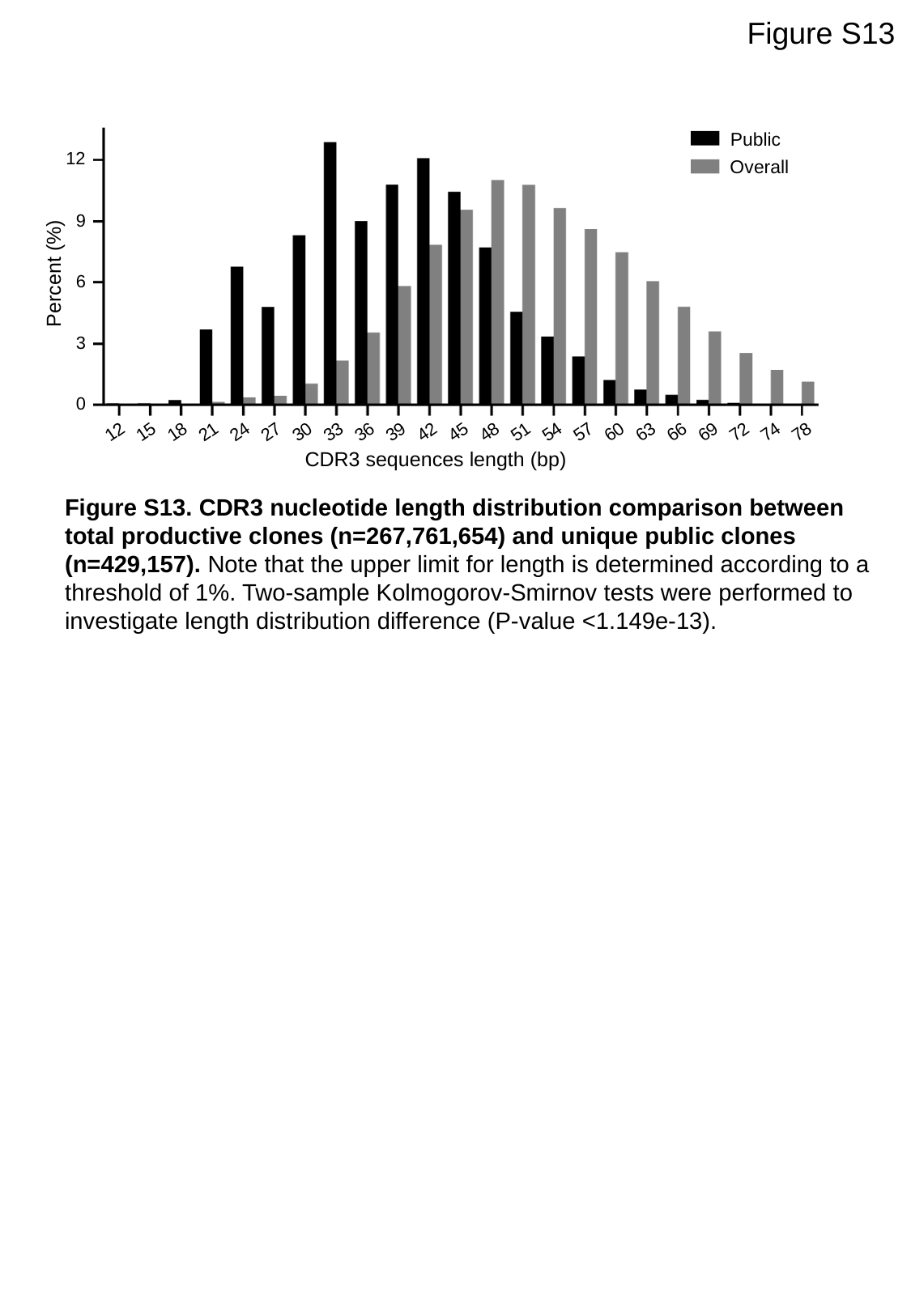

Figure S13
Public
12
9
6
3
0
Overall
Percent (%)
12
15
18
21
24
27
30
33
36
39
42
45
48
51
54
57
60
63
66
69
72
74
78
CDR3 sequences length (bp)
Figure S13. CDR3 nucleotide length distribution comparison between total productive clones (n=267,761,654) and unique public clones (n=429,157). Note that the upper limit for length is determined according to a threshold of 1%. Two-sample Kolmogorov-Smirnov tests were performed to investigate length distribution difference (P-value <1.149e-13).

### Slide 20
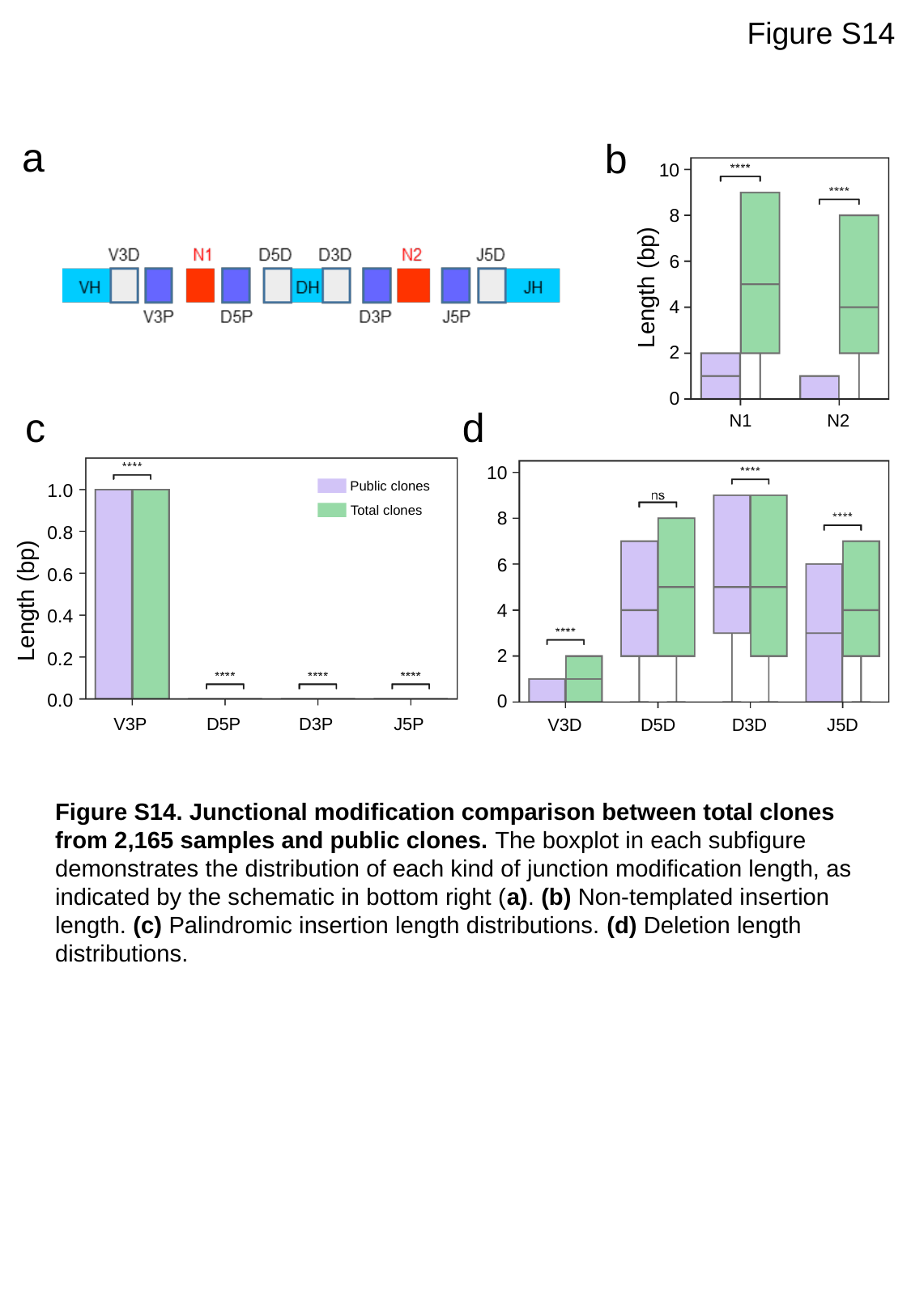

Figure S14
a
b
10
8
6
4
2
0
Length (bp)
c
d
N1
N2
10
8
6
4
2
0
Public clones
Total clones
1.0
0.8
0.6
0.4
0.2
0.0
Length (bp)
V3P
D5P
D3P
J5P
V3D
D5D
D3D
J5D
Figure S14. Junctional modification comparison between total clones from 2,165 samples and public clones. The boxplot in each subfigure demonstrates the distribution of each kind of junction modification length, as indicated by the schematic in bottom right (a). (b) Non-templated insertion length. (c) Palindromic insertion length distributions. (d) Deletion length distributions.

### Slide 21
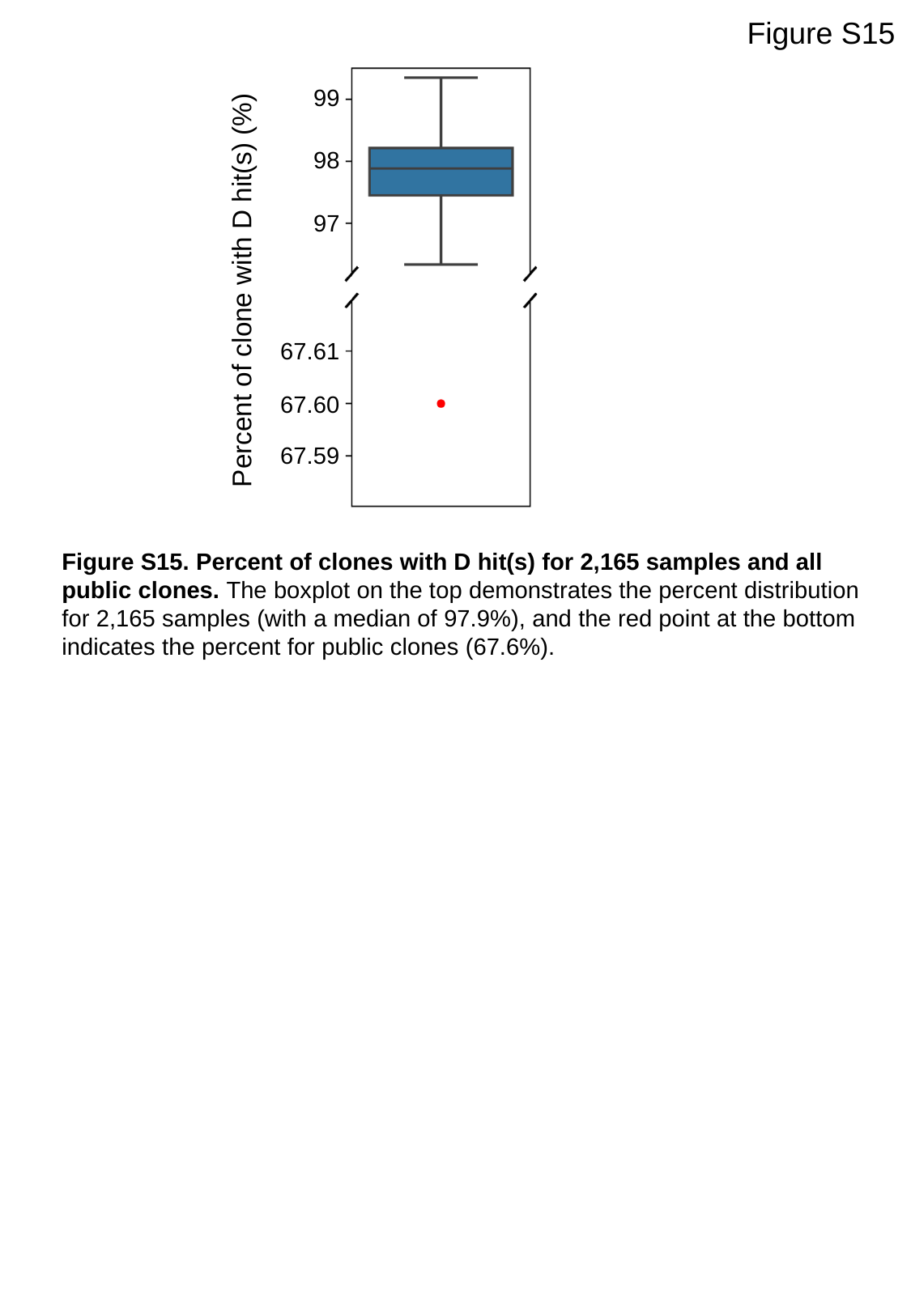

Figure S15
99
98
97
67.61
67.60
67.59
Percent of clone with D hit(s) (%)
Figure S15. Percent of clones with D hit(s) for 2,165 samples and all public clones. The boxplot on the top demonstrates the percent distribution for 2,165 samples (with a median of 97.9%), and the red point at the bottom indicates the percent for public clones (67.6%).

### Slide 22
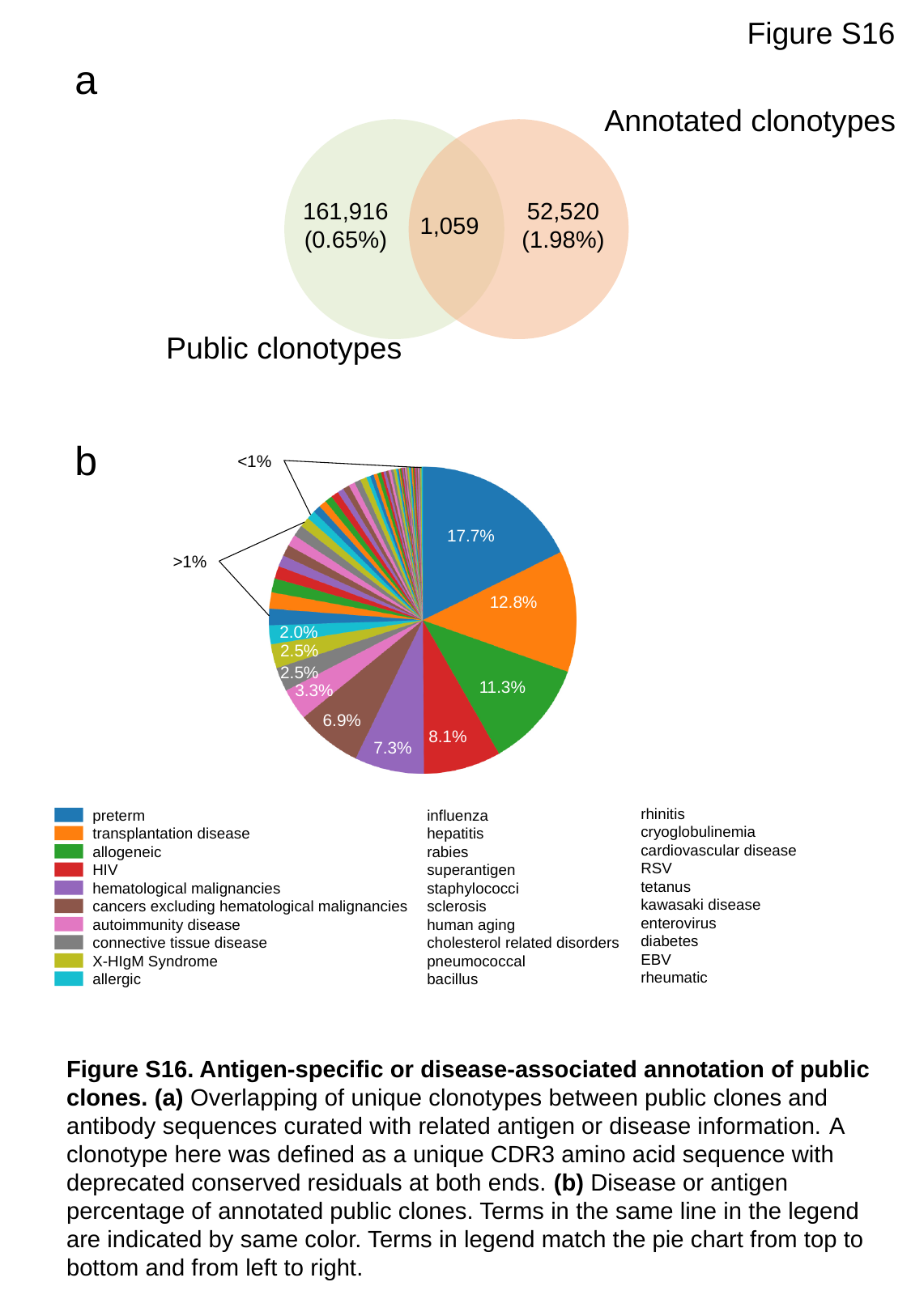

Figure S16
a
Annotated clonotypes
161,916
(0.65%)
52,520
(1.98%)
1,059
Public clonotypes
b
<1%
17.7%
>1%
12.8%
2.0%
2.5%
2.5%
11.3%
3.3%
6.9%
8.1%
7.3%
rhinitis
cryoglobulinemia
cardiovascular disease
RSV
tetanus
kawasaki disease
enterovirus
diabetes
EBV
rheumatic
influenza
hepatitis
rabies
superantigen
staphylococci
sclerosis
human aging
cholesterol related disorders
pneumococcal
bacillus
preterm
transplantation disease
allogeneic
HIV
hematological malignancies
cancers excluding hematological malignancies
autoimmunity disease
connective tissue disease
X-HIgM Syndrome
allergic
Figure S16. Antigen-specific or disease-associated annotation of public clones. (a) Overlapping of unique clonotypes between public clones and antibody sequences curated with related antigen or disease information. A clonotype here was defined as a unique CDR3 amino acid sequence with deprecated conserved residuals at both ends. (b) Disease or antigen percentage of annotated public clones. Terms in the same line in the legend are indicated by same color. Terms in legend match the pie chart from top to bottom and from left to right.

### Slide 23
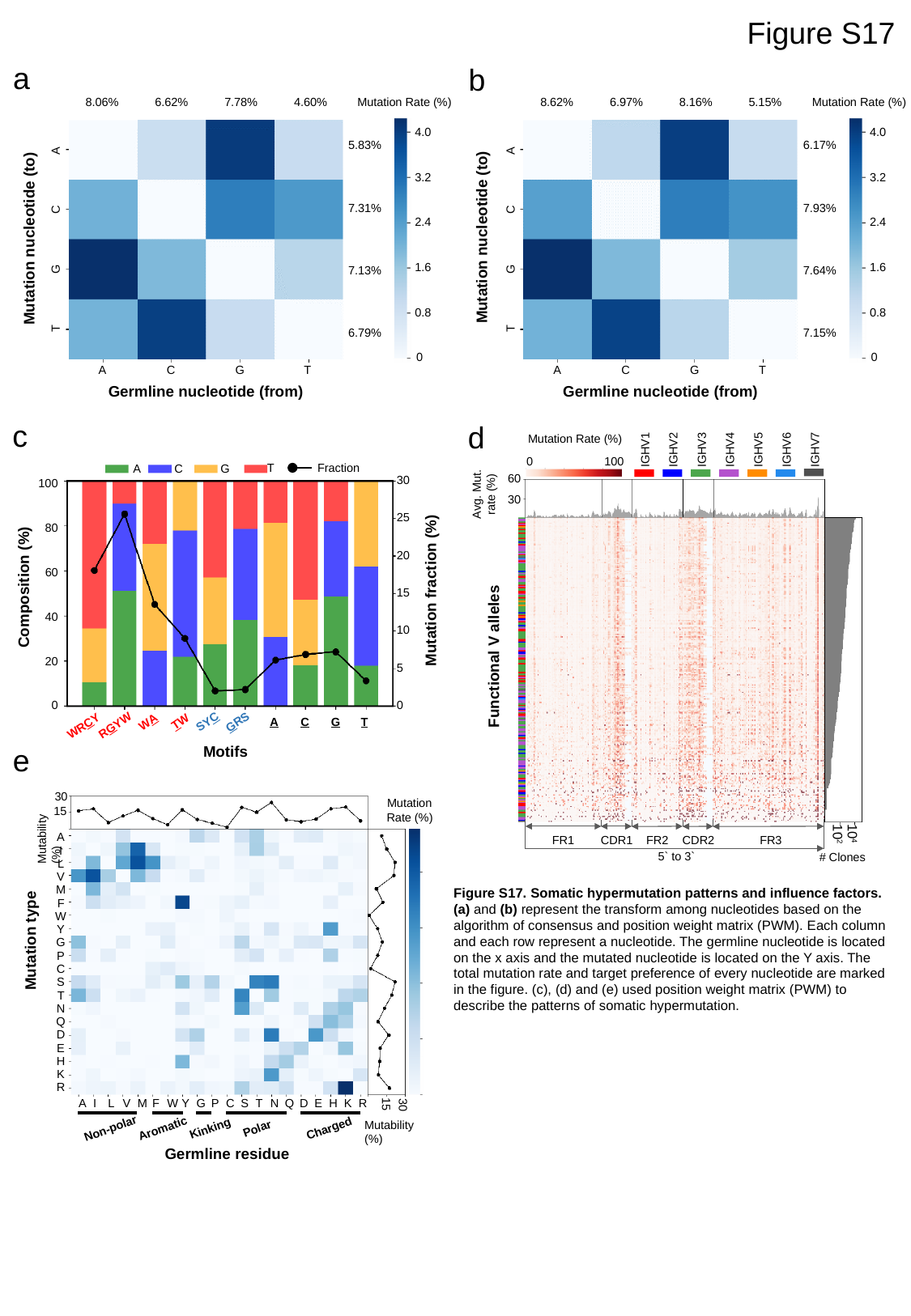

Figure S17
a
b
8.06%
6.62%
7.78%
4.60%
4.0
3.2
2.4
1.6
0.8
0
5.83%
A
7.31%
C
Mutation nucleotide (to)
G
7.13%
T
6.79%
A
C
G
T
Germline nucleotide (from)
Mutation Rate (%)
Mutation Rate (%)
8.62%
6.97%
8.16%
5.15%
4.0
3.2
2.4
1.6
0.8
0
6.17%
A
7.93%
C
Mutation nucleotide (to)
G
7.64%
T
7.15%
A
C
G
T
Germline nucleotide (from)
Mutation Rate (%)
IGHV1
IGHV2
IGHV3
IGHV4
IGHV5
IGHV6
IGHV7
100
0
60
Avg. Mut.
rate (%)
30
Functional V alleles
10^4^
10^2^
FR1
CDR1
FR2
CDR2
FR3
5` to 3`
#### Clones
c
d
Fraction
T
A
C
G
30
100
25
80
20
60
15
40
10
20
5
0
0
WA
TW
SYC
GRS
A
C
G
T
RGYW
WRCY
Composition (%)
Mutation fraction (%)
Motifs
e
30
Mutation Rate (%)
15
Mutability (%)
A
I
L
V
M
F
W
Y
G
P
C
S
T
N
Q
D
E
H
K
R
Mutation type
A
I
L
V
M
F
W
Y
G
P
C
S
T
N
Q
D
E
H
K
R
15
30
Mutability (%)
Non-polar
Aromatic
Kinking
Polar
Charged
Germline residue
Figure S17. Somatic hypermutation patterns and influence factors. (a) and (b) represent the transform among nucleotides based on the algorithm of consensus and position weight matrix (PWM). Each column and each row represent a nucleotide. The germline nucleotide is located on the x axis and the mutated nucleotide is located on the Y axis. The total mutation rate and target preference of every nucleotide are marked in the figure. (c), (d) and (e) used position weight matrix (PWM) to describe the patterns of somatic hypermutation.

### Slide 24
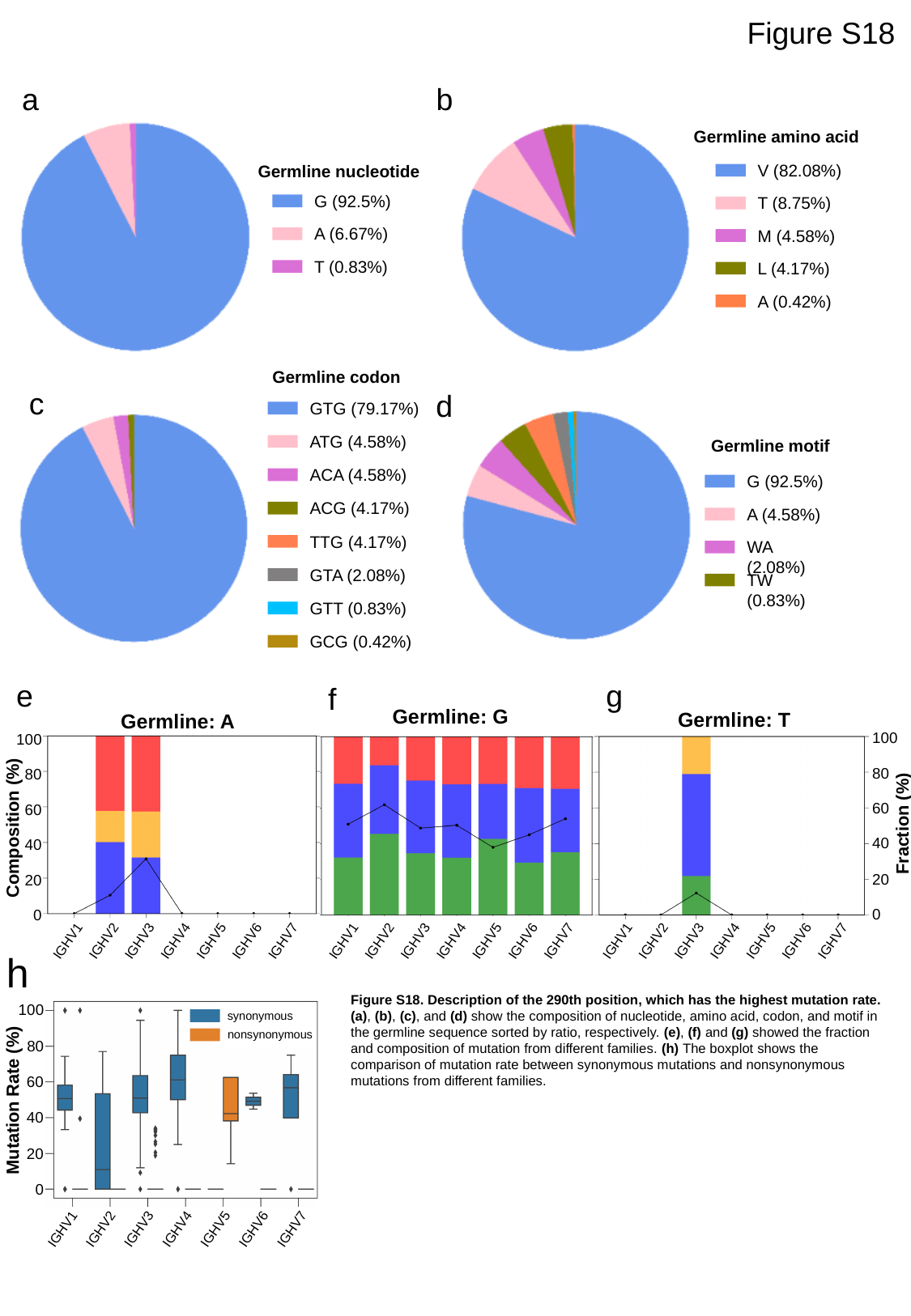

Figure S18
a
b
Germline amino acid
V (82.08%)
T (8.75%)
M (4.58%)
L (4.17%)
A (0.42%)
Germline nucleotide
G (92.5%)
A (6.67%)
T (0.83%)
Germline codon
GTG (79.17%)
ATG (4.58%)
ACA (4.58%)
ACG (4.17%)
TTG (4.17%)
GTA (2.08%)
GTT (0.83%)
GCG (0.42%)
c
d
Germline motif
G (92.5%)
A (4.58%)
WA (2.08%)
TW (0.83%)
g
e
f
Germline: G
Germline: T
Germline: A
100
80
60
40
20
0
100
80
60
40
20
0
A
C
G
T
Fraction (%)
Composition (%)
IGHV1
IGHV2
IGHV3
IGHV4
IGHV5
IGHV6
IGHV7
IGHV1
IGHV2
IGHV3
IGHV4
IGHV5
IGHV6
IGHV7
IGHV1
IGHV2
IGHV3
IGHV4
IGHV5
IGHV6
IGHV7
h
Figure S18. Description of the 290th position, which has the highest mutation rate. (a), (b), (c), and (d) show the composition of nucleotide, amino acid, codon, and motif in the germline sequence sorted by ratio, respectively. (e), (f) and (g) showed the fraction and composition of mutation from different families. (h) The boxplot shows the comparison of mutation rate between synonymous mutations and nonsynonymous mutations from different families.
100
80
60
40
20
0
synonymous
nonsynonymous
Mutation Rate (%)
IGHV1
IGHV2
IGHV3
IGHV4
IGHV5
IGHV6
IGHV7

### Slide 25
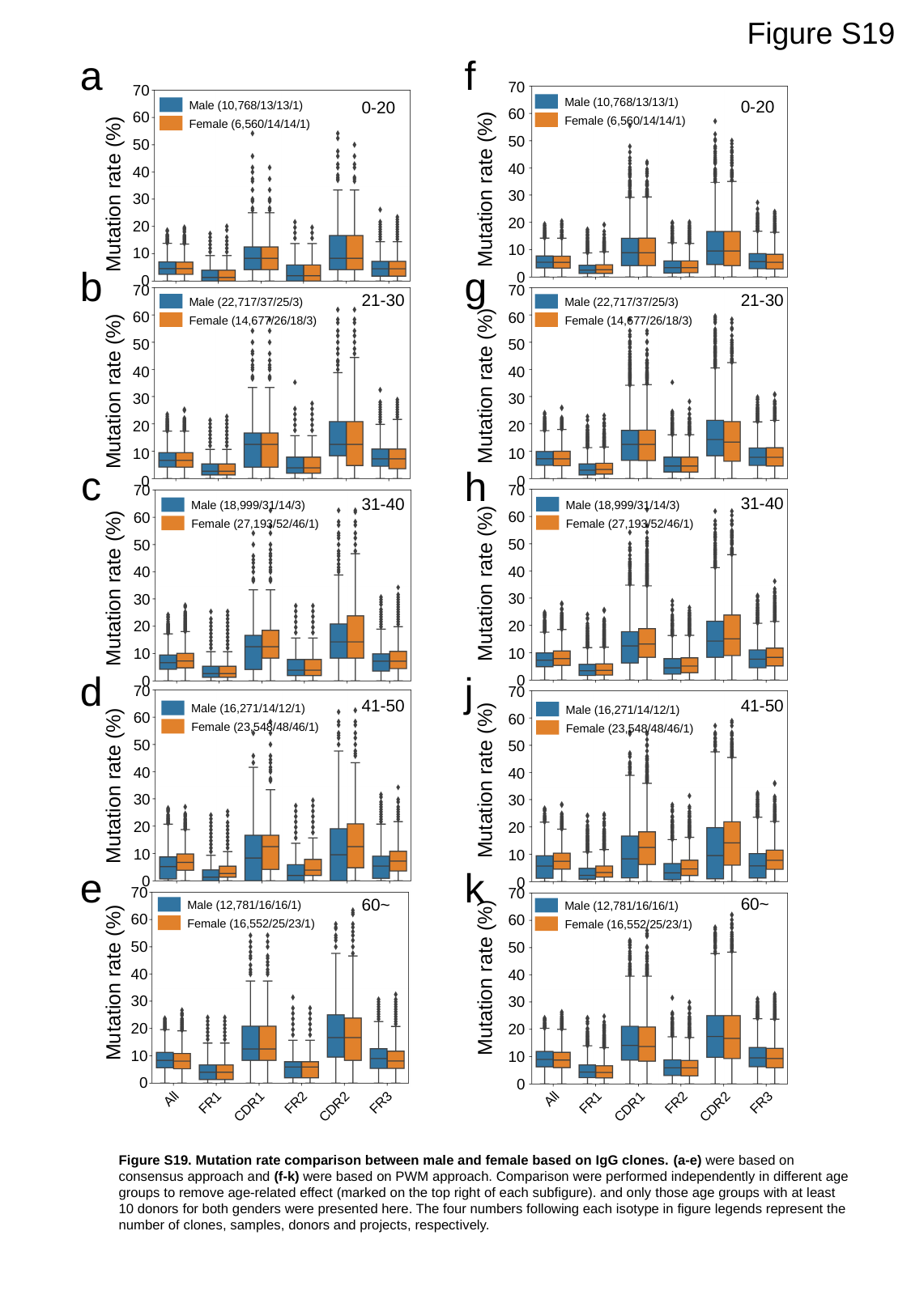

Figure S19
a
f
70
60
50
40
30
20
10
0
70
60
50
40
30
20
10
0
Male (10,768/13/13/1)
Female (6,560/14/14/1)
0-20
0-20
Male (10,768/13/13/1)
Female (6,560/14/14/1)
Mutation rate (%)
Mutation rate (%)
b
g
70
60
50
40
30
20
10
0
70
60
50
40
30
20
10
0
21-30
21-30
Male (22,717/37/25/3)
Female (14,677/26/18/3)
Male (22,717/37/25/3)
Female (14,677/26/18/3)
Mutation rate (%)
Mutation rate (%)
c
h
70
60
50
40
30
20
10
0
70
60
50
40
30
20
10
0
31-40
31-40
Male (18,999/31/14/3)
Female (27,193/52/46/1)
Male (18,999/31/14/3)
Female (27,193/52/46/1)
Mutation rate (%)
Mutation rate (%)
d
j
70
60
50
40
30
20
10
0
70
60
50
40
30
20
10
0
41-50
41-50
Male (16,271/14/12/1)
Female (23,548/48/46/1)
Male (16,271/14/12/1)
Female (23,548/48/46/1)
Mutation rate (%)
Mutation rate (%)
e
k
70
60
50
40
30
20
10
0
70
60
50
40
30
20
10
0
60~
60~
Male (12,781/16/16/1)
Female (16,552/25/23/1)
Male (12,781/16/16/1)
Female (16,552/25/23/1)
Mutation rate (%)
Mutation rate (%)
All
FR1
FR2
FR3
CDR1
CDR2
All
FR1
FR2
FR3
CDR1
CDR2
Figure S19. Mutation rate comparison between male and female based on IgG clones. (a-e) were based on consensus approach and (f-k) were based on PWM approach. Comparison were performed independently in different age groups to remove age-related effect (marked on the top right of each subfigure). and only those age groups with at least 10 donors for both genders were presented here. The four numbers following each isotype in figure legends represent the number of clones, samples, donors and projects, respectively.

### Slide 26
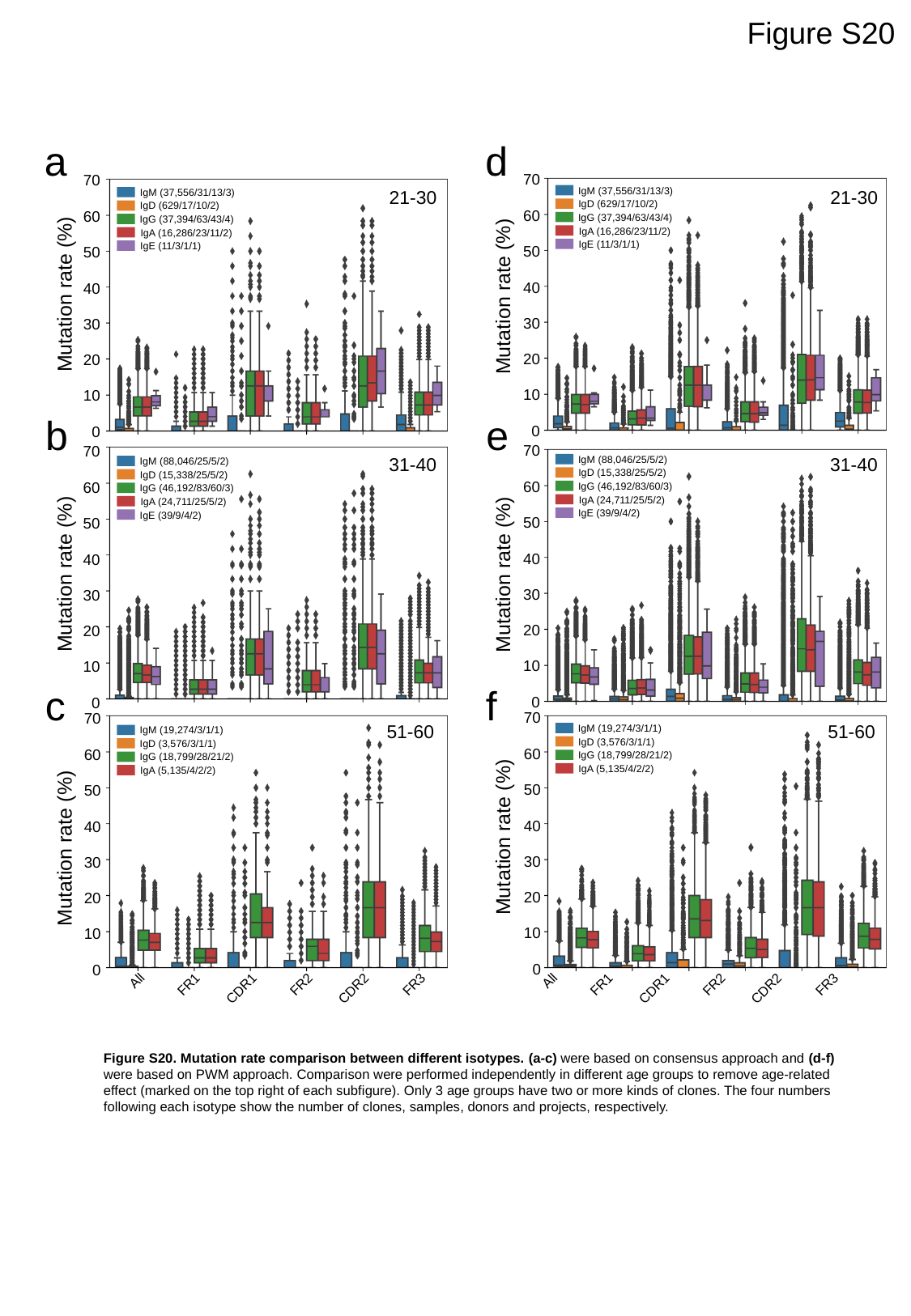

Figure S20
a
d
70
60
50
40
30
20
10
0
70
60
50
40
30
20
10
0
IgM (37,556/31/13/3)
IgD (629/17/10/2)
IgG (37,394/63/43/4)
IgA (16,286/23/11/2)
IgE (11/3/1/1)
21-30
21-30
IgM (37,556/31/13/3)
IgD (629/17/10/2)
IgG (37,394/63/43/4)
IgA (16,286/23/11/2)
IgE (11/3/1/1)
Mutation rate (%)
Mutation rate (%)
b
e
70
60
50
40
30
20
10
0
70
60
50
40
30
20
10
0
IgM (88,046/25/5/2)
IgD (15,338/25/5/2)
IgG (46,192/83/60/3)
IgA (24,711/25/5/2)
IgE (39/9/4/2)
31-40
31-40
IgM (88,046/25/5/2)
IgD (15,338/25/5/2)
IgG (46,192/83/60/3)
IgA (24,711/25/5/2)
IgE (39/9/4/2)
Mutation rate (%)
Mutation rate (%)
c
f
70
60
50
40
30
20
10
0
70
60
50
40
30
20
10
0
51-60
51-60
IgM (19,274/3/1/1)
IgD (3,576/3/1/1)
IgG (18,799/28/21/2)
IgA (5,135/4/2/2)
IgM (19,274/3/1/1)
IgD (3,576/3/1/1)
IgG (18,799/28/21/2)
IgA (5,135/4/2/2)
Mutation rate (%)
Mutation rate (%)
All
FR1
FR2
FR3
CDR1
CDR2
All
FR1
FR2
FR3
CDR1
CDR2
Figure S20. Mutation rate comparison between different isotypes. (a-c) were based on consensus approach and (d-f) were based on PWM approach. Comparison were performed independently in different age groups to remove age-related effect (marked on the top right of each subfigure). Only 3 age groups have two or more kinds of clones. The four numbers following each isotype show the number of clones, samples, donors and projects, respectively.
