## Supplemental Table 1 for "Large-scale Analysis of 2,152 dataset reveals key features of B cell biology and the antibody repertoire"

**Table S1a Overview of 53 core V gene set**

| **Gene** | **# Samples** | **# Donors** | **Min** | **Max** | **Mean** | **Median** |
| --- | --- | --- | --- | --- | --- | --- |
| IGHV5-51 | 2,125 | 582 | 0.0287 | 8.9112 | 2.6407 | 2.4573 |
| IGHV4-59 | 2,148 | 582 | 0.0308 | 49.1694 | 4.7669 | 4.3106 |
| IGHV4-4 | 2,148 | 582 | 0.0341 | 17.5948 | 2.5258 | 2.2752 |
| IGHV4-39 | 2,151 | 582 | 0.0044 | 21.3033 | 5.0249 | 4.7783 |
| IGHV4-34 | 2,145 | 582 | 0.0089 | 17.9332 | 4.2332 | 3.9451 |
| IGHV3-7 | 2,135 | 582 | 0.0076 | 77.1014 | 4.6246 | 4.1998 |
| IGHV3-66 | 2,134 | 582 | 0.0095 | 27.4476 | 1.3993 | 1.3033 |
| IGHV3-48 | 2,120 | 582 | 0.0186 | 20.2547 | 3.3840 | 3.3173 |
| IGHV3-33 | 2,147 | 582 | 0.0170 | 20.7993 | 3.1910 | 3.0822 |
| IGHV3-30 | 2,152 | 582 | 0.0295 | 18.8430 | 6.1333 | 6.0589 |
| IGHV3-23 | 2,152 | 582 | 0.0761 | 30.3371 | 8.4535 | 8.3656 |
| IGHV3-21 | 2,148 | 582 | 0.0832 | 34.7654 | 4.1481 | 3.9091 |
| IGHV1-69 | 2,148 | 582 | 0.0822 | 24.1197 | 5.0032 | 4.6882 |
| IGHV1-46 | 2,147 | 582 | 0.0133 | 13.8889 | 2.1305 | 1.9494 |
| IGHV1-18 | 2,149 | 582 | 0.0069 | 17.8841 | 4.0165 | 3.8104 |
| IGHV4-61 | 2,145 | 581 | 0.0218 | 16.8344 | 2.0663 | 1.9019 |
| IGHV3-74 | 2,147 | 581 | 0.0511 | 25.6112 | 3.0947 | 2.7316 |
| IGHV3-15 | 2,142 | 581 | 0.0227 | 12.4923 | 2.0282 | 1.8406 |
| IGHV3-11 | 2,147 | 581 | 0.0170 | 10.8959 | 2.4050 | 2.3426 |
| IGHV1-3 | 2,105 | 581 | 0.0028 | 12.0907 | 1.4448 | 1.3441 |
| IGHV1-2 | 2,144 | 581 | 0.0374 | 23.4793 | 3.5171 | 3.4370 |
| IGHV6-1 | 2,117 | 580 | 0.0009 | 7.9248 | 0.9117 | 0.7671 |
| IGHV3-53 | 2,139 | 580 | 0.0076 | 13.9795 | 2.1108 | 1.9331 |
| IGHV3-69-1 | 2,128 | 579 | 0.0076 | 8.4994 | 0.7376 | 0.5854 |
| IGHV2-5 | 2,112 | 579 | 0.0025 | 8.4298 | 1.4993 | 1.3095 |
| IGHV3-49 | 1,986 | 577 | 0.0067 | 7.0414 | 0.7862 | 0.7377 |
| IGHV1-8 | 2,122 | 577 | 0.0022 | 10.3514 | 1.5982 | 1.4905 |
| IGHV1-24 | 2,049 | 576 | 0.0060 | 10.2255 | 0.7599 | 0.6971 |
| IGHV3-30-3 | 2,109 | 575 | 0.0026 | 10.6330 | 2.1013 | 2.0393 |
| IGHV4-31 | 2,118 | 573 | 0.0007 | 9.4527 | 1.9032 | 1.6465 |
| IGHV4-38-2 | 2,051 | 572 | 0.0014 | 12.8967 | 1.1189 | 0.5846 |
| IGHV3-9 | 2,041 | 571 | 0.0011 | 11.7705 | 2.0359 | 1.9294 |
| IGHV3-73 | 2,077 | 571 | 0.0028 | 7.9491 | 0.5707 | 0.4796 |
| IGHV3-72 | 2,079 | 571 | 0.0150 | 3.6374 | 0.5426 | 0.3868 |
| IGHV3-64 | 2,059 | 569 | 0.0076 | 5.3632 | 0.4662 | 0.3776 |
| IGHV4-30-2 | 1,973 | 563 | 0.0009 | 11.6279 | 0.6241 | 0.4471 |
| IGHV3-43 | 1,999 | 558 | 0.0020 | 2.7835 | 0.3820 | 0.2719 |
| IGHV2-70 | 1,866 | 558 | 0.0002 | 7.8502 | 0.4759 | 0.3329 |
| IGHV3-13 | 2,043 | 550 | 0.0049 | 16.6667 | 0.5101 | 0.3927 |
| IGHV3-20 | 2,015 | 547 | 0.0016 | 4.2923 | 0.5653 | 0.3562 |
| IGHV3-30-5 | 1,723 | 537 | 0.0006 | 6.3258 | 0.3868 | 0.1600 |
| IGHV4-30-4 | 2,022 | 536 | 0.0007 | 12.8851 | 1.0364 | 0.7698 |
| IGHV7-4-1 | 1,852 | 535 | 0.0001 | 9.0114 | 0.6719 | 0.2304 |
| IGHV3-43D | 1,809 | 533 | 0.0002 | 2.0087 | 0.1968 | 0.1360 |
| IGHV5-10-1 | 1,655 | 532 | 0.0000 | 4.9083 | 0.6897 | 0.4599 |
| IGHV4/OR15-8 | 1,922 | 523 | 0.0006 | 8.5553 | 0.1991 | 0.0885 |
| IGHV4-28 | 1,819 | 508 | 0.0009 | 3.8003 | 0.1270 | 0.0630 |
| IGHV1-58 | 1,733 | 505 | 0.0049 | 2.5806 | 0.2757 | 0.2312 |
| IGHV2-26 | 1,823 | 495 | 0.0003 | 20.0000 | 0.2904 | 0.1619 |
| IGHV3-64D | 1,518 | 488 | 0.0002 | 2.6716 | 0.3317 | 0.2439 |
| IGHV3-23D | 881 | 486 | 0.0000 | 11.5998 | 1.0508 | 0.6474 |
| IGHV3-38 | 1,766 | 468 | 0.0003 | 2.2932 | 0.0788 | 0.0443 |
| IGHV3-71 | 1,691 | 464 | 0.0016 | 2.2382 | 0.0738 | 0.0389 |

**Table S1b Overview of 49 non-core V gene**

| **Gene** | **# Samples** | **# Donors** | **Min** | **Max** | **Mean** | **Median** |
| --- | --- | --- | --- | --- | --- | --- |
| IGHV4-55 | 1,621 | 461 | 0.0005 | 2.2026 | 0.0750 | 0.0275 |
| IGHV1/OR15-1 | 1,492 | 436 | 0.0002 | 1.7559 | 0.0641 | 0.0227 |
| IGHV1-69D | 782 | 424 | 0.0000 | 5.7050 | 0.2915 | 0.0950 |
| IGHV3-NL1 | 1,646 | 422 | 0.0004 | 1.1877 | 0.0620 | 0.0320 |
| IGHV1-69-2 | 1,475 | 415 | 0.0000 | 3.4494 | 0.1848 | 0.0576 |
| IGHV1/OR15-3 | 1,297 | 399 | 0.0000 | 0.8359 | 0.0367 | 0.0130 |
| IGHV1/OR15-2 | 1,366 | 394 | 0.0001 | 0.6711 | 0.0341 | 0.0151 |
| IGHV3-38-3 | 1,678 | 387 | 0.0009 | 1.8349 | 0.1094 | 0.0673 |
| IGHV1-45 | 1,439 | 378 | 0.0007 | 0.9840 | 0.0463 | 0.0262 |
| IGHV3/OR16-8 | 1,328 | 374 | 0.0004 | 0.6369 | 0.0365 | 0.0133 |
| IGHV3/OR16-10 | 1,332 | 344 | 0.0005 | 1.0135 | 0.0402 | 0.0136 |
| IGHV3/OR16-9 | 1,302 | 342 | 0.0002 | 1.3777 | 0.0386 | 0.0151 |
| IGHV1/OR15-4 | 1,051 | 320 | 0.0000 | 0.3398 | 0.0206 | 0.0068 |
| IGHV3-52 | 1,312 | 319 | 0.0003 | 2.3050 | 0.0402 | 0.0109 |
| IGHV7-81 | 906 | 301 | 0.0000 | 2.7026 | 0.0495 | 0.0072 |
| IGHV3-35 | 1,190 | 301 | 0.0003 | 0.6849 | 0.0297 | 0.0083 |
| IGHV1/OR15-9 | 961 | 301 | 0.0000 | 0.6652 | 0.0253 | 0.0057 |
| IGHV7-34-1 | 639 | 298 | 0.0000 | 3.1310 | 0.0846 | 0.0224 |
| IGHV3-22 | 1,167 | 298 | 0.0008 | 0.4310 | 0.0242 | 0.0093 |
| IGHV3-47 | 976 | 290 | 0.0001 | 0.4348 | 0.0275 | 0.0079 |
| IGHV1/OR15-5 | 795 | 270 | 0.0000 | 0.5089 | 0.0206 | 0.0044 |
| IGHV3/OR16-13 | 1,236 | 264 | 0.0002 | 2.3649 | 0.0549 | 0.0127 |
| IGHV3/OR16-12 | 807 | 255 | 0.0001 | 0.9662 | 0.0226 | 0.0045 |
| IGHV1-38-4 | 508 | 253 | 0.0000 | 0.2882 | 0.0203 | 0.0075 |
| IGHV3-62 | 827 | 239 | 0.0001 | 0.6369 | 0.0145 | 0.0045 |
| IGHV5-78 | 744 | 228 | 0.0000 | 0.6486 | 0.0227 | 0.0051 |
| IGHV3-25 | 585 | 228 | 0.0000 | 0.7812 | 0.0273 | 0.0059 |
| IGHV3/OR16-15 | 575 | 220 | 0.0000 | 0.7874 | 0.0218 | 0.0029 |
| IGHV1/OR21-1 | 482 | 220 | 0.0000 | 0.3306 | 0.0169 | 0.0042 |
| IGHV3-16 | 661 | 212 | 0.0000 | 0.3247 | 0.0142 | 0.0022 |
| IGHV3-19 | 1,038 | 211 | 0.0000 | 0.6875 | 0.0358 | 0.0118 |
| IGHV1-68 | 609 | 186 | 0.0000 | 0.7519 | 0.0153 | 0.0023 |
| IGHV2-70D | 712 | 182 | 0.0000 | 0.7358 | 0.0562 | 0.0191 |
| IGHV3/OR15-7 | 427 | 181 | 0.0000 | 0.3430 | 0.0228 | 0.0039 |
| IGHV2-10 | 511 | 178 | 0.0000 | 0.4584 | 0.0144 | 0.0042 |
| IGHV3-54 | 273 | 147 | 0.0000 | 0.3356 | 0.0233 | 0.0041 |
| IGHV3-30-33 | 264 | 124 | 0.0000 | 1.3514 | 0.0466 | 0.0110 |
| IGHV2/OR16-5 | 247 | 117 | 0.0000 | 0.3356 | 0.0140 | 0.0024 |
| IGHV7-40 | 181 | 109 | 0.0000 | 0.1912 | 0.0180 | 0.0075 |
| IGHV3/OR16-6 | 336 | 105 | 0.0000 | 0.4196 | 0.0129 | 0.0012 |
| IGHV3/OR16-14 | 360 | 89 | 0.0000 | 2.2546 | 0.1064 | 0.0092 |
| IGHV3/OR16-16 | 321 | 75 | 0.0000 | 0.5464 | 0.0067 | 0.0007 |
| IGHV3-33-2 | 115 | 57 | 0.0000 | 0.4065 | 0.0258 | 0.0028 |
| IGHV3-30-2 | 81 | 52 | 0.0000 | 0.3401 | 0.0199 | 0.0032 |
| IGHV3-29 | 36 | 25 | 0.0000 | 0.0537 | 0.0058 | 0.0013 |
| IGHV3-32 | 17 | 17 | 0.0000 | 0.0228 | 0.0036 | 0.0017 |
| IGHV3-63 | 19 | 15 | 0.0000 | 0.0115 | 0.0031 | 0.0015 |
| IGHV3-30-22 | 17 | 12 | 0.0000 | 0.1164 | 0.0182 | 0.0048 |
| IGHV3-30-52 | 1 | 1 | 0.0046 | 0.0046 | 0.0046 | 0.0046 |

**Table S1c Overview of 34 D gene**

| **Gene** | **# Samples** | **# Donors** | **Min** | **Max** | **Mean** | **Median** |
| --- | --- | --- | --- | --- | --- | --- |
| IGHD7-27 | 2,145 | 582 | 0.1427 | 16.1131 | 1.7883 | 1.6184 |
| IGHD6-6 | 2,147 | 582 | 0.0923 | 12.7841 | 2.8384 | 2.8043 |
| IGHD6-19 | 2,150 | 582 | 0.2468 | 20.5128 | 7.2338 | 7.2503 |
| IGHD6-13 | 2,152 | 582 | 0.3279 | 25.0965 | 6.0661 | 5.8851 |
| IGHD5-24 | 2,150 | 582 | 0.1243 | 8.6614 | 2.4795 | 2.5097 |
| IGHD5-18 | 2,149 | 582 | 0.2188 | 14.9776 | 3.5332 | 3.4921 |
| IGHD5-12 | 2,147 | 582 | 0.1548 | 16.6243 | 2.6577 | 2.5598 |
| IGHD4-23 | 2,142 | 582 | 0.1645 | 9.0426 | 1.9784 | 1.9599 |
| IGHD4-17 | 2,152 | 582 | 0.3871 | 83.0572 | 4.3680 | 4.1428 |
| IGHD4-11 | 2,130 | 582 | 0.0378 | 9.3835 | 1.1772 | 1.1216 |
| IGHD3-9 | 2,147 | 582 | 0.1692 | 10.2362 | 2.6835 | 2.4368 |
| IGHD3-3 | 2,151 | 582 | 0.0829 | 19.6486 | 6.8638 | 6.4192 |
| IGHD3-22 | 2,152 | 582 | 0.4340 | 25.0000 | 7.7725 | 7.4232 |
| IGHD3-16 | 2,152 | 582 | 0.5525 | 12.9600 | 4.6024 | 4.4027 |
| IGHD3-10 | 2,152 | 582 | 0.7874 | 24.3373 | 10.3293 | 9.5225 |
| IGHD2-8 | 2,142 | 582 | 0.1661 | 9.2754 | 2.0994 | 2.0073 |
| IGHD2-2 | 2,152 | 582 | 0.1243 | 21.7796 | 6.5921 | 5.7111 |
| IGHD2-21 | 2,150 | 582 | 0.2273 | 26.0000 | 3.9219 | 3.6489 |
| IGHD2-15 | 2,152 | 582 | 0.0414 | 19.2652 | 5.7032 | 5.5524 |
| IGHD1-26 | 2,152 | 582 | 0.3846 | 33.8164 | 5.6820 | 5.8107 |
| IGHD1-1 | 2,145 | 582 | 0.0414 | 10.8696 | 2.0476 | 1.9254 |
| IGHD1-14 | 2,135 | 582 | 0.0468 | 10.5820 | 0.9764 | 0.8869 |
| IGHD6-25 | 2,129 | 581 | 0.0614 | 6.5217 | 0.7749 | 0.6862 |
| IGHD3/OR15-3a | 2,128 | 581 | 0.0387 | 9.4431 | 0.8246 | 0.7309 |
| IGHD2/OR15-2a | 2,144 | 581 | 0.0414 | 15.3911 | 1.2681 | 1.1674 |
| IGHD1-7 | 2,139 | 580 | 0.0675 | 7.4836 | 1.4450 | 1.3913 |
| IGHD5/OR15-5a | 2,076 | 576 | 0.0224 | 3.6036 | 0.4495 | 0.3800 |
| IGHD4/OR15-4a | 2,106 | 576 | 0.0206 | 5.4054 | 0.6489 | 0.5883 |
| IGHD1/OR15-1a | 2,096 | 575 | 0.0248 | 19.9509 | 0.4798 | 0.3854 |
| IGHD1-20 | 2,111 | 571 | 0.0345 | 7.5601 | 0.6657 | 0.6231 |
| IGHD2/OR15-2b | 1,871 | 493 | 0.0056 | 1.4706 | 0.1273 | 0.0985 |
| IGHD4-4 | 1,601 | 433 | 0.0035 | 1.9231 | 0.0652 | 0.0415 |
| IGHD1/OR15-1b | 1,527 | 422 | 0.0032 | 2.4048 | 0.0519 | 0.0308 |
| IGHD3/OR15-3b | 1,450 | 384 | 0.0029 | 1.2526 | 0.0441 | 0.0248 |

**Table S1d Overview of 6 J gene**

| **Gene** | **# Samples** | **# Donors** | **Min** | **Max** | **Mean** | **Median** |
| --- | --- | --- | --- | --- | --- | --- |
| IGHJ6 | 2,151 | 582 | 0.7962 | 44.0765 | 18.5514 | 18.3702 |
| IGHJ5 | 2,152 | 582 | 1.6563 | 50.3005 | 13.7042 | 13.0507 |
| IGHJ4 | 2,152 | 582 | 7.5362 | 76.2035 | 51.5722 | 50.5575 |
| IGHJ3 | 2,152 | 582 | 1.1013 | 77.5155 | 10.9532 | 10.8086 |
| IGHJ2 | 2,150 | 582 | 0.1969 | 16.1421 | 2.8489 | 2.8217 |
| IGHJ1 | 2,149 | 582 | 0.3150 | 10.7430 | 2.3848 | 2.1519 |

*Note:* V genes marked in gray are pseudogenes.
