## Supplemental Table 2 for "Large-scale Analysis of 2,152 dataset reveals key features of B cell biology and the antibody repertoire"

**Table S2. P values for genes which express significantly between the female and male**

| **Gene type** | **Gene** | **P_value** | **Female_median** | **Male_median** | **Diff (Female - Male)** |
| --- | --- | --- | --- | --- | --- |
| IGHV | IGHV4-30-2 | 8.76E-14 | 0.5436 | 0.3401 | 0.2035 |
|  | IGHV3-53 | 1.14E-12 | 2.7408 | 1.9597 | 0.7811 |
|  | IGHV3-20 | 1.22E-09 | 0.2895 | 0.4707 | -0.1813 |
|  | IGHV1-3 | 4.25E-09 | 1.8340 | 0.9583 | 0.8757 |
|  | IGHV7-4-1 | 2.65E-08 | 0.7788 | 0.1599 | 0.6189 |
|  | IGHV1-2 | 5.94E-07 | 2.6746 | 3.7814 | -1.1068 |
|  | IGHV3-64 | 8.28E-07 | 0.4171 | 0.5609 | -0.1438 |
|  | IGHV3-30-3 | 8.47E-06 | 1.9860 | 1.3209 | 0.6651 |
|  | IGHV4-38-2 | 9.28E-05 | 0.5203 | 1.0585 | -0.5382 |
|  | IGHV3-7 | 9.67E-05 | 5.1671 | 4.5268 | 0.6403 |
|  | IGHV3-48 | 0.0001 | 3.3171 | 3.6434 | -0.3263 |
|  | IGHV1-24 | 0.0001 | 0.9445 | 0.8228 | 0.1217 |
|  | IGHV3-43 | 0.0007 | 0.2507 | 0.5110 | -0.2603 |
|  | IGHV5-51 | 0.0020 | 2.1209 | 2.3228 | -0.2019 |
|  | IGHV3-11 | 0.0021 | 1.6091 | 2.1413 | -0.5322 |
|  | IGHV4-4 | 0.0035 | 2.3362 | 2.4885 | -0.1523 |
|  | IGHV3-71 | 0.0036 | 0.0390 | 0.0374 | 0.0016 |
|  | IGHV3-9 | 0.0041 | 1.9562 | 1.6179 | 0.3383 |
|  | IGHV3-15 | 0.0141 | 1.4615 | 1.8434 | -0.3819 |
|  | IGHV3-38 | 0.0160 | 0.0481 | 0.0372 | 0.0109 |
|  | IGHV1-18 | 0.0191 | 3.9917 | 3.8343 | 0.1574 |
|  | IGHV4-30-4 | 0.0202 | 0.4559 | 0.5763 | -0.1204 |
|  | IGHV3-21 | 0.0271 | 3.8069 | 4.4685 | -0.6616 |
|  | IGHV3-13 | 0.0321 | 0.3481 | 0.3648 | -0.0167 |
| IGHD | IGHD7-27 | 2.40E-05 | 1.6115 | 1.5493 | 0.0622 |
|  | IGHD2-2 | 2.65E-05 | 5.7559 | 5.8915 | -0.1356 |
|  | IGHD2-21 | 0.0011 | 4.0531 | 3.4292 | 0.6239 |
|  | IGHD5-18 | 0.0039 | 3.4277 | 3.7142 | -0.2864 |
|  | IGHD4-4 | 0.0072 | 0.0457 | 0.0380 | 0.0077 |
|  | IGHD3/OR15-3b | 0.0079 | 0.0281 | 0.0229 | 0.0052 |
|  | IGHD6-19 | 0.0159 | 7.5681 | 7.0375 | 0.5306 |
|  | IGHD3-3 | 0.0177 | 6.1224 | 4.9242 | 1.1982 |
|  | IGHD3-9 | 0.0420 | 2.8117 | 3.0149 | -0.2031 |
| IGHJ | IGHJ3 | 0.0001 | 12.0166 | 11.3415 | 0.6751 |
|  | IGHJ1 | 0.0005 | 2.4758 | 1.9029 | 0.5730 |
|  | IGHJ6 | 0.0007 | 16.4400 | 17.8449 | -1.4048 |
|  | IGHJ5 | 0.0042 | 12.6917 | 13.7119 | -1.0202 |
|  | IGHJ2 | 0.0136 | 2.6301 | 2.4469 | 0.1832 |

*Note:* A two-tailed unpaired samples t-test was done for each gene.
